# *KAMALA:* improved yield by finetuning cytokinin oxidase activity in rice

**DOI:** 10.64898/2026.01.23.701329

**Authors:** Manish Solanki, Faisal Yousuf, R.M. Sundaram, Sumalatha Katta, G.K. Srividya, Eswarayya Ramireddy, Sourav Chatterjee, Aashish Ranjan, Bipin Singh, P. Brajendra, C.N. Neeraja, S.V. Sai Prasad, Aravind K. Jukanti, Abdul Fiyaz Rahaman, Akshay S. Sakhare, Viswanathan Chinnusamy, Eliza P.I. Loo, Bing Yang, Wolf B. Frommer, Satendra K. Mangrauthia

## Abstract

Increasing crop yield is of major importance for food security worldwide. *Knock out* mutants in the rice cytokinin oxidase gene *CKX2* had shown potential for yield improvement. Here we explored whether the Indian mega variety Samba Mahsuri could be improved by introducing mutations into CKX2 by simultaneous editing of two internal sites predicted to subtly impact enzyme activity. *Knock out* and single in frame mutants generated by CRISPR/Cas12a caused substantial yield increases. Notably, among the 80 *CKX2* alleles evaluated, seven lines with different in-frame mutations in both sites produced yield increased up to 98.6% and early flowering. One line, *KAMALA*, showed superior agronomic performance in 16 of 18 diverse field locations (irrigated and rainfed ecologies) over three seasons in AICRPR trials (All India Coordinated Research Project on Rice), with an average 19% grain yield increase, early maturity, complete panicle emergence, and unaltered grain quality. Notably, in comparison to previously described *ckx2* mutants, the *KAMALA* allele is dominant, indicating unique properties of the variant. *KAMALA* was registered as the first genome-edited variety ready for cultivation by Indian farmers. 80 tons of *KAMALA* seeds are available for distribution. A Cas12a license for humanitarian distribution to small and marginal farmers is underway.

## Introduction

Rice is the most widely cultivated crop with an estimated acreage of 51 million hectares and grain production of 149 million tons during 2024-25 in India alone. Despite substantial progress in total rice grain production, yield gaps remain, and productivity requires further improvement (national average ∼2.9 tons/ha, www.global-agriculture.com/india-region/is-indias-paddy-production-on-a-sustainable-growth-path-5-year-trends-say-so/. Among the mega rice cultivars grown in India, Samba Mahsuri (BPT5204) occupies ∼10% of area cultivated. Samba Mahsuri is a fine grain variety popular among farmers and consumers due to premium cooking and eating quality^1^ despite having moderate grain yield (4-5 tons/ha), late maturity (145-150 days), incomplete panicle emergence, and weak culm. Efforts to improve yield and other traits had limited success due to loss of grain characteristics after introgression of trait enhancing genes^2^. Genome editing (GE) mitigates linkage-drag and allows site specific modifications for fine-tuning specific yield-related genes without altering the overall genetic makeup^3, 4^. CRISPR-Cas has successfully been utilized to edit multiple effector binding elements of three *SWEET* genes to produce broad spectrum disease resistance^5–8^. Cytokinin affects cell division, leaf development and senescence, shoot growth, and source-sink relationships^9^. Cytokinin oxidases/dehydrogenases (CKX) are key enzymes using flavin adenine dinucleotide (FAD) as a cofactor to metabolize cytokinins^10, 11^,. The rice genome contains eleven CKX paralogs. *CKX2* negatively impacts grain yield^12, 13^. Fourteen *CKX2* alleles were found to be associated with grain yield and spikelet number variation^14^, caused by variations in the promoter, 5’-untranslated region (5’-UTR), and coding region (Supplementary Figs. 1,2). The recessive *Grain number 1a* (*Gn1a*) allele of *CKX2*, carrying a 16 bp deletion in 5’-UTR and a 6 bp deletion in exon 1 (E1) of the Japanese cultivar Habataki, was associated with grain number/panicle trait. *Gn1a* had been utilized in breeding programs to increase grain yield^12^. Other naturally occurring *CKX2 knock out* alleles carry an 11bp E3-deletion (Chinese cultivar-5150), or a combination of a 6bp E1-deletion and an 11bp E3-deletion (Chinese cultivar-R498) characterized by higher spikelet number and stronger culm compared to *Gn1a*^13^. CRISPR-Cas editing was utilized to generate E1 frameshift mutations in Nipponbare^15^. *Knock out* lines showed a ∼44% higher grain number per panicle compared to the parent in greenhouse conditions. E1-frameshift mutants had similar grain number increases, however, reduced tiller number in field experiments^16^. Although loss-of-function of *CKX2* appear advantageous, such mutants were not enriched by breeding and selection. Based on recessiveness of *ckx2* mutants, we hypothesized that partial enzyme activity could have intermediate effects^12^. Guided by CKX structures, regions for finetuning cytokinin and cofactor binding were identified in exons 3 (E3) and 4 (E4) (Supplementary Fig 3) and targeted for Cpf1/Cas12a editing. An allelic series of 77 single/double indel lines was analyzed. Lines harboring premature stop codons exhibited improved performance. Surprisingly, in frame mutants in E3 or E4 showed increased yield but exhibited differential effects for tiller number and flowering time and in frame mutants with indels in E3/E4 performed superior compared to complete *knock outs* (Fig. 1). One line (*KAMALA;* DRR Dhan 100), was subjected to multi-location field trials for three seasons performed independently by AICRPR (All India Coordinated Research Project on Rice; nodal agency for conducting blind coded field evaluation of promising breeding lines prior to variety release). *KAMALA* exhibited an average 19% yield increase, 7 days earlier maturation, 92% higher culm strength, and complete panicle emergence. *KAMALA* was approved by the Varietal Identification Committee (VIC) as one of the first two genome-edited rice lines in India, alongside Pusa DST Rice1^17^. Notably, the *KAMALA* allele was dominant when introgressed into other elite rice varieties.

**Fig. 1.**
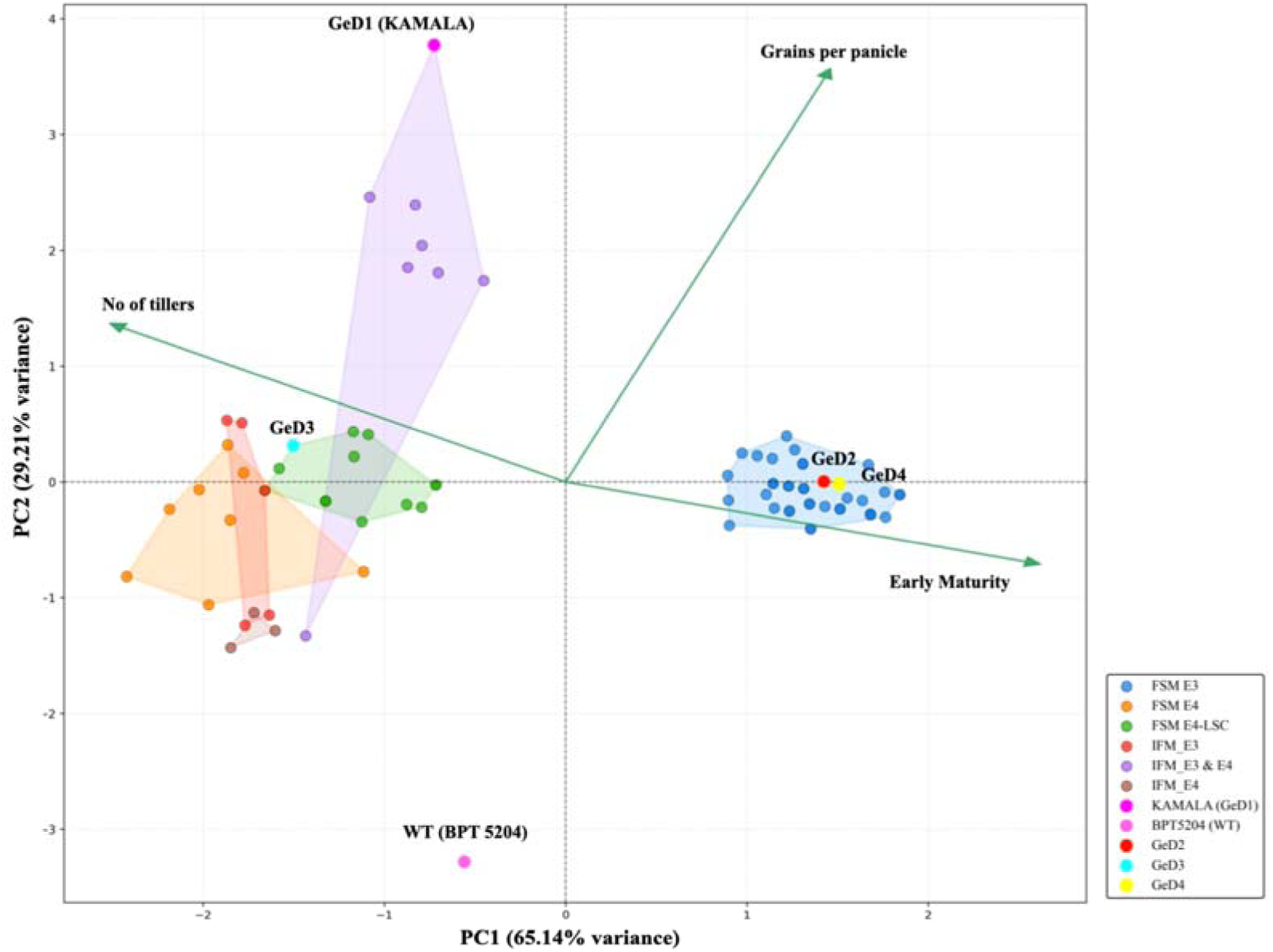
Principal Component Analysis (PCA) biplot of *CKX2* mutant lines for three traits: grains per panicle, number of tillers, and days to maturity. PC1 (66.44% variance) shows a trade-off between grain productivity per panicle and tillering capacity: negative values (left) indicate early maturity with more tillers; positive values (right) indicate later maturity with fewer tillers. PC2 (28.68% variance) represents grain productivity: higher values (top) indicate more grains per panicle; lower values (bottom) indicate fewer grains per panicle. Each point represents one GeD line. Green arrows (loading vectors) show trait contributions; direction indicates where high-trait genotypes are positioned, and length is proportional to contribution strength. Quadrants: upper-left (early maturity + high grain productivity), upper-right (later maturity + high grain productivity), lower-left (early maturity + lower grain productivity), lower-right (less desirable combinations). The GeD1 (*KAMALA*) is in the upper-left quadrant with the highest grains per panicle (480), high tillering (10 tillers), and early maturity (130 days). The details of mutations and phenotype of GeD lines are given in Supplementary Table 2. *GeD18 with deletion of only one amino acid at both the edited sites.

## Results

### Effects of in-frame mutations close to cytokinin and FAD-binding domains of CKX2

Analysis of natural variation for *CKX2* in 4,726 rice accessions (RiceVarMap) implicated 7 missense variants and 2 INDELs (insertion/deletion polymorphisms) in E1; 3 missense variants and one INDEL in E2; two missense variants and two INDELs in E3; and two missense variants in E4. Only eight accessions showed frameshift mutations causing protein disruption (Supplementary Fig. 2, Supplementary Table 1). While loss-of-function alleles showed higher grains/panicle, retention of functional CKX2 appears important and inactivation may have phenotypic trade-offs. We thus aimed at evaluating *CKX2* variants that retained function and only partially impacted CKX activity to evaluate a potential linear inverse correlation between CKX activity and improved characteristics. To generate a series of *ckx2* variants, we targeted E3 and E4 by CRISPR-Cas12a multiplex editing (Supplementary Fig. 4). We evaluated 80 variants carrying distinct edits in Samba Mahsuri (‘wildtype ‘(WT); Supplementary Fig. 5). Initially, lines were evaluated in greenhouse conditions for grain, panicle and productive tiller number and time-to-maturity (Supplementary Fig. 6). Principal Component Analysis (PCA) was used to evaluate correlation of mutation type and performance (Fig. 1, Supplementary Tables 2,3). Three GE lines (GeD7, 14, 52) were sterile (likely due to other mutations or conditions). PCA of key features of the 78 rice genotypes explained 94.4% of total variance. PC1 represented trade-off between grain number/panicle and tiller number, with negative values indicating early maturity with more tillers, and positive values indicating late maturity with fewer tillers. PC2 represented grain productivity. The PCA biplot revealed that lines carrying specific types of mutations grouped together. Surprisingly however, lines with premature stop codons and internal deletions in E3 and E4 separated for tiller number, maturity and grain yield. While mutants carrying premature stop codons showed comparable yield increases, E3 loss of function mutants behaved different from those in E4 regarding flowering time and tillering. Single in frame mutants in E3 or E4 behaved similar to those carrying premature stop in E4 with respect to tillering and flowering. Notably, mutants (with one exception; GeD18, carrying single amino acid deletions in E3 and E4) grouped together having substantially higher yield compared to those with premature stop codons or in frame mutations at one site (Fig. 1). A common feature of this group #1 was in-frame mutations in both E3 and E4. Group 1 lines showed much-improved agronomic profiles which broke the typical trade-offs between the traits. GeD1 (hereafter referred to as *KAMALA*) was positioned in the upper-left quadrant (low PC1, high PC2) and exhibited exceptional performance, with 480 grains per panicle (the highest in the dataset), 10 tillers, and early maturity at 130 days. In contrast, the WT was positioned in the lower-left quadrant, with lower grain productivity (170 grains per panicle) and later maturity (145 days) (Fig. 1).

### *CKX2* edited lines exhibited early flowering, complete panicle emergence, and high grain yield

Among the edits generated here (Fig. 1, Supplementary Fig. 5, Supplementary Table 2), four homozygous mutants carrying different mutations in E3 and/or E4, namely *KAMALA*, GeD2, GeD3, and GeD4, were chosen for a more detailed characterization (Supplementary Fig. 7-11). Overlap PCR with 39 primer sets (<500 bp each, >50 bp overlap) for the plasmid used for transformation did not detect transgene fragments, and all four lines were sensitive to hygromycin (Supplementary Figs. 12,13). *KAMALA* carries *in-frame* mutations in both exons (group 1), while GeD2 and GeD4 had frameshifts in E3 (group 4), and GeD3 an *in-frame* mutation in E3 and a frameshift in E4 (group 6). Morphological and grain yield attributes were superior in all four edited lines, with markedly enhanced vegetative vigor and early flowering relative to WT (Fig. 2, Supplementary Table 4). Improvements in panicle architecture were evident, with increased branching and spikelet density. Samba Mahsuri is known to frequently show incomplete panicle emergence, thereby limiting reproductive efficiency. Notably, all four GeD lines exhibited complete panicle exertion. Although grain appearance was similar to WT (Fig. 2d), grain length and width were noticeable reduced in GeD2, GeD3, and GeD4. Notably, the number of days to 50% flowering was reduced; combined with significant increases in grain yield per plant in all four mutants (Fig. 2). Increased grain yield per plant was primarily driven by higher numbers of grains/panicle, which corresponded to increased panicle length, increased number of primary branches/panicles, and increased panicle branch length. The average grain number per panicle increased from 242 in WT to 402 (*KAMALA*), 386 (GeD2), 327 (GeD3), and 356 (GeD4).

**Fig. 2.**
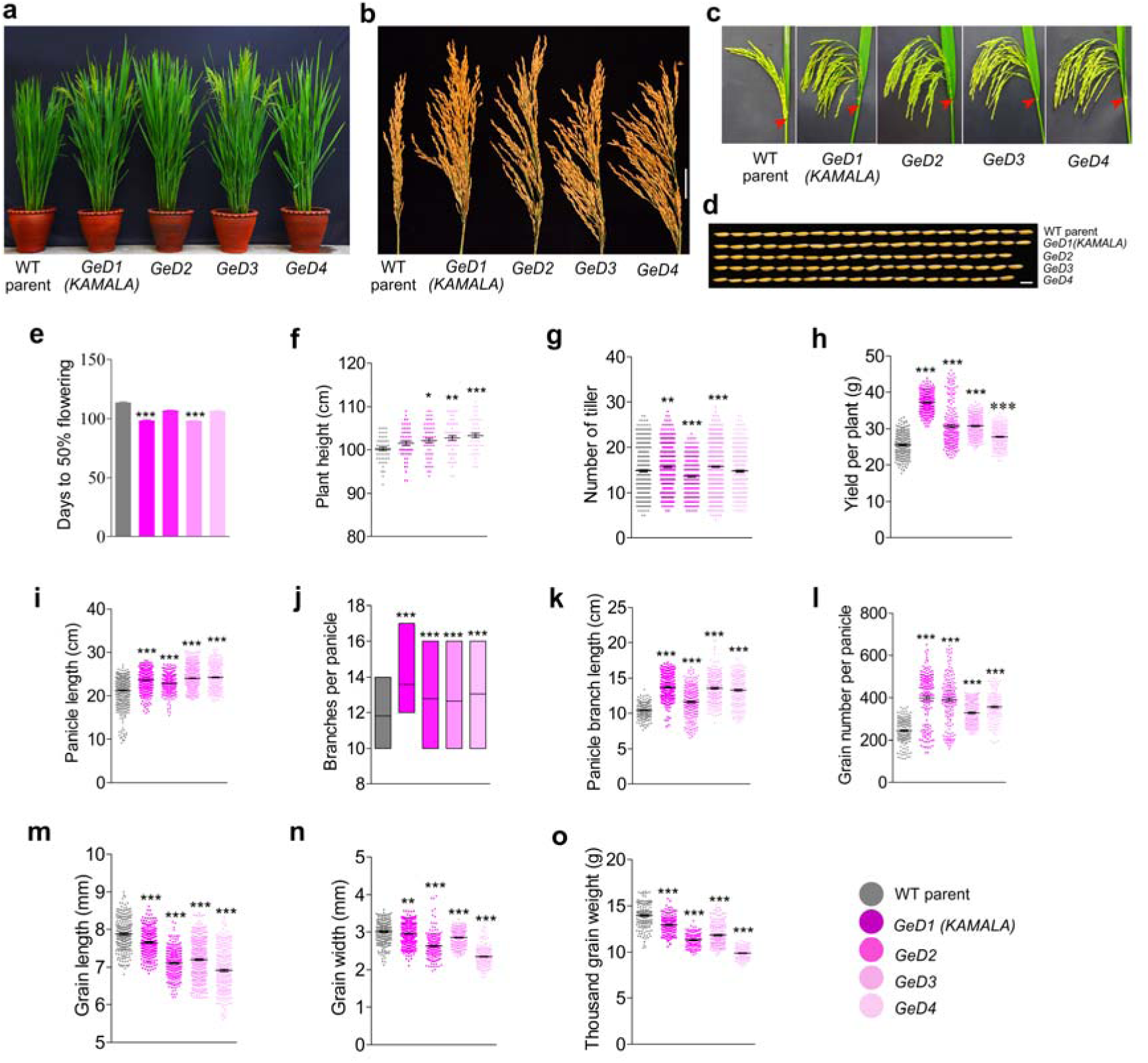
Targeted mutagenesis of *OsCKX2* enhances panicle architecture and grain yield in rice mega-variety Samba Mahsuri (WT parent). **a,** gross morphology of WT and four genome-edited lines (GeD1 to GeD4) at 100 days post-sowing. **b-d,** reproductive organ phenotypes: (**b**) mature panicle morphology (scale bar = 5 cm), (**c**) panicle exertion, and (**d**) grain size (scale bar = 5 mm). **e,** early flowering (5-20 days) in edited lines across dry and wet seasons. **f-g,** other traits impacted: (**f**) plant height and (**g**) tiller number per plant. h, grain yield per plant. **i-l,** improved panicle architecture: (**i**) panicle length, (**j**) primary branch number, (**k**) branch length, (**l**) grain number per primary panicle. **m-o,** grain characteristics: (**m**) grain length, (**n**) width, (**o**) thousand-grain weight. The raw data of these parameters is given in Supplementary Table 4. All plots show mean ± s.e.m.; sample sizes are given in methods. Statistical significance: one-way ANOVA with Tukey’s HSD. Asterisks (*) indicate significance levels: * = *P* value < 0.05; ** = *P* value < 0.001 and *** = *P* value < 0.0001.

### Regulatory exemption and environmental release of *KAMALA*

Based on superior performance, *KAMALA* was prepared for environmental release per Standard Operating Procedures (SOP) published by the Department of Biotechnology (DBT), Government of India (https://dbtindia.gov.in/sites/default/files/SOPs%20on%20Genome%20Edited%20Plants_0.pdf). A detailed dossier was submitted providing evidence for outcrossing of foreign DNA and homozygosity of edits, using overlap PCR, hygromycin sensitivity, and sequence of mutated in T2 and T3 (Supplementary Figs. 12,13). Transgene fragments were not detected in *KAMALA* using protocols prescribed by DBT (Section 5, pp.12-13, SOP). Recent advances offer more reliable approaches for evaluating transgene presence^6, 8^. Using PacBio HiFi whole-genome sequencing (WGS), reads from axenically grown WT and *KAMALA* that met our sequencing standards^8^ (Supplementary Fig. 14a) were mapped against the vector (Supplementary Fig. 14b). We did not detect reads for vector sequences, except the endogenous rice ubiquitin promoter in *KAMALA* and WT (563 reads, mean overlap 470 bp; 612 reads, mean overlap 473 bp). Regions upstream of rUbiP showed marginal depth overlap (mean read length 1-2 bp) to vector sequences in WT and *KAMALA*; likely read-trimming artifacts. Thus, vector sequences had apparently been eliminated by outcrossing. In May 2023, regulatory agencies granted exemption, making *KAMALA* the first genome-edited crop approved for environmental release in India (https://www.pib.gov.in/PressReleasePage.aspx?PRID=2126802&reg=3&lang=2). The exemption enabled field trials in Telangana, and in the countrywide AICRPR program (Stepwise procedures described in Supplementary Note 1).

### Two-year field evaluation of *KAMALA* in Telangana

Improved greenhouse performance often does not hold up under field conditions. To evaluate field performance, *KAMALA* was cultivated in Southern India (semi-arid) at the ICAR-IIRR research fields in Hyderabad, Telangana in 2023 and 2024 (Fig. 3a). *KAMALA* exhibited markedly enhanced seedling vigor, thicker and more highly branched roots (Fig.3b,c,f), broader leaves (Fig. 3d, g), and thicker culms (2x thicker culm wall; Fig. 3e,h). Root architecture (diameter, forks, tips) was superior for *KAMALA* (Supplementary Fig. 15). Vigor index-I and II, chlorophyll, and carotenoid content of *KAMALA* were significantly higher compared to WT (Supplementary Fig. 15). At the reproductive (booting) stage, *KAMALA* maintained its superior phenotype (greater growth vigor, thicker culms, increased root volume relative to WT; Supplementary Fig. 16). Notably, grain yield increased 41-52% and the Harvest Index (HI) was improved in *KAMALA* across three seasons (Fig. 3l,k, 47% vs. 43% in WT). Culm strength at maturity was significantly higher (0.73 kg cm ¹ in *KAMALA* vs. 0.38 kg cm ¹ in WT) likely reducing lodging (Fig. 3i). The root:shoot ratio of length, fresh and dry weight indicated desirable agronomic features of *KAMALA* (Supplementary Fig. 16). The flag leaf, which plays critical roles in grain filling, showed increased length, width, enhanced photosynthesis rate and stomatal conductance in *KAMALA* (Fig. 3j, Supplementary Fig. 16). Photosynthesis rates were 22.2% higher in *KAMALA* (Fig. 3j), consistent with the increase of steady state mRNA levels for key chlorophyll biosynthesis and photosynthesis-related genes in flag leaves (Supplementary Fig. 17).

**Fig. 3.**
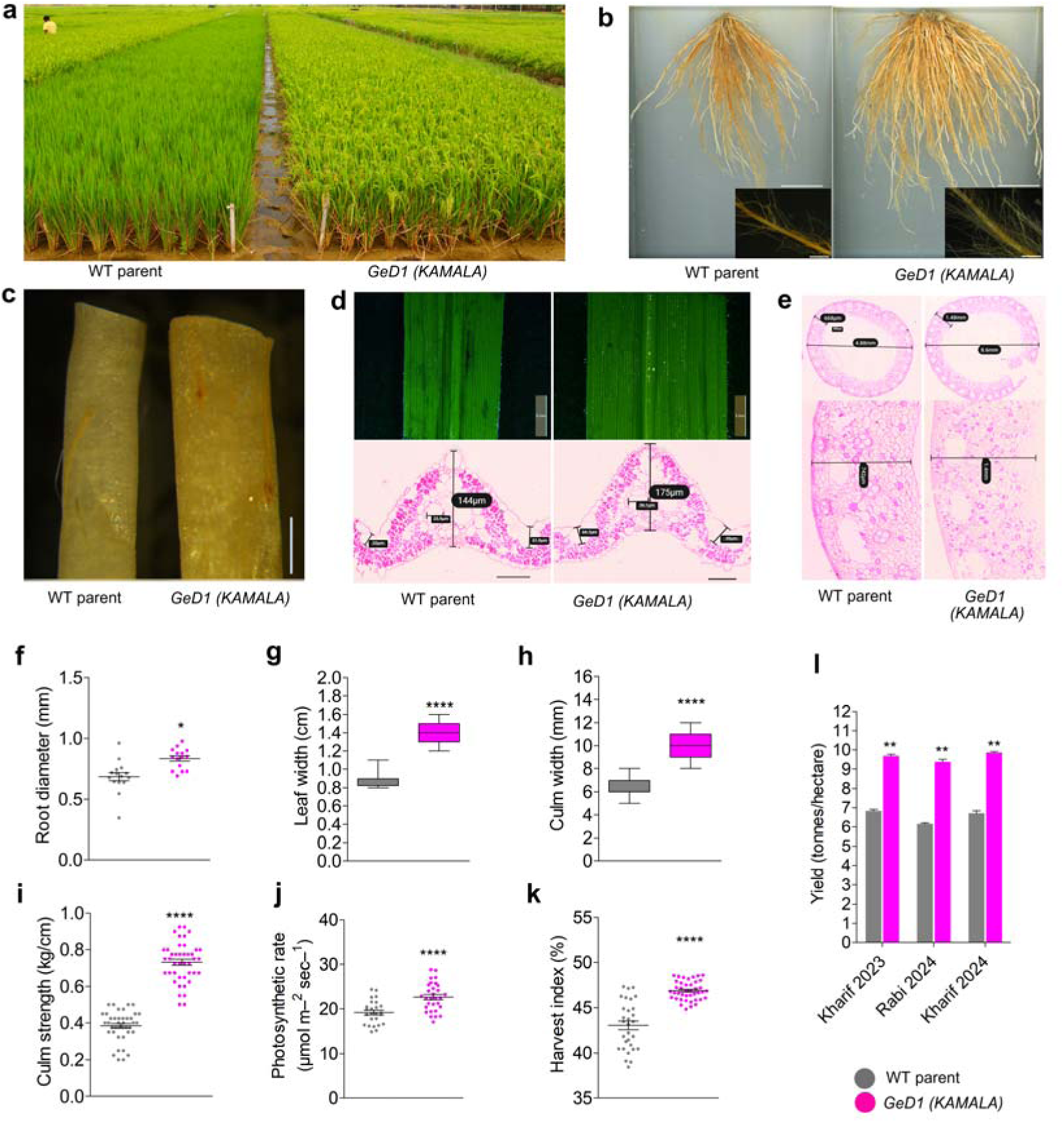
GeD1 (*KAMALA)* exhibits superior growth and enhanced grain yield compared to Samba Mahsuri (WT parent) under field conditions. **a**, field trial at ICAR–Indian Institute of Rice Research (IIRR), Hyderabad, shows robust growth of *KAMALA* in comparison to WT. **b-h**, analysis of agronomic and physiological traits at vegetative growth stage: **b**, seedling root morphology (scale bar, 4 cm) and root hair density insets. **c**, comparison of root thickness (scale bar, 0.5 mm). **d**, leaf surface view (upper) and transverse sections (lower) show increased thickness in *KAMALA* (175 μm) compared to WT (144 μm) (scale bar, 100 μm). **e**, culm transverse sections demonstrate vascular bundle distribution and diameter. **f**, root diameter (mm); **g**, leaf width (cm); **h**, culm width (mm); **i-l**, quantitative analysis of agronomic and physiological traits at reproductive growth stage: **i**, culm strength (kg/cm); **j**, photosynthetic rate (μmol m ²s ¹); **k**, Harvest Index (%); **l,** grain yield (tonnes per hectare) over three cropping seasons (Kharif/wet 2023, Rabi/dry 2024, Kharif 2024). Data are mean ± s.e.m. (n in Methods). *KAMALA* (pink) and WT parent (grey) were compared with one-way ANOVA and Tukey’s HSD test. Asterisks (*) indicate significance levels: * = *P* value < 0.05; ** = *P* value < 0.001 and **** = *P* value < 0.00001.

### Superior panicle architecture while retaining original grain quality traits in *KAMALA*

Grain number harvested from a single panicle of *KAMALA* ranged from 491±51 (primary panicle) to 195±46 (tertiary or last panicle); WT grain number ranged from 292±37 to 156±61 (Fig. 4a, Supplementary Table 5). Grain morphology (with/without husk) was similar to WT (Fig. 4b,c). Average dry weight of primary panicles was 5.8 g for *KAMALA* and 3.5 g for WT (Fig. 4d). Notably, the proportion of unfilled grains per panicle did not differ significantly (Fig. 4e). Importantly, critical grain quality and milling parameters (head rice recovery, hulling and milling percentage, amylose content, gel consistency) were similar to WT (Fig. 4f-j). Given the remarkable improvements in panicle architecture and yield traits observed in *KAMALA*, we investigated mRNA levels of known yield-associated genes (Supplementary Table 6). Transcript profiling of inflorescence primordia revealed increased mRNA levels for several genes in *KAMALA* that play important roles in determining different yield components, i.e. *OsTGW3, OsNAL1, OsNOG1, OsSPL14*, *OsGSN1*, and *OsEP3* (Fig. 4k, Supplementary Fig. 18). *OsTGW3* and *OsGSN1* impact grain size; *OsNOG1* and *OsSPL14* affect panicle branching; and *OsNAL1* and *OsEP3* play roles in photosynthesis and plant architecture^18–20^. RNA-seq analysis of inflorescence tissues identified genes that may contribute to the phenotypic differences observed in *KAMALA* relative to WT (Supplementary Note 2, Supplementary Figure 19, Supplementary Tables 7, 8). *OsANK1* has been identified as a key regulator of panicle branching and spikelet number in indica rice^21^. Correspondingly, *KAMALA* exhibited increased panicle length, more branches per panicle, and extended branch length (Fig. 4 k,l).

**Fig. 4.**
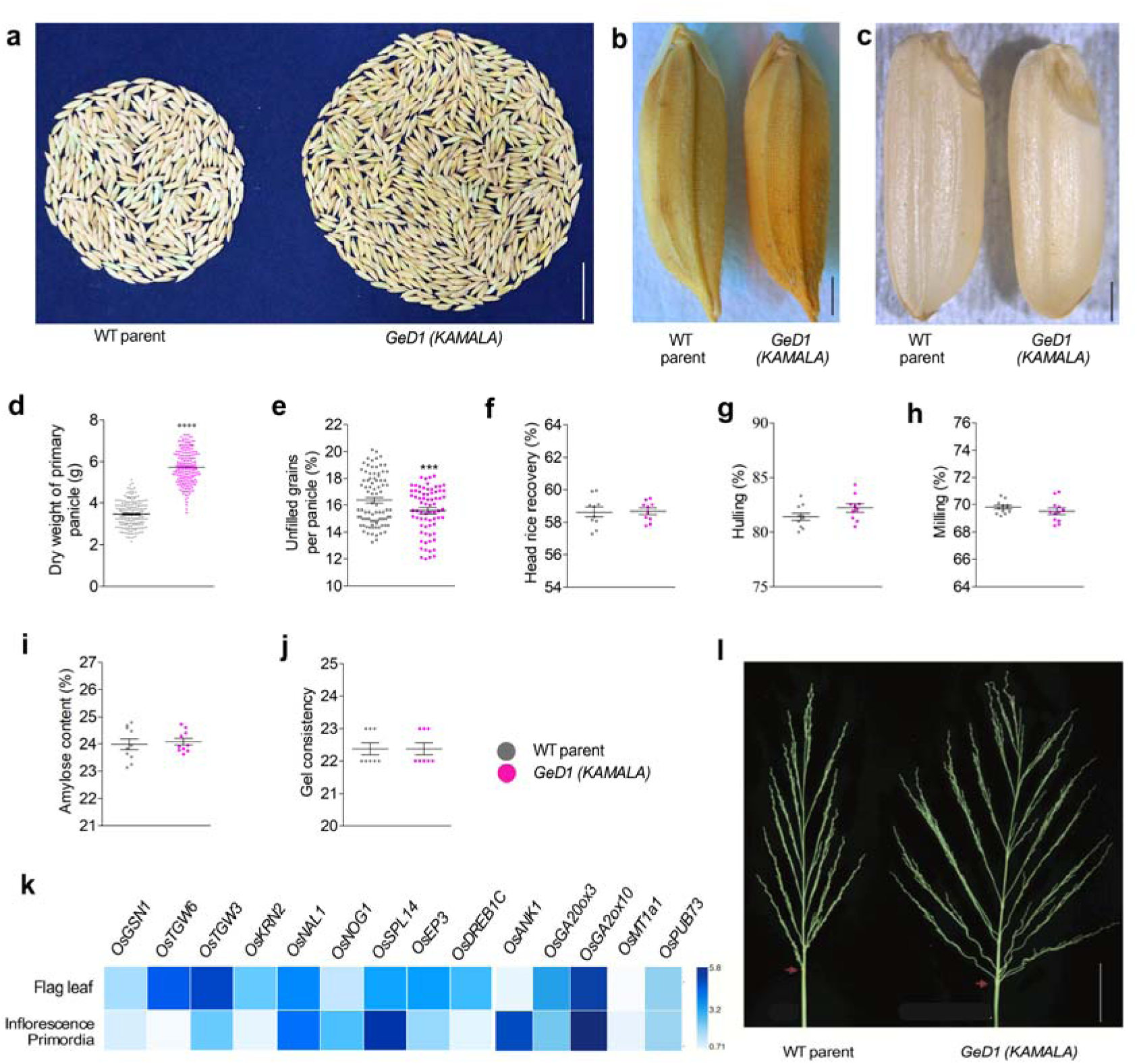
GeD1 (*KAMALA)* produces higher grains per panicle without compromising quality traits. **a,** comparison of grains harvested from a single panicle of Samba Mahsuri (WT parent) and *KAMALA* (scale bar = 1.5 cm). **b-c,** grain morphology shown with husk (**b**) and without husk (**c**) (scale bar = 1 mm). Quantitative analysis of grain filling and quality parameters, **d**, dry weight of primary panicle, **e**, percentage of unfilled grains, **f**, head rice recovery, **g**, hulling, **h**, milling, **i**, amylose content, **j**, gel consistency. **k**, relative expression of genes in the flag leaf and inflorescence tissues of *KAMALA*, referenced against WT parent. RT-qPCR data supporting this analysis are provided in Supplementary Fig. 17. **l,** compares panicle architecture, highlighting enhanced branching in *GeD1* (*KAMALA)*, which mirrors *OsANK1*-overexpression phenotype. Data are presented as mean ± s.e.m., *n* = as specified in the Methods section for each trait. Statistical significance between *KAMALA* and WT parent was evaluated using one-way ANOVA followed by Tukey’s Honestly Significant Difference (HSD) test. (*P* < 0.05). Asterisks (*) indicate significance levels: ** = *P* value < 0.001 and **** = *P* value < 0.00001.

### Early maturity of *KAMALA* in field conditions

Notably, *KAMALA* achieved the yield increase in a shorter maturation time relative to WT, due to early flowering in both greenhouse and diverse field conditions (Figs. 2a,e; 3a; Supplementary Fig. 20a). In Hyderabad field conditions, flowering initiated 20-25 days earlier compared to WT (Supplementary Fig. 20). Although maturation varied in the different AICRPR testing locations (Supplementary Table 9), maturity occurred on average seven days earlier relative to WT. Eight weeks after sowing, the *KAMALA* shoot apices transitioned to reproduction, while most WT apices were still in the vegetative phase (Supplementary Fig. 20b). Candidate genes were chosen for RT-qPCR to identify gene networks involved in flower time regulation (Supplementary Fig. 21). Relative mRNA levels of genes with roles in heading date/flowering differed in WT and *KAMALA*. The mRNA levels of negative factors influencing heading date (*OsPINE1, OsHD2, OsGI, OsHDR1, OsRCN2, OsETR2, OsHd1*, *OsDTH8*) were significantly lower in *KAMALA* (Supplementary Fig. 20c). Conversely, mRNA levels of factors positively affecting heading date (*OsFTIP1, OsMADS18, OsMADS51, OsSDG725, OsZHD2, OsAP1, OsELF3.2*, fold changes 17, 12, 7, 10, 9, 6, 3, respectively) were higher in *KAMALA* (Supplementary Fig. 20d). In summary, *KAMALA* exhibited accelerated reproductive transition, underpinned by altered mRNA levels of flowering time genes and enhanced meristematic activity, producing more yield faster relative to WT.

### Multi-location field trials of *KAMALA* and identification as a farmer-suitable variety

Based its performance, *KAMALA* was nominated for evaluation under the AICRPR field trials. Independent AICRPR trials, responsible for evaluating >1200 breeding lines each year in multi-location field trials (here wet season 2023, dry season 2024, wet season 2024) across 10 major rice-growing states including diverse agroclimatic zones, demonstrated *KAMALA’s* broad adaptability and yield advantage (Fig. 5a). Yield gains ranged from 4.7% (Kerala) to 99% (Madhya Pradesh) relative to WT. We note that *KAMALA* showed inferior performance in two out of the eighteen locations, namely Nellore (Andhra Pradesh) and Pattambi (Kerala) with yields 11/7% below WT, respectively. On average, *KAMALA* exhibited a ∼19% yield advantage over WT (Supplementary Table 9). The Varietal Identification Committee (VIC) approved *KAMALA* (DRR Dhan 100) as India’s first genome-edited plant variety for cultivation. To evaluate the effect of the *KAMALA* allele in other elite lines, PCR (Fig. 5b) and CAPS (Cleaved Amplified Polymorphic Sequences) markers (Fig. 5c; Supplementary Fig. 22) were developed and used for introgresssion into popular varieties (DRR Dhan 48, DRR Dhan 53, DRR Dhan 60, DRR Dhan 62, Improved Samba Mahsuri, and Vardhan; Supplementary Fig. 22). The offspring of these lines showed superior characters for plant vigor, growth, and grain yield (Fig. 5d-g).

**Fig. 5.**
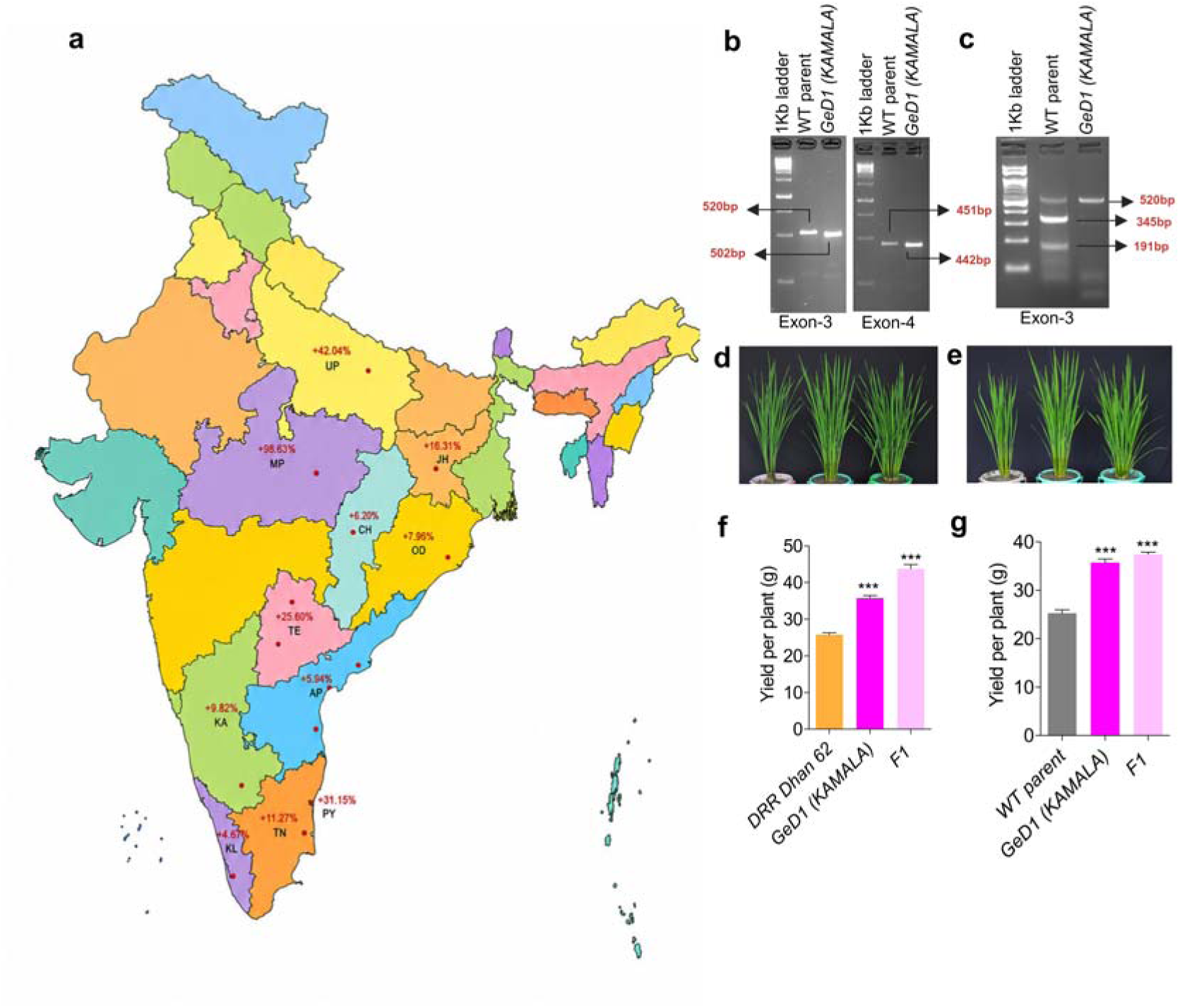
Multi-location and multi-season field evaluation of GeD1 *(KAMALA)* and development of allele-specific DNA markers for trait introgression. **a,** Map of India showing the percentage yield increase of *KAMALA* over Samba Mahsuri (WT parent) across different rice-growing states, based on AICRPR Progress Reports (2023 & 2024). Red dots indicate the specific locations where field trials were conducted in the respective states (see Supplementary Table 9 for details). The locations were mapped using geographic coordinates in R. The map was coloured using ChatGPT. **b,** development and validation of PCR based DNA marker specific to *OsCKX2* allele in *KAMALA*. **c,** CAPS (Cleaved Amplified Polymorphic Sequence) DNA marker specific to *OsCKX2* allele in *KAMALA*. PCR products resolved on 2.5% agarose gels with a 1 kb ladder. **d-g,** introgression of *KAMALA CKX2* allele into DRR Dhan 62 and WT parent using allele specific DNA markers developed in this study. **d,** morphology images sequence: DRR Dhan 62, *KAMALA*, F1 plant. **e,** morphology images sequence: WT parent, *KAMALA*, F1 plant. f, comparative yield per plant for DRR Dhan 62, *KAMALA*, and F1 progenies. g, comparative yield per plant for WT parent, *KAMALA*, and F1 progenies. Data are mean ± s.e.m. (n = 20). One-way ANOVA and Tukey’s HSD test. Asterisk (*) indicate significance levels: *** = *P* value < 0.0001.

### Dominance of the *KAMALA* mutation

The parental Samba Mahsuri *Gn1a* allele of *CKX2* is characterized by a 16 bp 5′-UTR deletion, a 2 aa deletion plus two aa substitutions in E1 appeared semi-dominant with grain yield highest in homozygous lines and intermediate in heterozygotes (Supplementary Fig. 1)^12^. In-frame deletion mutants would then be expected to show an intermediate phenotype and be semi-dominant. Surprisingly, F plants of crosses between *KAMALA* and Samba Mahsuri or DRR Dhan 62 showed dominance of the *KAMALA* allele (Fig. 5d,e). Given the potential influence of heterosis and hybridization-induced epigenetic reprogramming, including trans-chromosomal methylation in F1 generation^22^, BC F populations of crosses of *KAMALA* with Improved Samba Mahsuri and DRR Dhan 48 showed no significant phenotypic differences between homo- and heterozygous plants, supporting dominance (Supplementary Figs. 23,24). We compared structural flexibility for CKX2 of ancestral alleles (not carrying CKX2 variants present in Samba Mahsuri), Gn1a (Samba Mahsuri), and *KAMALA* using CABS-Flex3^23^. The *KAMALA* flexibility profile differed from Gn1a and was seemingly restored ancestral CKX2 (Supplementary Figs. 25,26). Deletion of residues 376-383 shortens the C-terminal α-helix from aa364-378 in Samba Mahsuri to 364-370 in *KAMALA*, whereas insertion at aa384-387 leads to an extension of neighboring loops. Deletion of aa512-514 lead to a shortened loop (Fig. 6a). Despite these deletions and the insertion, *KAMALA* did not exhibit substantial deviations in global flexibility compared to the ancestral CKX2 (Supplementary Fig. 25). *KAMALA* was predicted to maintain a flexibility profile similar to ancestral CKX2, while exhibiting reduced cavity volume, implying reduced enzymatic activity alongside preservation of core protein function (Fig. 6b, Supplementary Fig. 27).

**Fig. 6.**
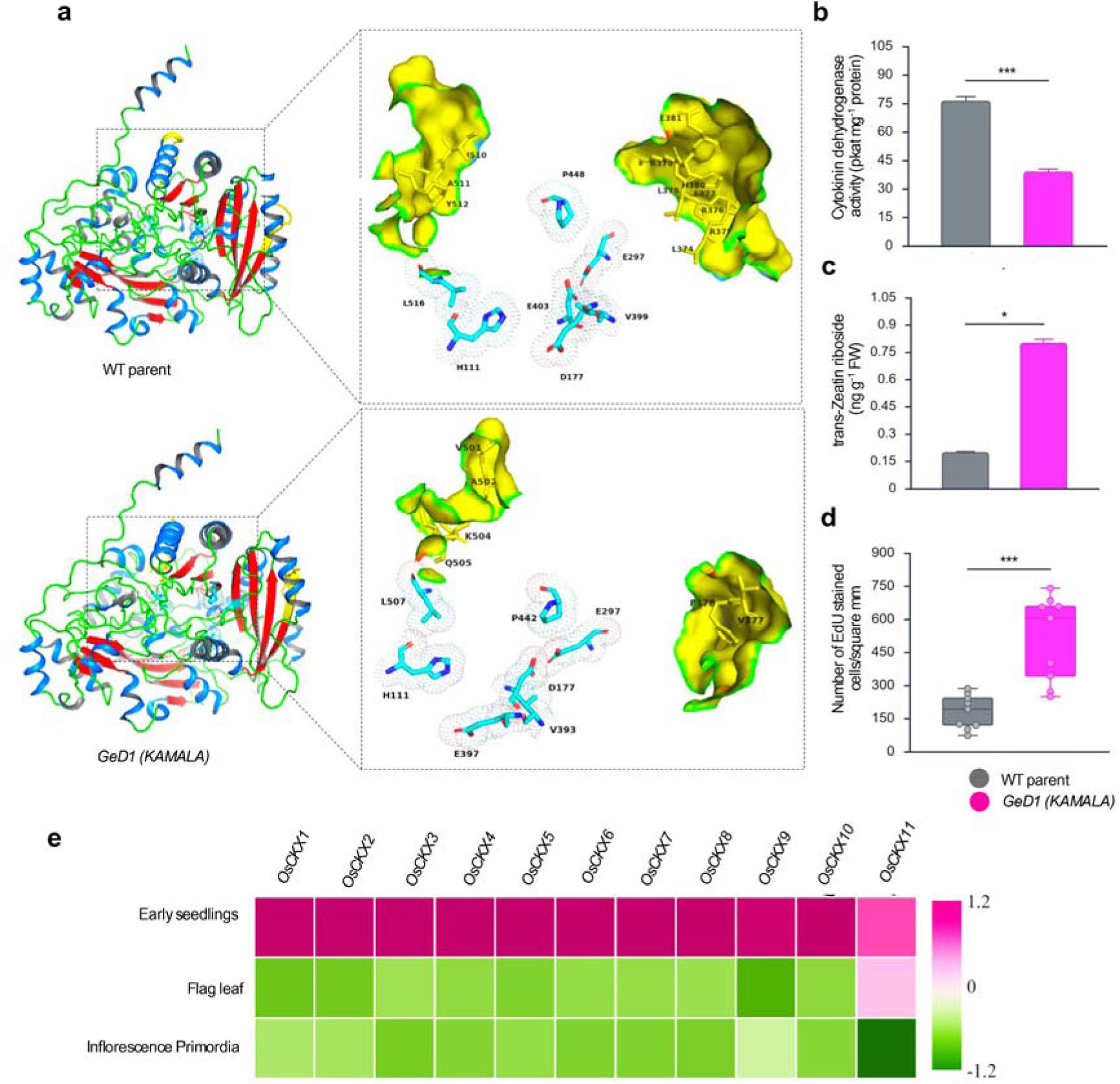
Fine-tuning of cytokinin metabolism drives enhanced performance of GeD1 (*KAMALA)*. **a**, AlphaFold predicted three-dimensional structural models of CKX2 protein of Samba Mahsuri (WT parent) and *KAMALA*. Left panels: Secondary structure elements are depicted with colour-coding (α-helices, marine; β-strands, red; loops, green). Right panels: Residues critical for catalytic activity are rendered in cyan stick representation. Regions harbouring deletions or insertions in *KAMALA* relative to WT-CKX2 are highlighted in yellow surface representation. Compared to Samba Mahsuri, the *KAMALA* variant exhibits a reduced cavity volume suggesting impaired substrate interaction. **b**, reduced substrate affinity corresponds to decreased cytokinin dehydrogenase activity. **c,** increased trans-zeatin riboside content in *KAMALA* inflorescence tissue. **d**, quantification of EdU-stained actively dividing cells. **e,** heat map representing expression profiles of *OsCKX1-OsCKX11* in different tissues and growth stages. RT-qPCR data supporting this analysis are provided in Supplementary Fig. 27. Results are shown as averages with standard error, with three separate samples for each test. Statistics used one-way ANOVA with Tukey’s HSD test. Asterisks (*) indicate significance levels: ** = *P* value < 0.001 and **** = *P* value < 0.00001.

To gain insights into the superior agronomic performance of *KAMALA*, structural models of the CKX2 proteins were analyzed, and overall, *in-planta* enzyme activity and cytokinin levels were measured in inflorescences (development stage 5cm length; Supplementary Fig. 28). AlphaFold3 predictions revealed unstructured regions in GeD2, GeD3, and GeD4, whereas *KAMALA* maintained a well-defined structure, but contains an additional cavity (Fig. 6a; Supplementary Fig. 28a). All four lines had reduced CKX activity in inflorescence extracts, with *KAMALA* having intermediate activity. *trans*-zeatin riboside levels correlated with enzyme activity. Thus, in contrast to what was hypothesized initially, intermediate activity did not lead to reduced performance, indicating that reducing activity to intermediate levels might be advantageous over loss (Figs. 1,2).

### Increased cytokinin levels in *KAMALA*

Evaluation of the effect of CKX2 mutants on *in-planta* cytokinin oxidase/dehydrogenase activity is challenging due to the presence of 11 CKX paralogs in rice^16^. A reduced overall CKX activity could lead to reduced degradation resulting in increased cytokinin levels. Although only *CKX2* was edited, *KAMALA* showed elevated cytokinin levels (Fig. 6c: Supplementary Fig. 28c), and higher cell numbers entering the cell cycle in the shoot apical meristem (SAM; Fig. 6d). This effect is likely complex, since relative mRNA levels of *CKX2* as well as ten other CKX family members was decreased in inflorescence primordia and flag leaf (except *OsCKX11*; Fig. 6e; Supplementary Fig. 29).

## Discussion

Successful development of genome-edited crops for large-scale cultivation remains rare, e.g. GABA-enriched tomatoes in Japan^24^. Edited waxy corn and high oleic acid soybean advanced beyond greenhouse tests, while the majority of genome-edited rice research has lagged behind in field translation^25^. Among the genome edited crops, rice accounts for the highest number of documented studies (2076 research papers), underscoring its prominence in genome editing research^26^. Previous efforts focused on easier transformable *japonica* varieties (*Kitaake*, *Nipponbare*, and *Taipei 309)*, but are not cultivated in India or much of Southeast Asia^27–29^. The majority of rice cultivated in India belongs to the *ssp. indica*, which is recalcitrant to tissue culture-based transformation including Samba Mahsuri^30, 31^,.

Climate change impact, shrinking farm resources and low rice productivity is a foremost concern for food security in India. Low productivity is mainly ascribed to a lack/limited presence of superior alleles of key yield-related genes within indigenous rice germplasm. Here, targeted genome editing of two sites close to important residues involved in cytokinin and FAD-binding in CKX2 was done in Samba Mahsuri. Surprisingly, combined in-frame mutations in E3/E4 yielded superior phenotypes, being markedly better relative to lines carrying frameshift mutations or in-frame edits at single sites, although all mutants were superior over WT. Surprisingly, lines with premature stop codons in E3 behaved differently with respect to tillering and maturation to those with premature stop codons in E4. Earlier efforts to target *CKX2* through CRISPR/Cas genome editing focused either E1 or E2 or both, and most of these edits resulted into frameshift mutations^13, 15, 16, 32, 33^.

Analyses of the predicted structure indicated that the CKX2-fold may not be impacted to a large degree, rather more subtle changes like the orientation of α-helix (364-370) and binding pocket appeared different. Structural predictions combined with Boltz-2 analyses supported subtle activity changes. Direct structural and functional analyses will help evaluating the predictions. Notably, *KAMALA* behaved dominant. Whether dominance is due to unidentified roles of the full-length CKX2, e.g. as an interaction partner for other proteins, effects due to potential mistargeting, or whether the binding properties of the mutant form are responsible for dominance remains to be evaluated. Notably, mRNA levels of all 11 *CKX* paralogs were reduced in reproductive tissues, indicating regulatory feedback. Consistent with the *CKX* mRNA level reduction, *trans*-zeatin riboside levels were elevated. Excessive cytokinin elevation disrupts source-sink relationships^34^, while modest increases observed in *KAMALA* enhanced yield while preserving agronomic stability^35, 36^. These findings indicate that fine-tuning of cytokinin metabolism, rather than complete disruption, may offer promising ways for sustainable crop improvement^35^.

Beyond the enhancement of several agronomic traits, all the edited *CKX2* lines exhibited complete panicle exertion, representing a previously unreported phenotype in *CKX2*-edited rice. This phenotype is particularly significant because incomplete/partial panicle exertion, commonly observed in Samba Mahsuri, is associated with decreased yield and increased disease risk due to moisture accumulation at the flag leaf-panicle junction^37, 38^. Recently, an EMS mutant of Samba Mahsuri with elevated mRNA levels for cytokinin biosynthetic genes demonstrated full panicle emergence, supporting the hypothesis that elevated cytokinin contributes directly to this key trait^39^.

Notably, multi-season field trials conducted at IIRR revealed that *KAMALA* demonstrated a 4% increase in HI, enhanced photosynthesis rates, and greater flag leaf area, essential for efficient grain filling. The HI is a critical parameter used for assessing reproductive efficiency, nutrient budgets, and stover carbon inputs. A recent study in maize reported an annual increase of 0.26% in HI, primarily attributed to genetic gain rather than crop management^40^. Despite the significant leap over agronomic traits, the success of any cultivar depends on its acceptability by rice millers and farmers. In particular, the market of Samba Mahsuri in India has been still intact after its ∼40 years of release, mainly because of its grain quality^41^. While the quantum leap in yields and culm strength were highly encouraging attributes, we were equally cautious about milling and grain quality of *KAMALA*. The critical milling and grain quality parameters like head rice recovery, hulling, milling, amylose content and gel consistency of *KAMALA* were equivalent to Samba Mahsuri. Earlier study indicated that mutation in *CKX2* does not affect grain quality^13^.

The early flowering trait observed across all four *CKX2*-edited lines under glasshouse conditions was recapitulated in multi-season field trials conducted at IIRR Hyderabad, a finding not previously reported for *CKX2*-edited rice lines. Early maturation provides considerable agronomic and environmental advantages, by shortening the growing season by 20-25 days, thereby reducing water, fertilizer, and labor needs, helping the crop escape terminal heat during the dry season, and lowering methane emissions^42^. In Arabidopsis, root-derived cytokinins play roles in flowering time by modulating *SPL* transcription factors and the miR156/172 microRNA cascade^43^. Cytokinin levels also were shown to be linked to gibberellin (GA) biosynthesis and signaling, establishing a dynamic cross-talk between CK and GA pathways that jointly govern meristem fate and developmental timing^44^. Consistent with a link to GA, mRNAs levels of GA pathway and floral organ identity genes, for example, *OsFTIP1* and floral meristem identity genes such as *OsMADS51, OsMADS18*, and *OsAP1* were increased, while mRNAs for repressors of flowering, including *OsHd1, OsDTH8*, and *OsPINE1* decreased.

A detailed analysis of the mechanisms leading to the striking effects of the *KAMALA* mutations, both genetically and physiologically may provide a basis for a more rational approach for engineering productivity. *KAMALA* can thus serve as a model for fundamental research to explore this surprising productivity increase. However, multi-year, multi-site field trials revealed no apparent trade-offs, except in Nellore and Pattambi, where *KAMALA* showed inferior performance, possibly due to effects of maturation time in conditions of coastal salinity in Nellore and heat in Pattambi. It will be interesting to explore whether similar mutations will yield in other crops. While farmers showed enthusiasm to adopt GE varieties, broader deployment continues to face challenges from concerned citizens (www.pib.gov.in/PressReleasePage.aspx?PRID=2194569&reg=48&lang=2; www.deccanherald.com/india/gm-free-india-coalition-slams-centre-for-releasing-genetically-modified-rice-varieties-3525792). Effective public communication was part of the effort towards releasing *KAMALA*. Proactive and pragmatic support by the Indian government was instrumental for advancing *KAMALA* close to deployment. The adoption of the technology in major importing regions, e.g. Europe, will be relevant for the global impact^45^ (www.healthycrops.org/). Globally coordinated regulatory genome editing frameworks, coupled with freedom-to-operate, could benefit millions of small and marginal farmers by improving income and livelihoods, while advancing environmental sustainability, food security, and resilient agricultural growth.

## Online Methods

### Plasmid construction

The genomic sequence of the *CKX2* gene (*LOC_Os01g10110*) in *Oryza sativa* L. was retrieved from the RAP-DB database (https://rapdb.dna.affrc.go.jp/)^46, 47^. Overlapping primers were designed for sequencing of the *CKX2* allele from the Samba Mahsuri. Two guide RNAs (gRNAs) were designed to target regions close to the cytokinin binding (gRNA-1) and FAD binding (gRNA-2) domains (Supplementary Table 11). The gRNAs were chemically synthesized and cloned into unit vectors under the control of *OsU6*, *OsU3*, *TaU3*, and *ZmU3* promoters. Each gRNA was cloned independently into two distinct unit vectors to ensure robust expression (Supplementary Fig. 4). Subsequently, the gRNA cassettes were assembled into the pENTR4-Cpf1-ccdb entry vector using the NEBridge® Golden Gate Assembly Kit (BsaI-HF® v2). The complete CRISPR/Cas12a editing module-including all four gRNAs and the Cas12a expression cassette was mobilized into the pBY02-rUbi-hlbCpf1_ccdb binary vector. In this construct, Cas12a is driven by the rice ubiquitin promoter (*rUbiP*), and hygromycin resistance–based selection is facilitated by the *hptII* gene under the control of the CaMV 35S promoter. The gRNA array was assembled in the following order: gRNA-1–gRNA-2–gRNA-1–gRNA-2 (Supplementary Fig. 4). The binary construct was introduced into *Agrobacterium tumefaciens* strain EHA105 using the freeze-thaw method^48^. A complete list of primers used for cloning and validation is provided in Supplementary Table 12.

### Plant materials, growth conditions and generation of CRISPR plants

Genetically pure seeds of Samba Mahsuri were obtained from the Agricultural Research Station Bapatla of Acharya NG Ranga Agricultural University and were subsequently maintained in every growing season at ICAR-Indian Institute of Rice Research, Hyderabad, India. Mature seeds were used for embryogenic calli formation on MS basal media (Cat#M5519, Sigma-Aldrich) supplemented with 30 g/L sucrose and 2.5 mg/L 2,4-D (2,4-dichlorophenoxyacetic acid) in the dark^49^. Calli were transformed with *Agrobacterium tumefaciens* strain EHA105 harboring pBY02*gRNA:ckx3-4 as described earlier^50^. Subsequently, the co-cultivation and selection process on 50 mg/L hygromycin were followed to generate the transgenic calli. Putative transgenic calli were regenerated on MS basal media supplemented with 30 g/L maltose and 2.5 mg/L kinetin at 16 h light/8 h dark photoperiod^51^. Green shoots were transferred to the half MS media without any hormone for root development and plantlets formation. Initial plantlets were acclimatized in the liquid Yoshida media. The acclimatized plants were transplanted to soil pots and shifted to a biosafety glasshouse for growth and seed setting (Supplementary Fig. 6).

### Confirmation of transgene integration

For genotyping of the T0 transgenic lines, leaf samples were collected, frozen in liquid nitrogen, and stored at -80 °C. The DNA isolation from leaves was performed as per CTAB methodology described by Aboul-Maaty and Oraby^52^. The vector pBY02 backbone-specific PCR primers designed from Cas12a, hpt, and 35S CaMV promoter regions were used for detection of transgene integration (Supplementary Fig. 30). Transgene-positive plants were screened by amplification of the target sequences (E3 and E4), followed by Sanger sequencing to confirm target gene editing (Supplementary Figs. 5,7).

### PCA analysis of edited lines and identification of biallelic homozygous mutants

The mutations in target regions of genome-edited lines were identified in T0 generation through PCR product sequencing. To validate the clustering patterns observed in PCA space, K-means clustering was performed on the PC1 and PC2 coordinates. Silhouette analysis identified three optimal clusters (silhouette score = 0.7285), indicating strong separation and good clustering quality. Statistical validation using one-way ANOVA revealed highly significant differences (*P* < 0.001) between clusters for all three traits: grains per panicle (F = 104.28), number of tillers (F = 141.11), and early maturity (F = 257.35), confirming that the clusters represent distinct biological groups. Cluster 0 (n=39) was characterized by higher grain productivity (352.05 ± 8.79 grains per panicle), fewer tillers (5.56 ± 0.50), and later maturity (142.18 ± 1.71 days), representing genotypes optimized for grain production at the expense of tillering and early maturity. Cluster 1 (n=32) showed lower grain productivity (287.81 ± 35.54 grains per panicle), more tillers (7.69 ± 0.69), and earlier maturity (128.16 ± 3.67 days), representing early-maturing genotypes with moderate productivity. Cluster 2 (n=7) was the most desirable, combining the highest grain productivity (418.57 ± 28.54 grains per panicle), high tillering (8.57 ± 0.79), and early maturity (127.43 ± 2.57 days), representing an ideal combination that breaks typical trade-offs.

Seeds were collected from these genome-edited lines and advanced (by selfing) to T1, T2, and T3 generations under a biosafety screenhouse (Supplementary Fig. 6c). From each line, 100 plants were screened in T1 and T2 generations to identify homozygous bi-allelic mutations and to confirm effective outcrossing of transgene fragments using overlap PCR. Amplicons were obtained from 39 overlapping primer sets covering the complete binary plasmid. Each PCR amplicon was <500 bp and the overlap between consecutive PCR amplicon was of least 50 bp. Each PCR reaction included endogenous control (actin), transgene positive control (DNA from T0 transgene positive line), PCR sensitivity control (DNA of WT spiked with 1/1000^th^ (w/w) of the plasmid DNA), negative control (WT DNA). Each reaction included one of the 39 overlapping primer sets and the controls mentioned above (Supplementary Fig. 12). We did not detect PCR amplification with vector backbone primers in the four genome edited lines, while expected PCR fragments were detected in all positive controls. In addition, a hygromycin sensitivity test was performed for these selected lines. Seeds from the edited lines failed to grow in growth media with 70 mg/L hygromycin. By comparison transgene positive seeds of same line (T0 seeds) were able to grow (Supplementary Fig. 13). To confirm the homozygosity of mutations, both DNA strands of the target regions were sequenced across two generations (T2 and T3), using five independent PCR amplifications followed by Sanger sequencing in each generation.

### Evaluation for agronomic performance of four homozygous edited lines without detectable transgenes

Four homozygous mutant lines-*KAMALA*, GeD2, GeD3, GeD4 along with the WT were evaluated under biosafety screenhouse conditions. These lines were grown in a 10×10 m² (∼330 plants) block design experiment with three replications (Supplementary Fig. 6c). The days to 50% flowering, plant height (n = 50), number of tillers (n = 500), panicle length (n = 500), branches (primary) per panicle (n = 250), panicle branch length (n = 200), grains per panicle (n = 160 primary panicles), grain length (n = 310), grain width (n = 300), thousand-grain weight (n = 160 batches of 1000 grains), dry weight of panicle (n = 192 panicles), unfilled grains per panicle (n = 70), and yield per plant (n = 250) were recorded.

### CKX2 variants, modelling, and structural flexibility analysis

The natural variations in rice *CKX2* were analyzed using the RiceVarMap database (https://ricevarmap.ncpgr.cn), comprising 4,726 accessions representing diverse subpopulations (indica, japonica, aus, aromatic, and admixed) from multiple geographical regions. Variants were extracted from the *CKX2* genomic region, including 1 kb upstream and downstream flanking sequences to capture both coding and regulatory polymorphisms. Single Nucleotide Polymorphisms (SNPs) and INDELs were retrieved using the RiceVarMap interface, and variants with incomplete annotation or poor quality were excluded. Subsequent analyses were restricted to protein-coding regions. Three dimensional (3D) structural models of CKX2 variants were predicted using AlphaFold3^53^ (Supplementary Note 3). Three-dimensional structures highlighting the cavities were computed using CavitOmiX. The analysis of structural flexibility of proteins was done using CABS-Flex 3^23^. We used Boltz-2^54^ to predict the structural complexes with iP (N6-(2-isopentenyl) adenine) and tZ (trans-zeatin). The multiple sequence alignment of deduced amino acids of CKX2 of *KAMALA*, GeD2, GeD3, and GeD3 was generated using BioEdit version 7.0^55^ (Supplementary Fig. 31).

### Biosafety clearance of *KAMALA*

Based on agronomic performance, *KAMALA* was shortlisted to develop the necessary data and information to get an exemption from GM rule 1989 (http://www.geacindia.gov.in/acts-and-rules.aspx). The data was generated as per the guidelines of SOPs for Regulatory Review of SDN-1 & SDN-2 Genome Edited Plants, 2022 (https://dbtindia.gov.in/sites/default/files/SOPs%20on%20Genome%20Edited%20Plants_0.pdf), released by Department of Biotechnology, Ministry of Science & Technology, Government of India. The dossier was prepared and submitted to IBSC (Institutional Biosafety Committee) and RCGM (Review Committee on Genetic Manipulation). After careful scrutiny of the data, the biosafety regulatory bodies gave exemption to *KAMALA* in May 2023.

The genomic DNA of WT and *KAMALA* were subjected to PacBio HiFi WGS. Sequencing quality was assessed using SAMtools v1.18. Alignment statistics including total read count, mapping rate, and supplementary alignment frequency were obtained using SAMtools flagstat. Read-level metrics were extracted using SAMtools stats (Supplementary Figure 14a). Reads from WT Samba Mahsuri and *KAMALA* were mapped to the transformation vector sequence using minimap2 v2.17 (map-pb preset, k-mer seed size of 19 bp, single secondary alignments disabled). Per-base sequencing depth across the vector was computed using SAMtools v1.18. Coverage was summarised in 100 bp bins, retaining regions with ≥3× depth across ≥25% of bases per bin (Supplementary Figure 14b). Reads overlapping high-coverage bins were extracted and remapped to the annotated vector, and alignments were intersected with vector feature annotations using BEDTools v2.26. Sequence analysis revealed no mutations in the other *CKX* family members, which was further supported by an in-silico guide RNA–binding analysis of the remaining 10 *CKX* paralogs (Supplementary Fig. 32; Supplementary Table 10).

### Yield evaluation of *KAMALA* under multi-location field trials

Field evaluation of *KAMALA* along with its WT was done at the research field of the IIRR Hyderabad for three seasons [wet season (Kharif) of 2023, dry season (Rabi) of 2024, and wet season of 2024]. The evaluation was done in a randomized block design (RBD) with three replications. Each replication had a plot size of 1000 m^2^. First the seeds were grown in nursery to obtain 21-day-old seedlings. These seedlings were transplanted into field plot with a row-to-row spacing of 20 cm and 15 cm between plant-to-plant. A recommended fertilizer amount @ 120 kg nitrogen (urea), 60 kg phosphorus (single super phosphate), 60 kg potassium (muriate of potash), and 25 kg zinc (zinc sulphate) per hectare was applied. Except nitrogen, all other fertilizers were applied as basal dose, while nitrogen was applied as 50% basal dose, 25% at 30 days after transplantation, and remaining 25% at 60 days after transplantation. Regular irrigation was provided to maintain 5 cm water in field until maturity. For yield estimation, each plot was harvested individually, and seeds were threshed and weighed.

Similarly, *KAMALA* and WT were evaluated under multi-location field trials of AICRPR as per the standard procedures. *KAMALA* was coded as IET 32072, and was evaluated by AICRPR scientists in three agro-climatic zones, i.e. Zone III (comprising states/provinces-Odisha, Jharkhand, Bihar, Uttar Pradesh and West Bengal), Zone V (Chhattisgarh, Madhya Pradesh and Maharashtra states), and Zone VII (Andhra Pradesh, Telangana, Karnataka, Tamil Nadu, Kerala and Puducherry states). It was evaluated for two seasons (wet season of 2023 and 2024) in zone III and V, while three seasons (wet season 2023, dry season 2024, wet season 2024) in zone VII. Rice is not grown in dry season of zone III and V due to low temperatures associated with cold weather conditions. In each location of AICRPR, *KAMALA* and WT were evaluated in a RBD with three replications (each replication with plot size of 15 m^2^). Here, 21-day-old seedlings were transplanted with a row-to-row spacing of 20 cm and 15 cm between plant-to-plant. The recommended dose of fertilizer and irrigation were provided as per the standard procedures under AICRPR. The analysis of variance (ANOVA) was conducted using data generated from a randomized complete block design (RCBD) across multiple locations. The pooled ANOVA was performed following standard statistical procedures for combined analysis. The analysis was executed in SAS software, version 9.3, using the PROC GLM procedure (SAS Institute Inc., 2011). The least significant difference (LSD) test was applied as a post hoc comparison to assess the performance of the experimental entries. It should be noted that the data only from qualified locations [based on monitoring team (a team of independent scientists deputed by AICRPR) report on proper conduct of trial and statistical significance of trial] is considered by AICRPR unit for data analysis, and the same criteria were applied for *KAMALA*.

### Phenotypic characterization of *KAMALA* in field conditions in 2023 and 2024

Data were collected from the research field of IIRR at Hyderabad. The vigor index was calculated at 21-days seedling stage using the formulas for VI-1 = Seedling Length × Germination %, and VI-2 = Seedling Dry Weight × Germination %^56^. Root diameter, number of crossings, forks, and tips were analyzed at 45 days after sowing using a root scanner and WinRHIZO from 15 randomly selected plants each of WT and *KAMALA*. At the same stage, above-ground physiological parameters were noted that includes leaf width (n = 78) and culm width (n = 75), measured using a scale and vernier calliper, respectively. Also, the photosynthetic pigments-chlorophyll-a, chlorophyll-b and total carotenoids, were measured in randomly selected 30 plants each, and calculated the quantity as per the Arnon equations^57^. At the reproductive stage, root and shoot lengths of 285 randomly selected plants were measured, and the root-to-shoot length ratio was subsequently calculated. Out of these, 160 plants each were used to estimate the root fresh and dry weight as well as shoot fresh and dry weight, and the resulting data were used for the calculation of root/shoot fresh and dry weight ratio. Similarly, 400 plants each of WT and *KAMALA* were used to measure flag leaf width and length at the booting stage. Photosynthetic parameters such as photosynthetic rate, stomatal conductance, transpiration rate and intracellular CO_2_ concentration, were measured in 25 flag leaves from independent plants using a portable photosynthesis system LI-COR Biosciences device (Nebraska, USA) at PAR 600–1200 nm. Measurements were taken in the morning hours of 09:00–11:30 AM. Approximately 35 fully mature plants were used to calculate the HI using the formula: HI (%) = Grain yield / Biological yield × 100.

The DIK-7401 Prostrate Tester (Daiki Rika Kogyo Co., Ltd., Japan) was used to measure the culm (stem) strength after 20 days of flowering. All readings were taken 20 cm above from the ground, and the stem was carefully bent at a 45° angle to record the load-bearing capacity of stem. The readings of WT and *KAMALA* were converted into kg/cm units based on the reference table provided with the instrument manual.

### Grain quality parameters

Post-harvest mature seeds were properly sundried and analyzed for the standard grain quality parameters of AICRPR, i.e., head rice recovery (%), hulling (%), milling (%), amylose content, and gel consistency^58^. Quality trait means and standard errors were calculated from ten independent batches of grain harvested from a single season.

**Tissue staining and microscopy, RNA isolation, RNA sequencing and RT-qPCR, measurement of cytokinin oxidase activity, phytohormone extraction and profiling**: methods described in Supplementary Note-4.

### Allele specific DNA markers for *KAMALA* and introgression of allele into rice cultivars

The DNA markers to identify deletion of 18 and 9 bp in *KAMALA* allele, were designed from E3 (forward 5’-CTCACTTTATCCTTGCATGAC-3’; reverse 5’-TGAGGGGTCGTCATTTTG-3’) and E4 (forward 5’-TGGTGGTAACTAAACATA-3’; reverse 5’-ATGCACAGTACACCCATA-3’) of *CKX2*, respectively. Primer performance was validated in *KAMALA,* WT, and 30 diverse genotypes. The CAPS markers were developed based on the abrogation of the *Nru*l restriction enzyme site due to deletion of an 18 bp sequence in E3 of *KAMALA*. Detailed protocols for the identification of *KAMALA* allele, including PCR components, primers, PCR and gel electrophoresis conditions, and restriction enzyme analysis procedures, are provided in Supplementary Fig 22.

### Genetic analysis of *KAMALA-CKX2*

To investigate the genetic basis of the mutation, *KAMALA* was crossed with Improved Samba Mahsuri (a near-isogenic line of Samba Mahsuri) and the rice variety DRR Dhan 48. *KAMALA* was used as the male parent in all crosses to eliminate potential cytoplasmic effects during genetic analysis. The resulting F plants were backcrossed with the respective female parent to generate BC F progeny. These were subsequently advanced through selfing to obtain BC F, BC F, BC F, and BC F generations. All BC F plants were screened using a *KAMALA* allele-specific DNA marker, and homozygous and heterozygous plants were identified and tagged accordingly.

### Data analysis and figure generation

Edits at desired locus and chromatogram alignment were checked and analyzed in BioEdit version 7.0^55^ and Synthego’s ICE CRISPR analysis tool (https://www.synthego.com/). All the edited alleles of *CKX2* gene were translated to obtain protein sequences. The protein versions of CKX2 were analyzed using UniProt (https://www.uniprot.org/) and ProtParam (https://web.expasy.org/protparam/) bioinformatics tools. Similarity and differences among the amino acid sequences of translated proteins were demonstrated through Multiple Sequence Alignment by CLUSTALW. Data analysis was performed using Microsoft Excel (version 7), and graphs and scatter plots were generated using GraphPad Prism software (https://www.graphpad.com/). Data are presented as mean ± s.e.m. (standard error of the mean). Numbers of measurements and biological replicates are specified in respective sections describing each trait. For comparisons involving more than two groups, a non-parametric Kruskal–Wallis test followed by Dunn’s post hoc test was employed to determine significant differences for the measured trait. When comparisons involved only two groups, statistical significance was evaluated using a two-tailed Student’s t-test. One-way ANOVA was conducted on all measurements. Tukey’s honestly significant difference test was used for post-ANOVA pairwise tests for significance (set at *P* <0.05). Exact *P* values, statistical tests used and sample numbers (n) can be found in figure legends or in graphs. Figures were produced using elements from BioRender.com.

## Data Availability

RNAseq data is available online (IBDC Study Accession: INRP000477; INSDC SRA Project Accession: PRJEB100853). All the other data is submitted in manuscript main figures or supplementary information. The references cited in supplementary figures and tables are provided as supplementary note 5. The multi-location field evaluation data is available online at AICRPR website (http://www.aicrip-intranet.in/). The *KAMALA* code is IET 32072. The materials developed in this study will be made available as per the rules and guidelines of ICAR, Government of India, biosafety regulatory bodies, and other relevant national-international guidelines. The *CKX2-GE1* mutant was designated *KAMALA*, and the mutation is protected by patents filings (Indian Patent Office, No.202311030876 A, PCT/IB2024/054133). The data that support the findings of this study are available from the corresponding author upon request.

Correspondence and requests for materials should be addressed to Satendra K. Mangrauthia.

## Acknowledgments

Financial support for this study was received from ICAR-NASF project entitled “Genetic improvement of rice for yield, NUE, WUE, abiotic and biotic stress tolerance through RNA guided genome editing (CRISPR-Cas9/Cpf1)” (NASF/CRISPR-Cas-7003/2017-18), and “CRISPR Crop Network: Targeted improvement of stress tolerance, nutritional quality and yield of crops by using genome editing” (NASF/ BGAM-9021/2022-23). WBF was supported by an Alexander von Humboldt professorship. Authors thank researchers of different AICRPR centers involved in multi-location field evaluation of *KAMALA*. We acknowledge the phytohormone analysis facility provided by Repository of Tomato Genomics Resources (RTGR), a DBT-Sahaj national facility via grant “Research and Service facilities for Plant Metabolomics and Proteomics” (BT/INF/22/SP44787/2021).

## Contributions

S.K.M, M.S, R.M.S conceived and designed experiments. M.S, S.K.M, F.Y, S.K, E.R, S.C, A.R, B.S, S.V.S.P, J.A.K, Brajendra, A.F.R, and A.S.S performed experiments. V.C, R.M.S, C.N.N, W.B.F, and B.Y provided resources. M.S, S.K.M, E.R, G.K.S, W.B.F, E.L, S.V. S.P analyzed the data. M.S, S.K.M, E.R, and W.B.F. wrote the manuscript.

