## Supplementary Figures for "*KAMALA:* improved yield by finetuning cytokinin oxidase activity in rice"

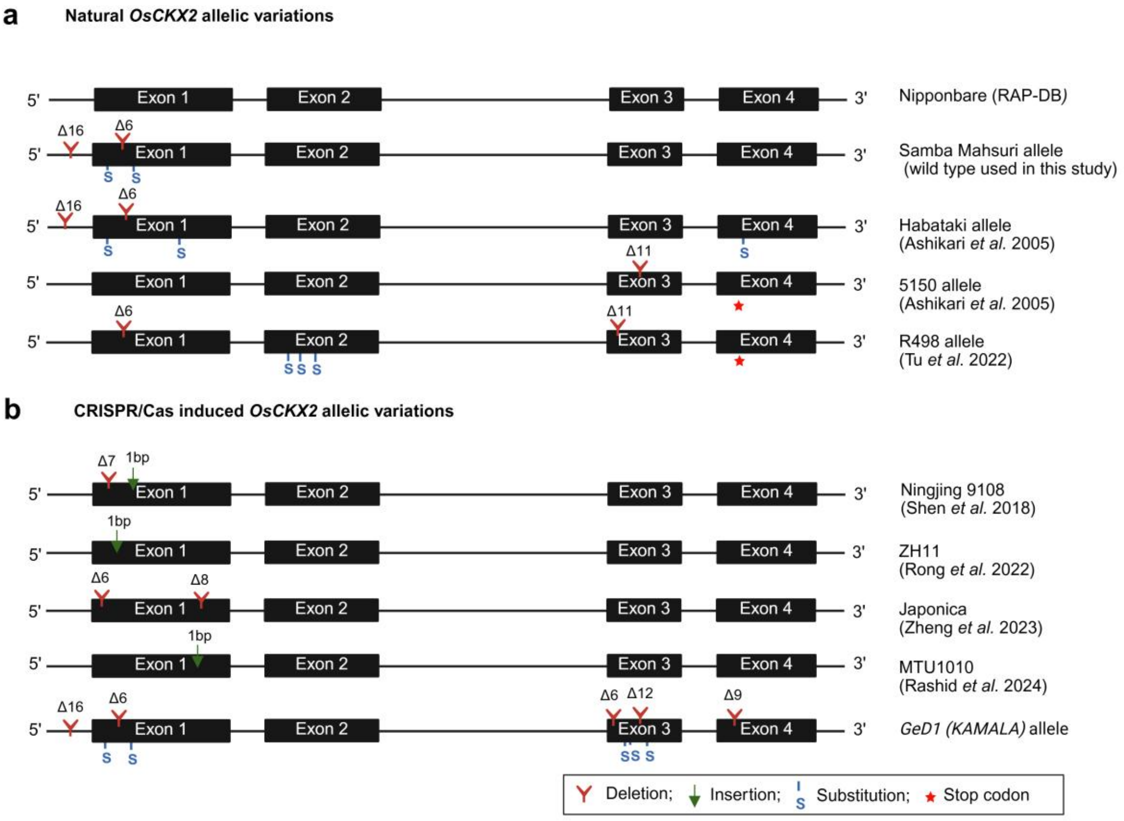


**Supplementary Fig.1| A summary of *OsCKX2* major allelic variations. a**, natural allelic variations of *OsCKX2* (Nipponbare sequences were obtained from RAP-DB portal; Habataki and 5150 allelic variations were obtained from Ashikari *et al.* 2005; R498 allele information obtained from *Tu et al.* 2022). **b**, CRISPR/Cas induced *OsCKX2* allelic variations (Ningjing 9108 from Shen *et al.* 2018; ZH11 from Rong *et al.* 2022; Japonica from Zheng *et al.* 2023 and MTU1010 from Rashid *et al.* 2024). GeD1 (*KAMALA)* allele developed in this study.

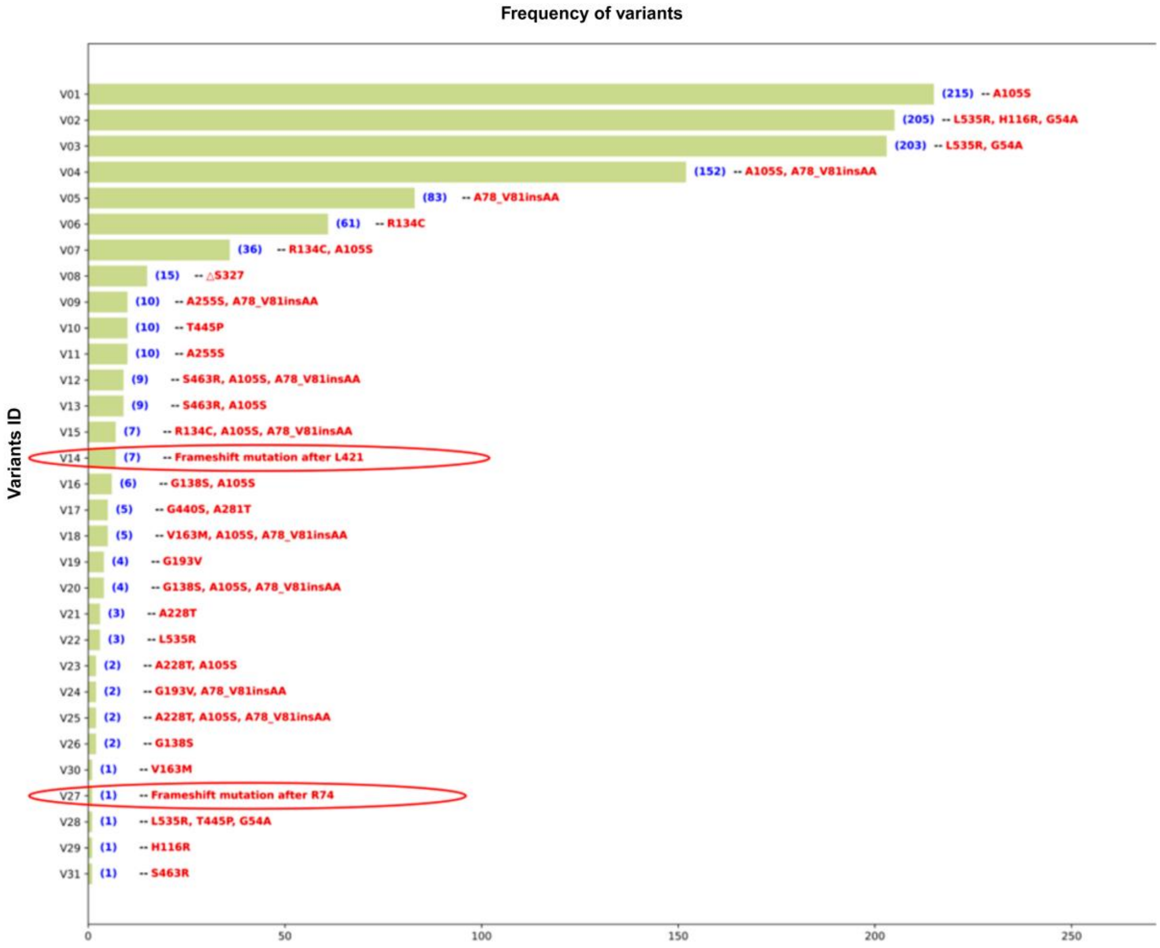


**Supplementary Fig.2| Natural CKX2 protein variants.** Horizontal bar plot illustrating the distribution of single- and multiple-amino acid variant combinations identified in CKX2 among 4,726 global rice accessions available in the RiceVarMap database. The x-axis represents the number of cultivars/genotypes associated with each variant combination, while the y-axis denotes the corresponding variant IDs (V01–V31). Genotypes with no amino acid variation in CKX2 are not included in the plot. The details of genotype and mutations are included as Supplementary Table 1.

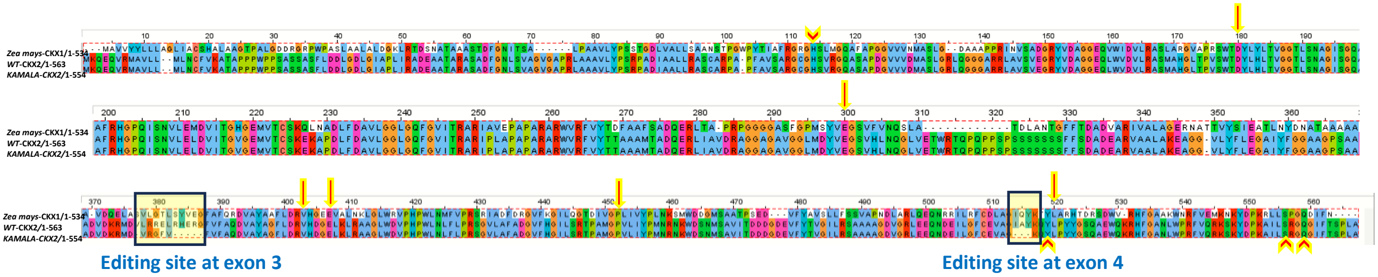


Critical residues involved in cytokinin binding

| CKX-Maize | Samba Mahsuri (WT) | *KAMALA* |
|---|---|---|
| **D169** | D177 | D177 |
| **E288** | E297 | E297 |
| **V378** | V399 | V393 |
| **E381** | E403 | E397 |
| **P427** | P448 | P442 |
| **L492** | L516 | L507 |

Cytokinin binding residue

FAD binding residue

Editing sites

**Supplementary Fig.3| The identity and position of cytokinin and FAD binding residues in WT parent and *KAMALA* CKX2.** The critical substrate and cofactor binding residues were predicted using the kinetics and X-ray crystallography information available from CKX of *Zea mays* (Kopečný et al 2016). It was identified by performing the multiple sequence alignment between the amino acid sequences using the MUSCLE (Edgar, 2004) method implemented in Jalview (Waterhouse et al 2009). Additionally, the FAD binding residues were identified from UniProt database also (https://www.uniprot.org/uniprotkb/Q4ADV8/feature-viewer).

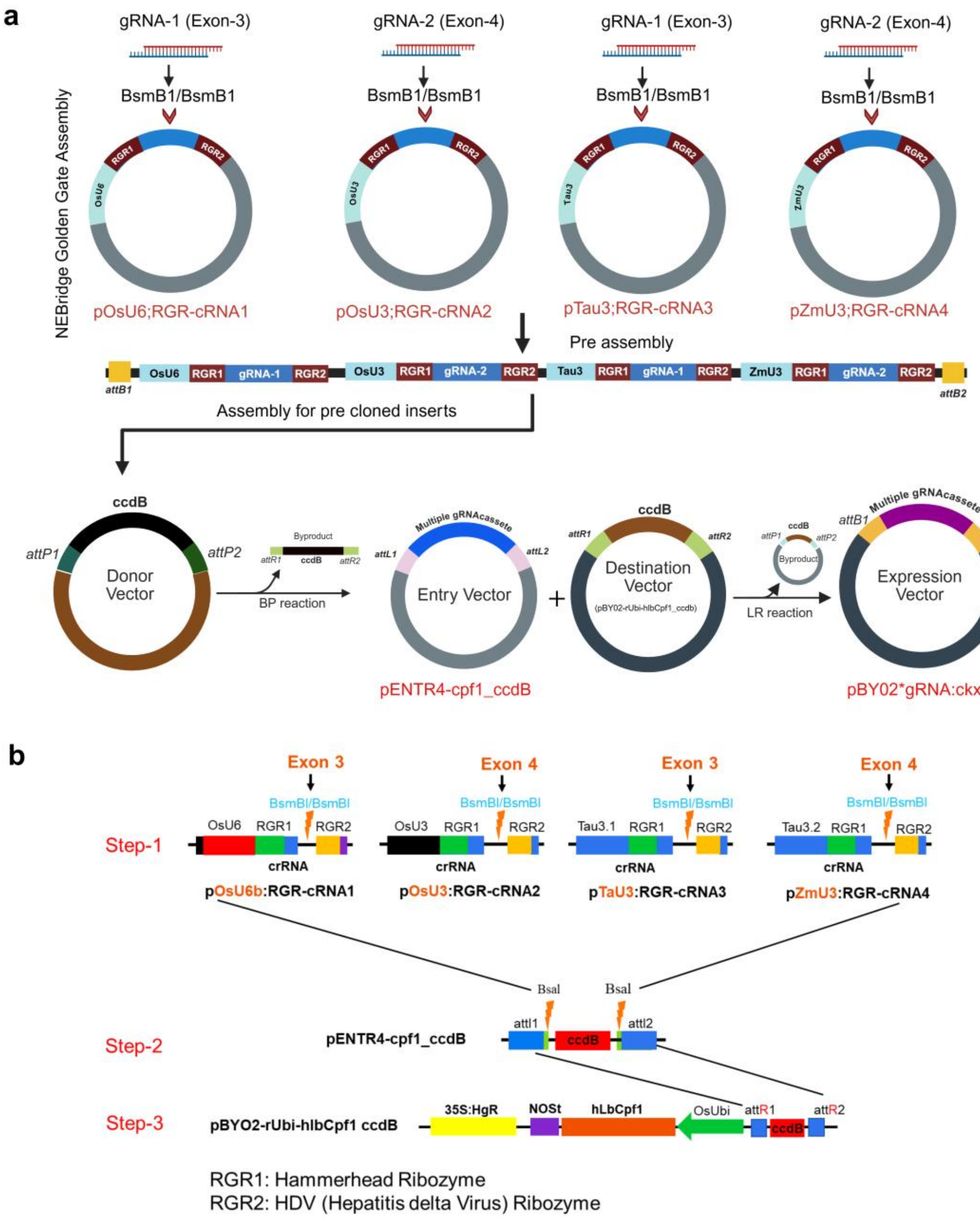


**Supplementary Fig. 4| Schematic representation of the pBY02*gRNA:ckx3-4 construct preparation. a**, Overview of the NEBridge Golden Gate Assembly process used to integrate gRNA cassettes (gRNA-1 and gRNA-2) into pre-cloned inserts, followed by Gateway cloning (BP and LR reactions) to generate the final expression vector. **b**, Step-wise assembly strategy: Step 1 involves the generation of crRNA-expressing cassettes via Golden Gate assembly; Step 2 demonstrates the cloning of these cassettes into the pENTR4-cpf1_ccdB donor vector; Step 3 illustrates the final LR reaction to produce the pBYO2*gRNA:ckx3-4 expression vector, containing the hlbCpf1 nuclease.

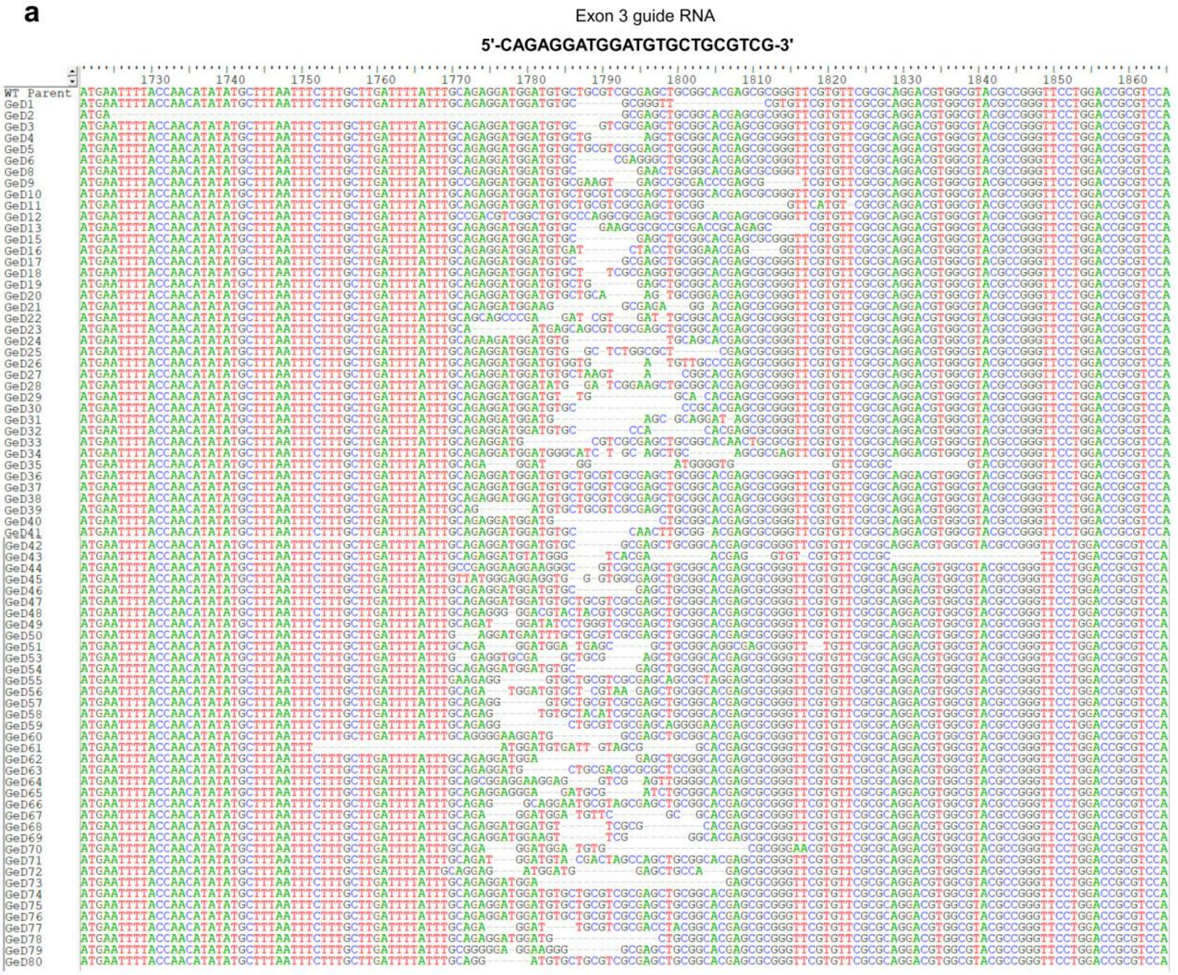


**Supplementary Fig. 5a| Sequence alignment of CRISPR/Cas12a induced mutations at the exon 3 target locus.** Multiple sequence alignment of independent mutants (GeD1-GeD80) compared to the unedited WT Parent sequence. The 23-nucleotide exon 3 guide RNA sequence (5'-CAGAGGATGGATGTGCTGCGTCG-3') used for targeted editing is displayed at the top. The alignment shows various insertion and deletion (indel) mutation profiles generated at the expected cleavage site. Dashes (-) represent deleted nucleotides, while standard nucleotide letterings represent retained sequence or substitution.

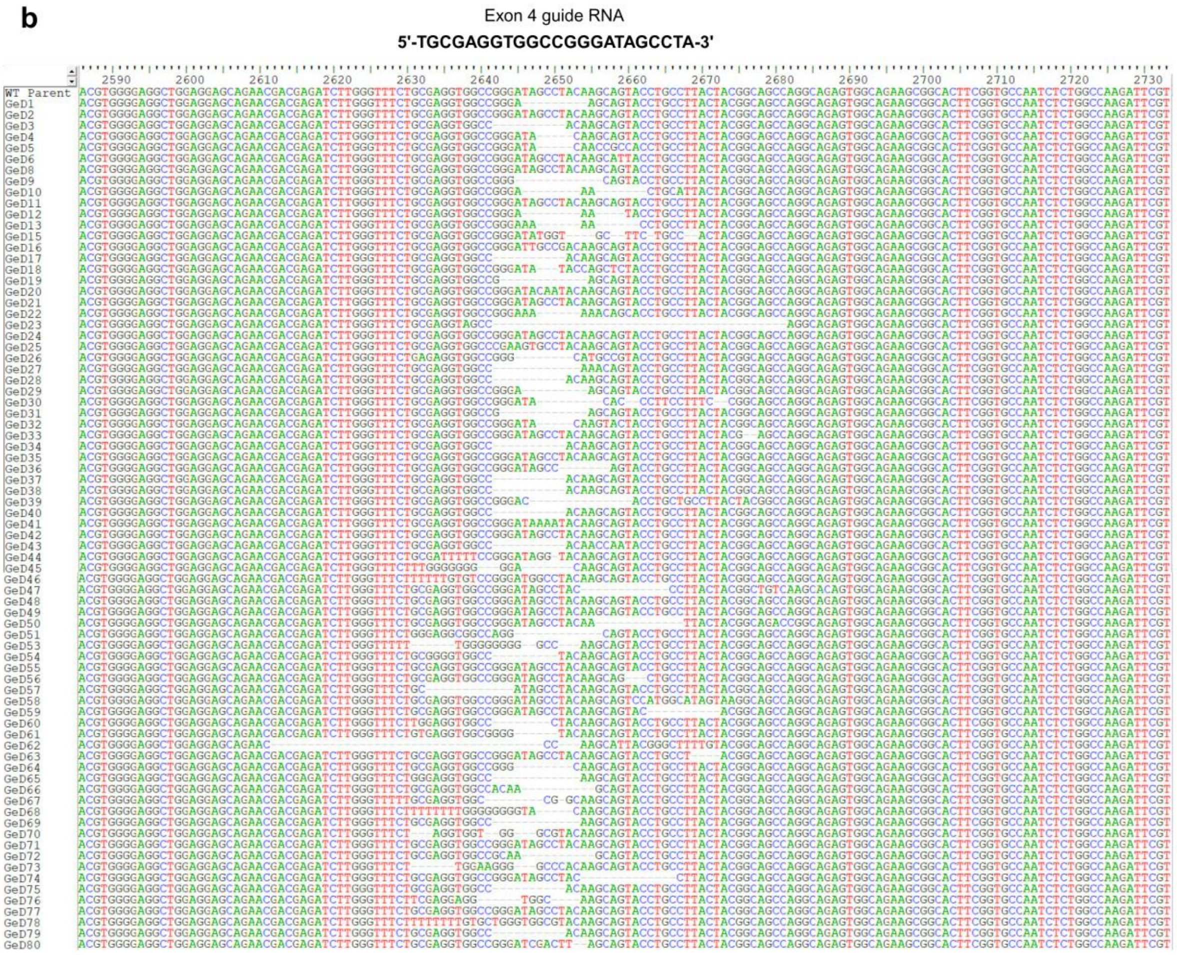


**Supplementary Fig. 5b| Sequence alignment of CRISPR/Cas-induced mutations at the Exon 4 target locus.** Multiple sequence alignment of independent mutants (GeD1-GeD80) compared to the unedited WT Parent sequence. The Exon 4 guide RNA sequence (5'-TGCGAGGTGGCCGGGATAGCCTA-3') used for targeted editing is displayed at the top. The alignment shows various insertion and deletion (indel) mutation profiles generated at the expected cleavage site. Dashes (-) represent deleted nucleotides, while standard nucleotide letterings represent retained sequence or substitutions.

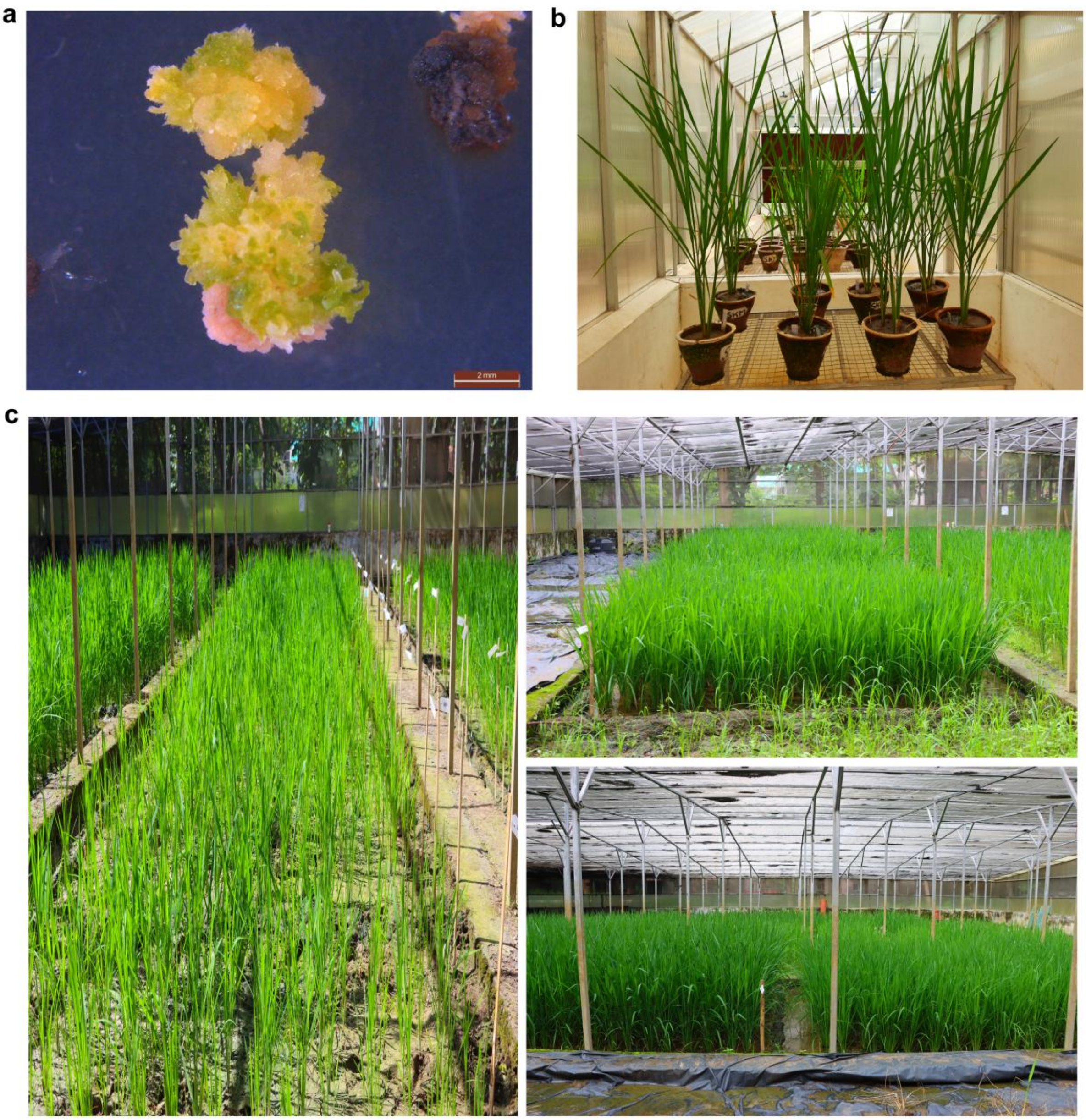


**Supplementary Fig. 6 | Schematic development and phenotypic progression of GeD1 (*KAMALA)* from tissue culture to biosafety screening. a,** Greening and regeneration of Samba Mahsuri rice calli following *Agrobacterium*-mediated transformation with the CRISPR/Cas vector *pBY02-sgRNA:OsCKX2-Ex3/4*. **b**, Plants from the T0 generation were acclimatized and their phenotypes were evaluated under controlled conditions in a Biosafety glasshouse. **c**, Phenotype evaluation of T1 and T2 progeny. These lines were grown and screened for agronomic performance under regulated conditions in the Biosafety Screenhouse at the ICAR-Indian Institute of Rice Research (IIRR), Hyderabad, India.

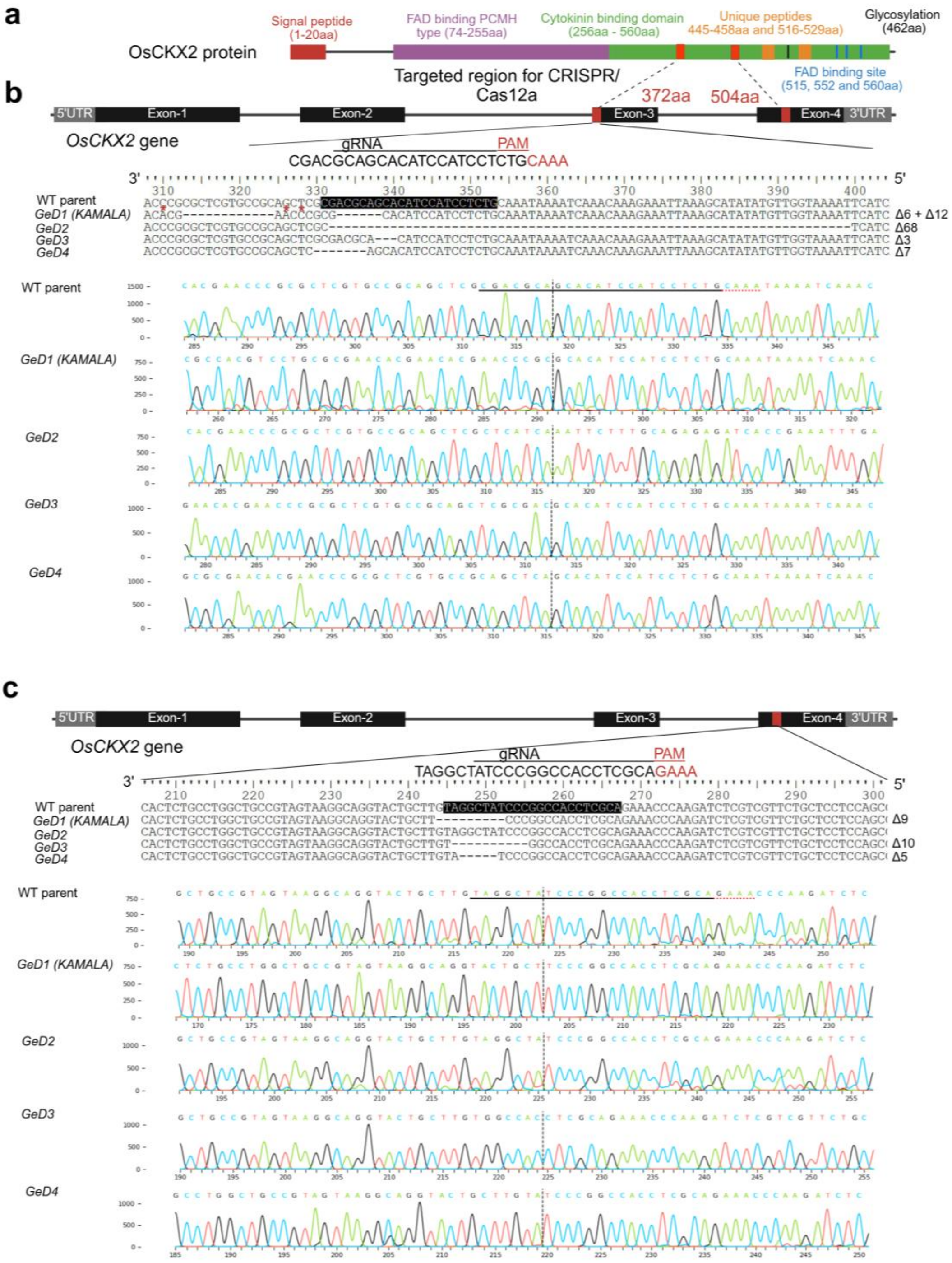


**Supplementary Fig. 7 | CRISPR/Cas12a-mediated multiplex editing of two sites close to cytokinin and FAD-binding residues of CKX2 in rice variety Samba Mahsuri (WT parent). a,** Domain architecture of the OsCKX2 protein. The CRISPR/Cas12a primary target region is located within the C-terminal cytokinin-binding domain (residues 256-560), which encompasses essential functional features including unique peptide motifs (445-458, 516-529), a glycosylation site (462), and FAD-binding residues (515, 552, 560). **b, c,** Strategic editing of novel sites in exon 3 **(b)** and exon 4 **(c)**. Schematic diagrams indicate the genomic location of the target sites (red boxes), gRNA sequences, and PAM sequences. Representative Sanger sequencing chromatograms (bottom) confirm the mutations in four independent genome-edited lines (GeD1 to GeD4) relative to the WT parent. Red asterisks denote nucleotide substitutions; specific deletion sizes are indicated to the right of each sequence alignment. Comprehensive chromatogram analysis is provided in Supplementary Figs. 7–10.

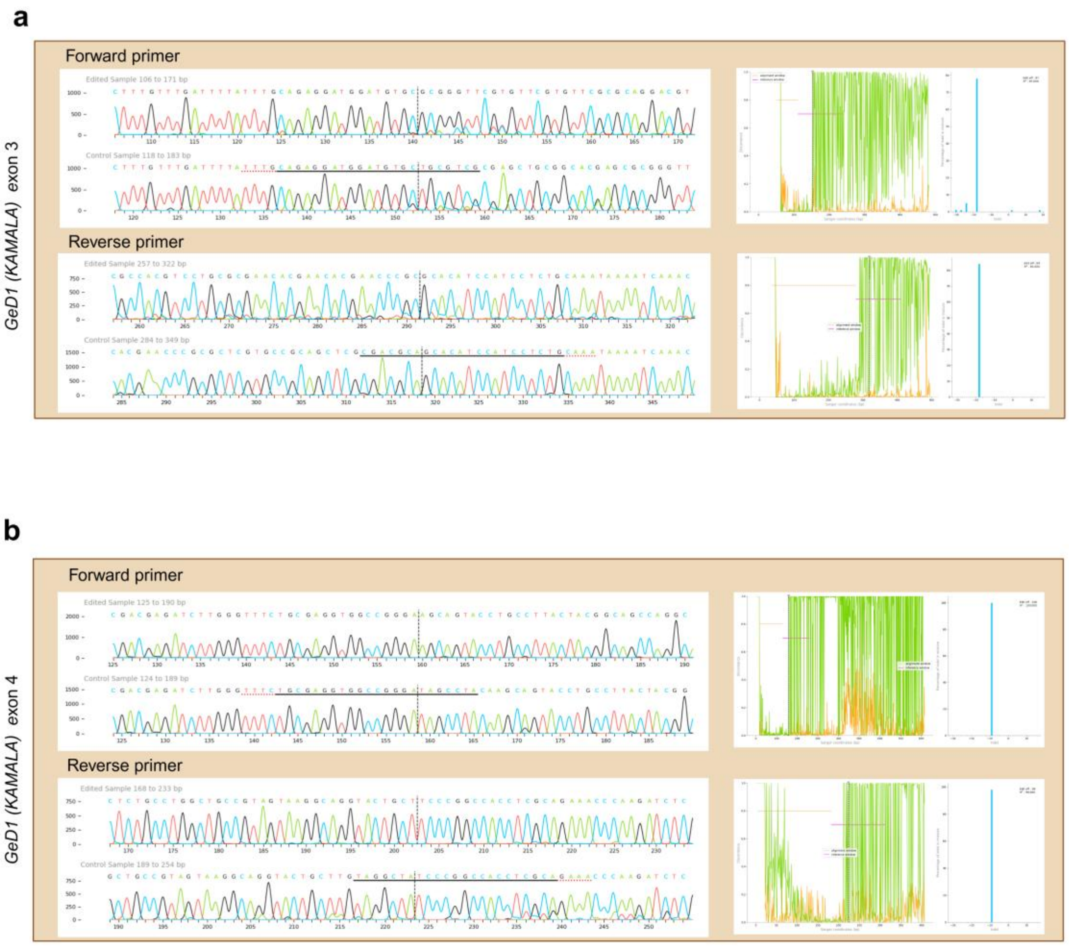


**Supplementary Fig. 8| Chromatogram alignment of *CKX2* gene between WT parent and GeD1 *(KAMALA)*.** The chromatogram alignment of +Ve and -Ve strand of the *CKX2* gene at exon 3 and exon 4 target sites. The underline sequences represent the sgRNA of the particular strands. The mean of discord score before the first cut site and after the cut site is also shown. CRISPR edit at desired location and chromatogram alignment were checked and analyzed in BioEdit and ICE CRISPR analysis tool provided by the SYNTHEGO (https://www.synthego.com/).

**a**

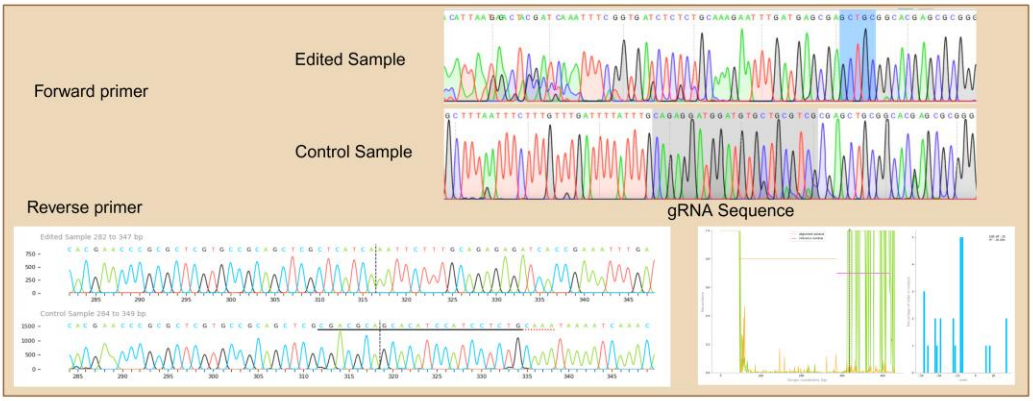


**b**

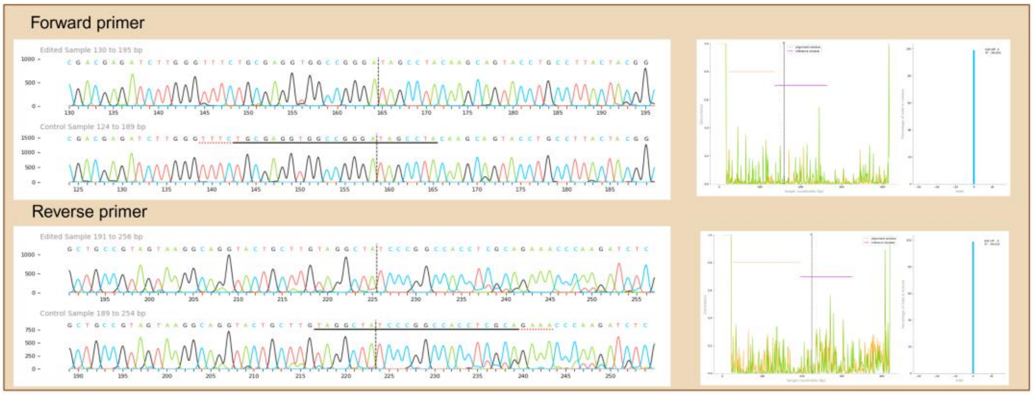


**Supplementary Fig. 9| Chromatogram alignment of *CKX2* gene between WT parent and GeD2.** The chromatogram alignment of +Ve and -Ve strand of the *CKX2* gene at exon 3 and exon 4 target site. The underline sequences represent the sgRNA of the particular strands. The mean of discord score before the first cut site and after the cut site is also shown. CRISPR edit at desired location and chromatogram alignment were checked and analyzed in BioEdit and ICE CRISPR analysis tool provided by the SYNTHEGO (https://www.synthego.com/). Note: The forward allele of exon 3 could not be analyzed using the Synthego ICE CRISPR analysis tool for undetermined reasons (potentially due to a large deletion spanning the guide RNA target site). However, the editing event was confirmed via sequence alignment using BioEdit. Additionally, the SnapGene-analyzed chromatogram for the forward allele is presented here.

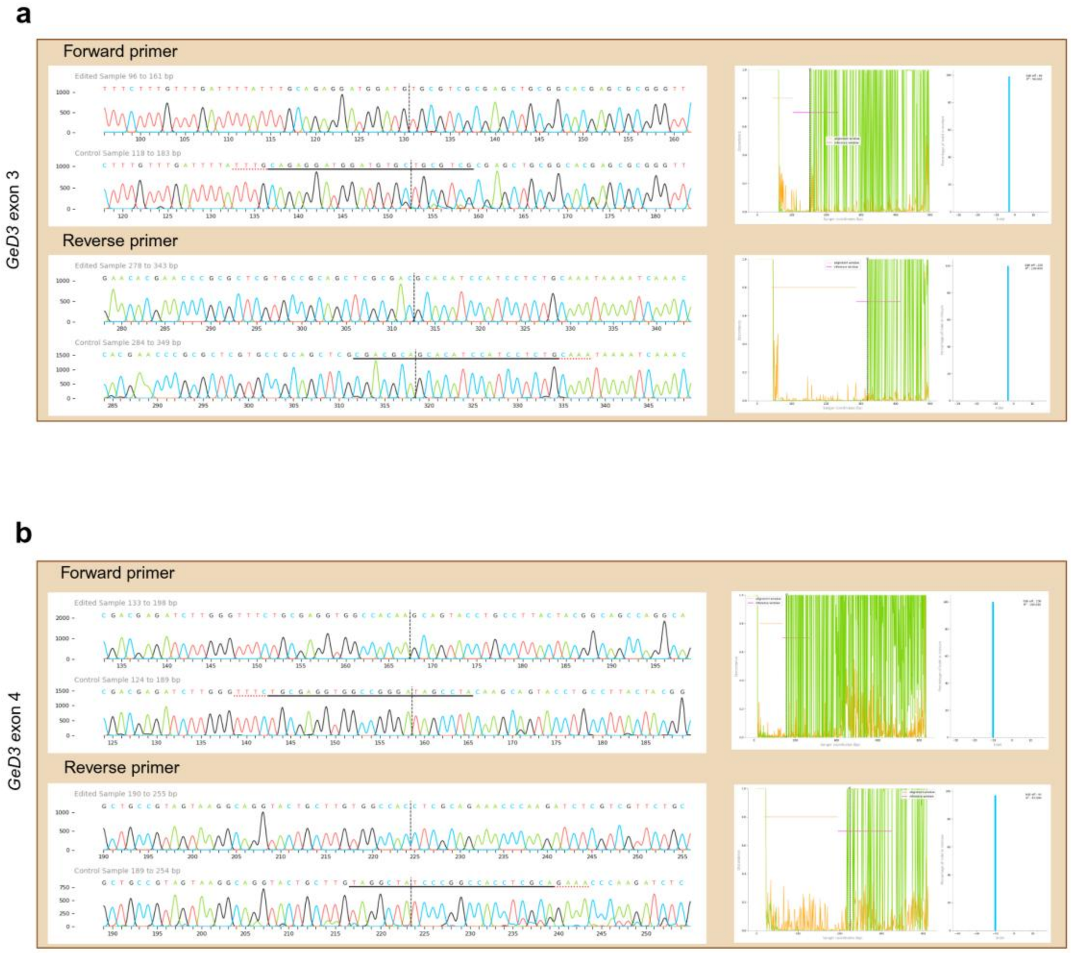


**Supplementary Fig. 10| Chromatogram alignment of *CKX2* gene between WT parent and GeD3.** The chromatogram alignment of +Ve and -Ve strand of the *CKX2* gene at exon 3 and exon 4 target site. The underline sequences represent the sgRNA of the particular strands. The mean of discord score before the first cut site and after the cut site is also shown. CRISPR edit at desired location and chromatogram alignment were checked and analyzed in BioEdit and ICE CRISPR analysis tool provided by the SYNTHEGO (https://www.synthego.com/).

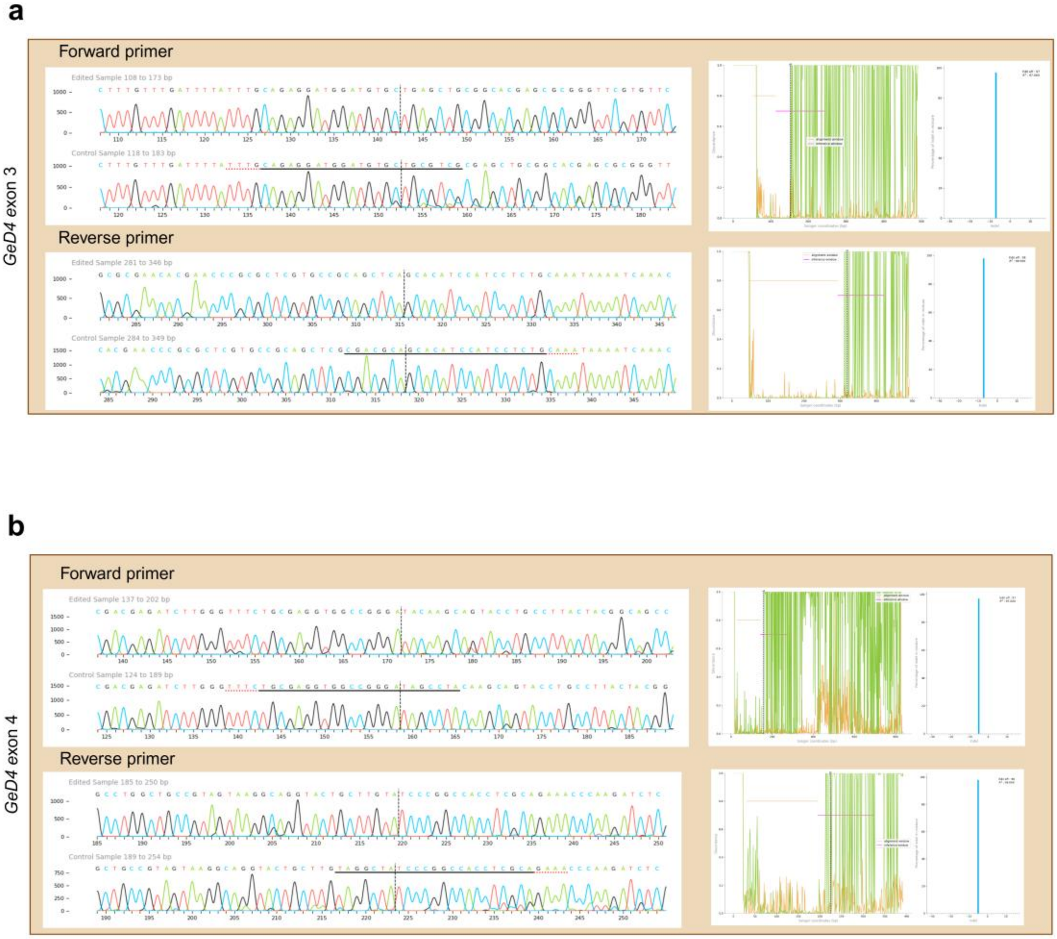


**Supplementary Fig. 11| Chromatogram alignment of *CKX2* gene between WT parent and GeD4.** The chromatogram alignment of +Ve and -Ve strand of the *CKX2* gene at exon 3 and exon 4 target site. The underline sequences represent the sgRNA of the particular strands. The mean of discord score before the first cut site and after the cut site is also shown. CRISPR edit at desired location and chromatogram alignment were checked and analyzed in BioEdit and ICE CRISPR analysis tool provided by the SYNTHEGO (https://www.synthego.com/).

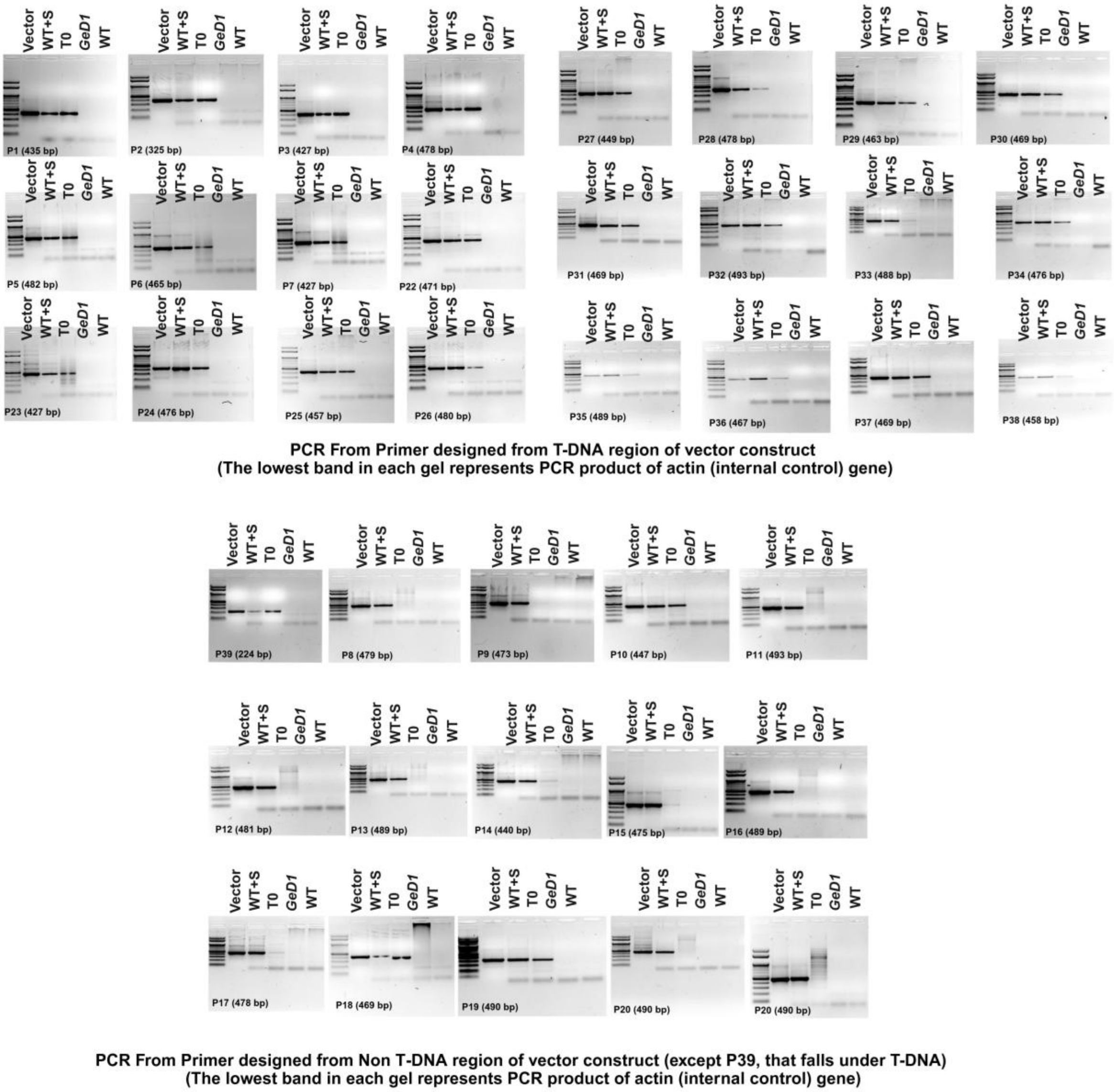


**Supplementary Fig. 12| Overlap PCR amplification corresponding to vector DNA in GeD1.** The amplifications using primer sets (P1-P7, P22-P38 and P39) spanning the T-DNA region of the transformation vector. The amplifications using primer sets (P8–P21) covering the non-T-DNA vector backbone (Supplementary Table 12). Expected amplicon sizes are indicated below each gel. Lanes correspond to: positive control plasmid DNA (vector), wild-type genomic DNA spiked with vector DNA to serve as a matrix control (WT+S), genomic DNA from the edited line (T0 stage plant ), transgene free edited line and wild-type parent genomic DNA negative control (WT). The uniformly amplified lowest band in all plant DNA lanes represents the endogenous *actin* internal control gene. Overlap PCR assays conducted on GeD2, GeD3, and GeD4 also produced similar results.

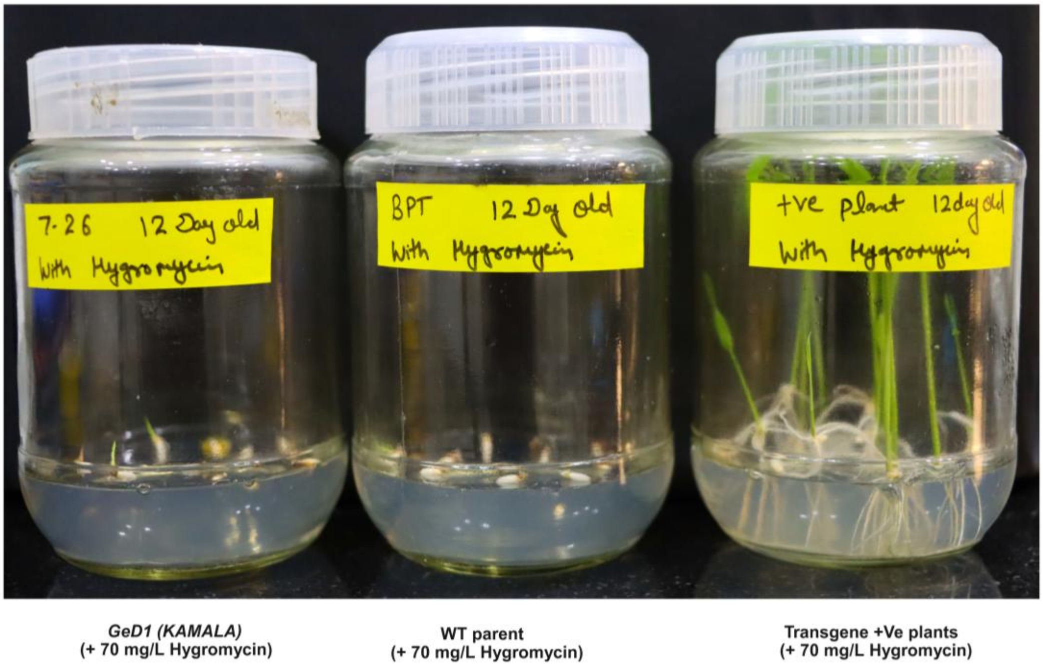


**Supplementary Fig. 13| Antibiotic sensitivity assay of GeD1 *(KAMALA)* line.** The photograph shows 12-day-old seedlings cultured on selection media supplemented with 70 mg/L hygromycin. Both the edited *KAMALA* line (left) and the unedited WT parent (middle) exhibit complete growth inhibition and failure to thrive. In contrast, the transgenic positive control (T0 seeds of *KAMALA*) demonstrates robust shoot and root development under identical selection pressure. Antibiotic sensitivity assays conducted on GeD2, GeD3, and GeD4 also produced similar results..

**a**

| Metric | KAMALA |
|---|---|
| Total reads | 2,483,541 |
| Avg. read length | 10, 498 bp |
| Max. read length | 46, 496 bp |
| Avg. base quality | Q38.3 |
| Mean depth | ~63x |
| Coverage breadth at 30x | ~11.2 Gb |

| Line | Feature | Start position | End position | Number of reads |
|---|---|---|---|---|
| WT parent | Rice ubiquitin promoter (rUbiP) | 12138 | 13854 | 612 |
| *KAMALA* | Rice ubiquitin promoter (rUbiP) | 12138 | 13854 | 563 |

**b**

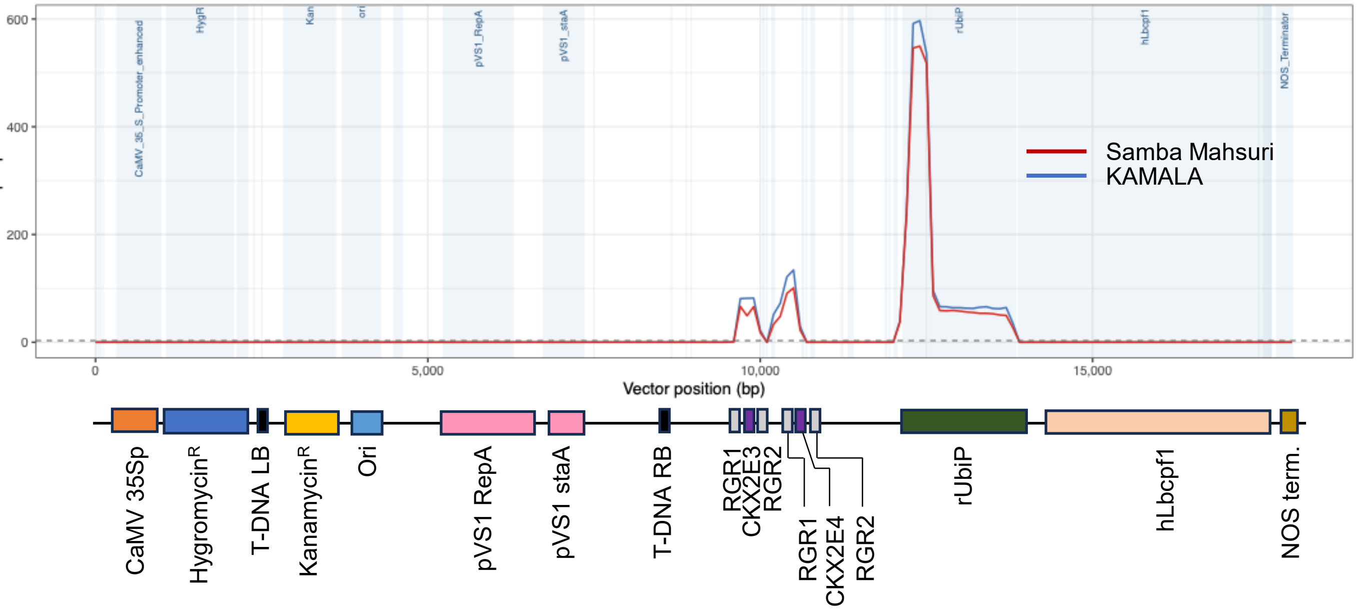


**Supplementary Fig. 14| Transgene analysis of *KAMALA* via long-read whole-genome sequencing.** **a,** WGS quality control of *KAMALA* for transgene analysis. **b,** Per-base sequencing depth across the transformation vector in *KAMALA* and wild-type Samba Mahsuri rice lines. Mean number of read depth per 100 bp bin in the genomes of *KAMALA* or WT parent (Samba Mahsuri) plotted against the length of the transformation vector (bp position of the vector). Shaded regions indicate annotated vector features.

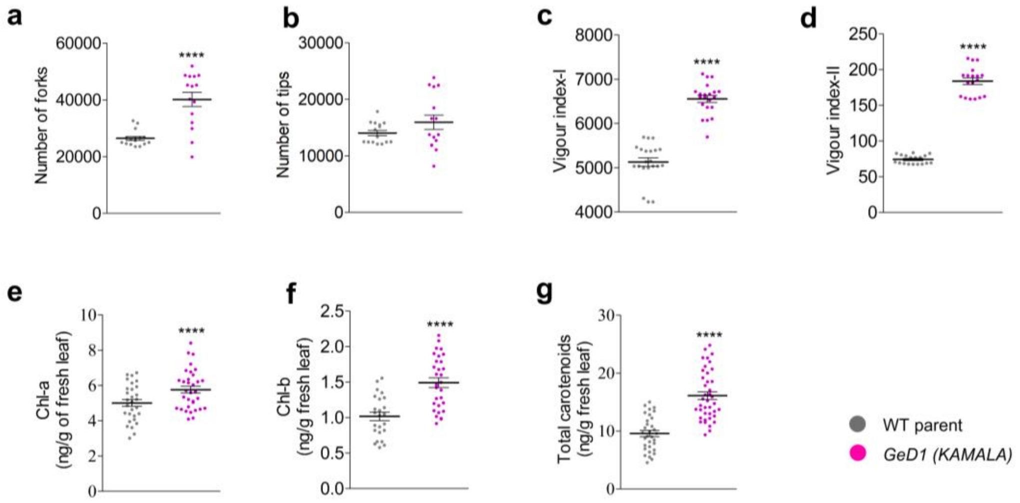


**Supplementary Fig. 15| Comparative analysis of GeD1 (*KAMALA)* and WT parent at vegetative stage. a, b,** quantification of root system architecture, including **a**, number of forks and **b**, number of tips. **c, d,** assessment of seedling performance based on **c**, Vigour Index-I and **d**, Vigour Index-II. **e–g,** concentrations of photosynthetic pigments in leaf tissues, including **e**, Chlorophyll-a (Chl-a), **f**, Chlorophyll-b (Chl-b), and **g**, total carotenoids. All plots show mean ± s.e.m.; sample sizes given in Methods. Statistical significance: one-way ANOVA with Tukey's HSD. Asterisks (*) indicate significance levels: * = $P$ value < 0.05, ** = $P$ value < 0.001, *** = $P$ value < 0.0001.

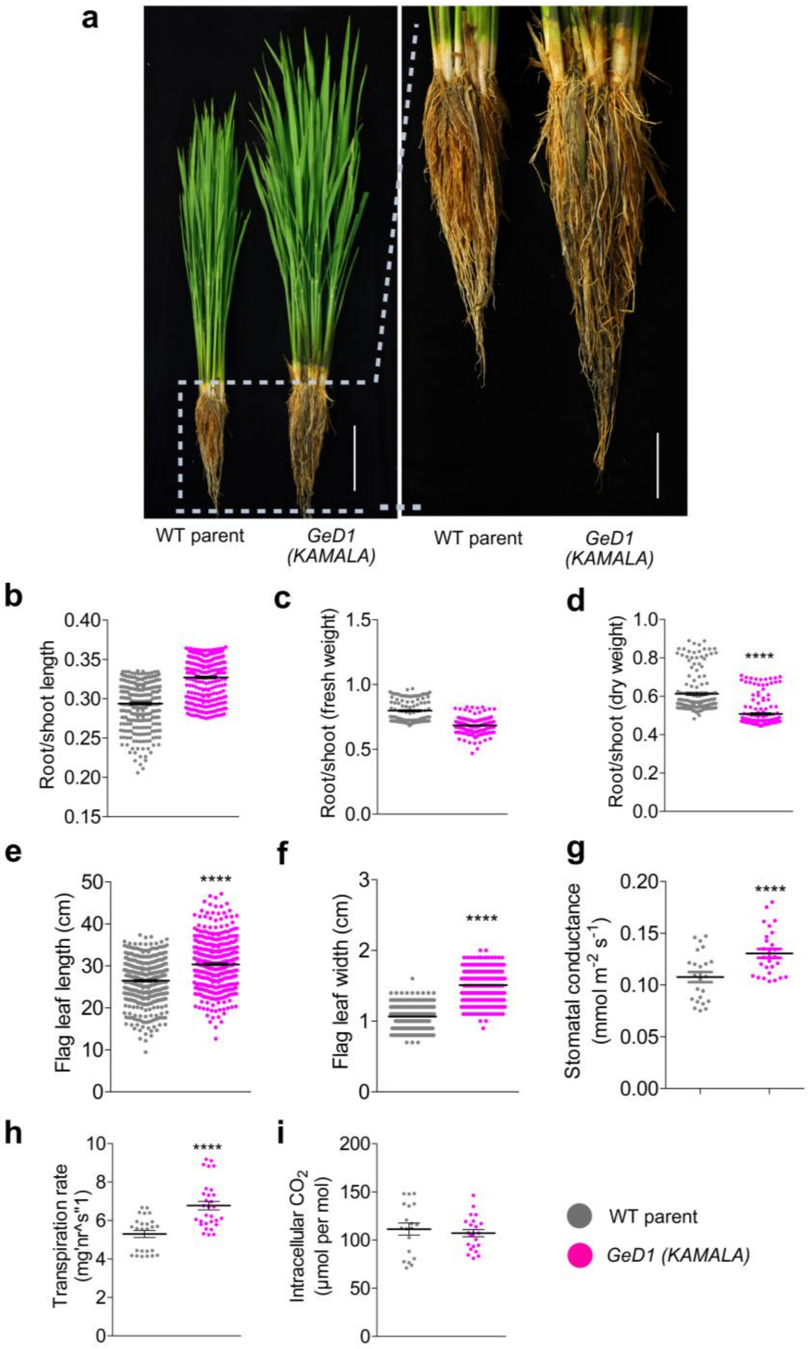


**Supplementary Fig. 16| Comparative analysis of GeD1 (*KAMALA)* compared to WT parent at reproductive stage. a,** plant stature and root system architecture are compared under field conditions (scale bars, 15 cm for plants; 5 cm for roots).**b–d,** ratios of root to shoot metrics including length (**b**), fresh weight (**c**), and dry weight (**d**). **e,f,** morphological measurements of the flag leaf, showing significantly increased flag leaf length (**e**) and width (**f**). **g–i,** gas exchange parameters measured under controlled conditions, including stomatal conductance (**g**), transpiration rate (**h**), and intracellular $CO_2$ concentration (**i**). All plots show mean ± s.e.m.; sample sizes in Methods. Statistical significance: one-way ANOVA with Tukey's HSD. Asterisks (*) indicate significance levels: * = *P* value < 0.05, ** = *P* value < 0.001, *** = *P* value < 0.0001.

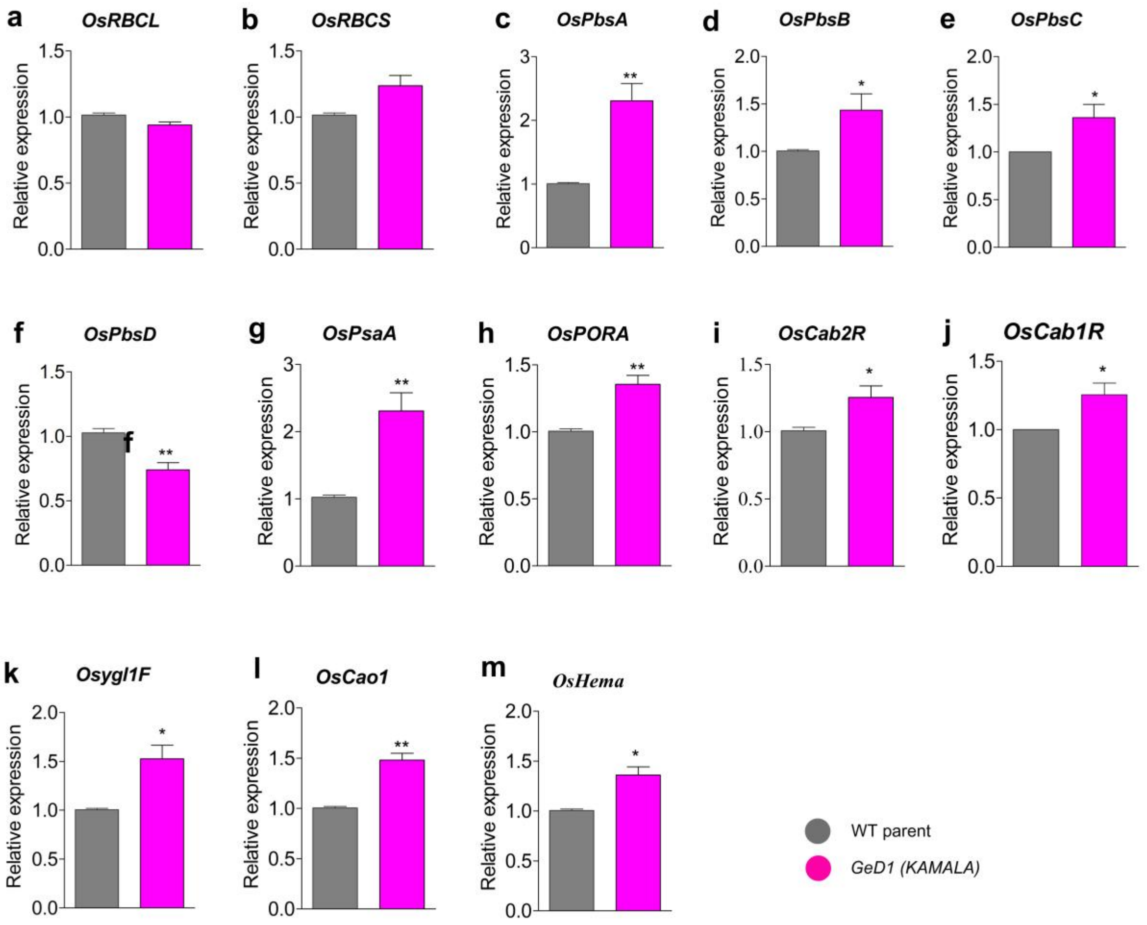


**Supplementary Fig. 17| Relative expression of the photosynthesis genes in the flag leaf of the GeD1 (*KAMALA)* and WT parent**. The chlorophyll biosynthesis, photosynthesis, and chloroplast development associated genes. **a**, *OsRBCL*, **b**, *OsRBCS*, **c**, *OsPbsA*, **d**, *OsPBsb*, **e**, *OsPbsc*, **f**, *OsPbsd*, **g**, *OsPsaA*, **h**, *OsPORA*, **i**, *OsCab2R*, **j**, *OsCab1R* and **k**, *Osygl1F*, **l**, *OsCao1* and **m**, *OsHema*. The qPCR primers of these genes were obtained from Gong et al., (2013). Rice Actin gene is used as the internal control. Data are presented as mean values + s.e.m., n = 3 biological replicates. Statistical significance was assessed using one-way ANOVA followed by Tukey's Honestly Significant Difference (HSD) test ($P < 0.05$). Asterisks (*) indicate significance levels: * = $P$ value < 0.05 and ** = $P$ value < 0.001.

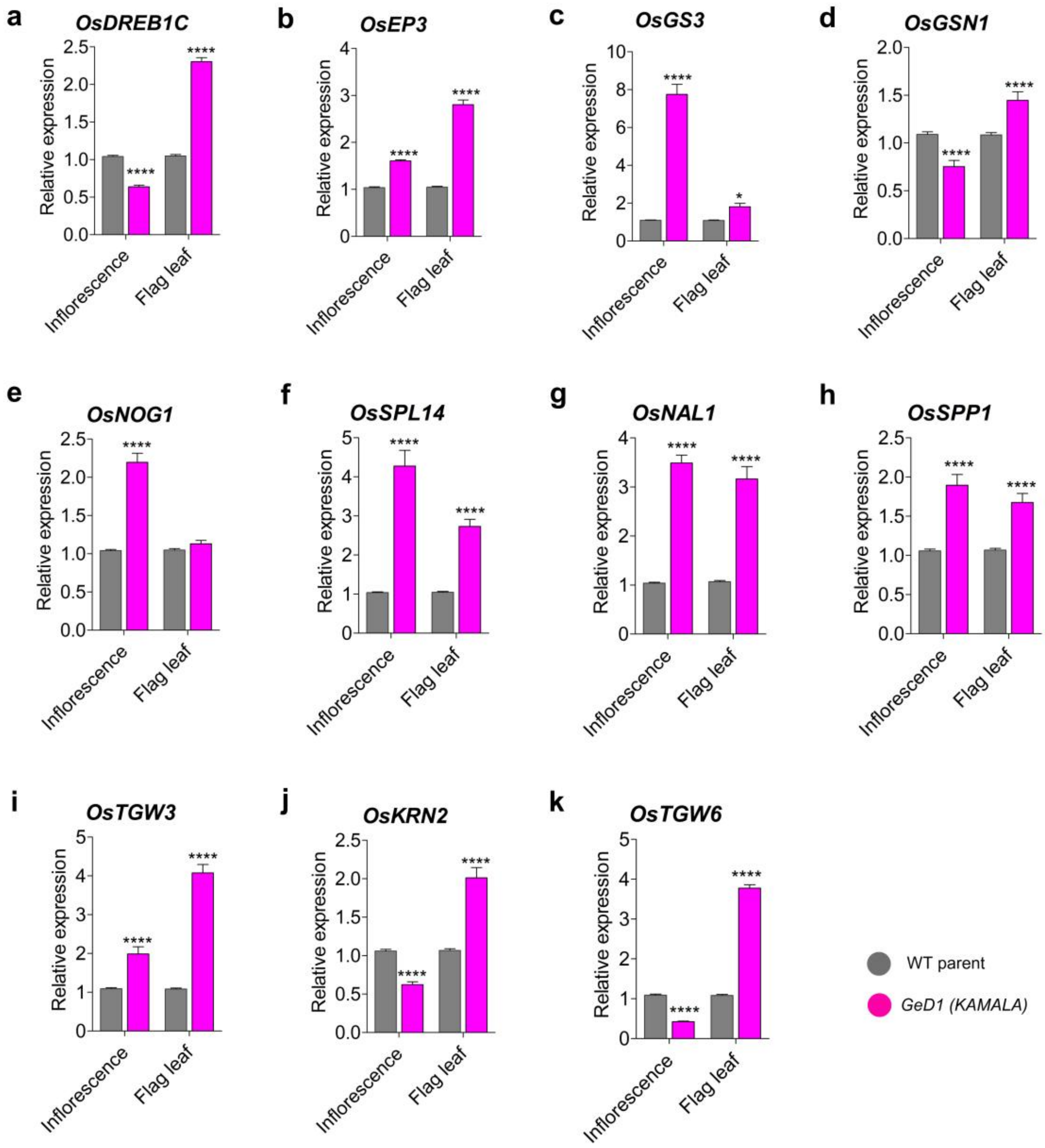


**Supplementary Fig. 18| Relative expression of yield trait related genes in inflorescence and flag leaf tissues of the GeD1 (*KAMALA)* and WT parent. a,** *OsDREB1C* (Dehydration-responsive element binding protein 1C ; *Os06g0127100*), **b,** *OsEP3* (Erect panicle 3; *Os02g0260200*), **c**, *OsGS3* (Grain size 3 ; *Os03g0407400*), **d**, *OsGSN1* (Grain size and number1 ; *Os05g0115800*), **e**, *OsNOG1* (Number of grains 1 ; *Os01g0752200*), **f**, *OsSPL14* (Squamosa promoter binding protein like-14 ; *Os08g0509600*), **g**, *OsNAL1* (Narrow leaf 1 ; *Os04g0615000*), **h**, *OsSPP1* (Signal peptide peptidase 1; *Os02g0117400*)**, i,** *OsTGW3* (Thousand grain weight 3 ; *Os03g0841800*)**, j,** *OsKRN2* (Kernel row number 2 ; *Os04g0568400*), and **k**, *OsTGW6* (Thousand grain weight 6 ; *Os06g0623700*). Rice Actin gene is used as the internal control. Data are presented as mean values + s.e.m., n = 3 biological replicates. Statistical significance was assessed using one-way ANOVA followed by Tukey's Honestly Significant Difference (HSD) test ($P < 0.05$). Asterisks (*) indicate significance levels: **** = $P$ value < 0.00001.

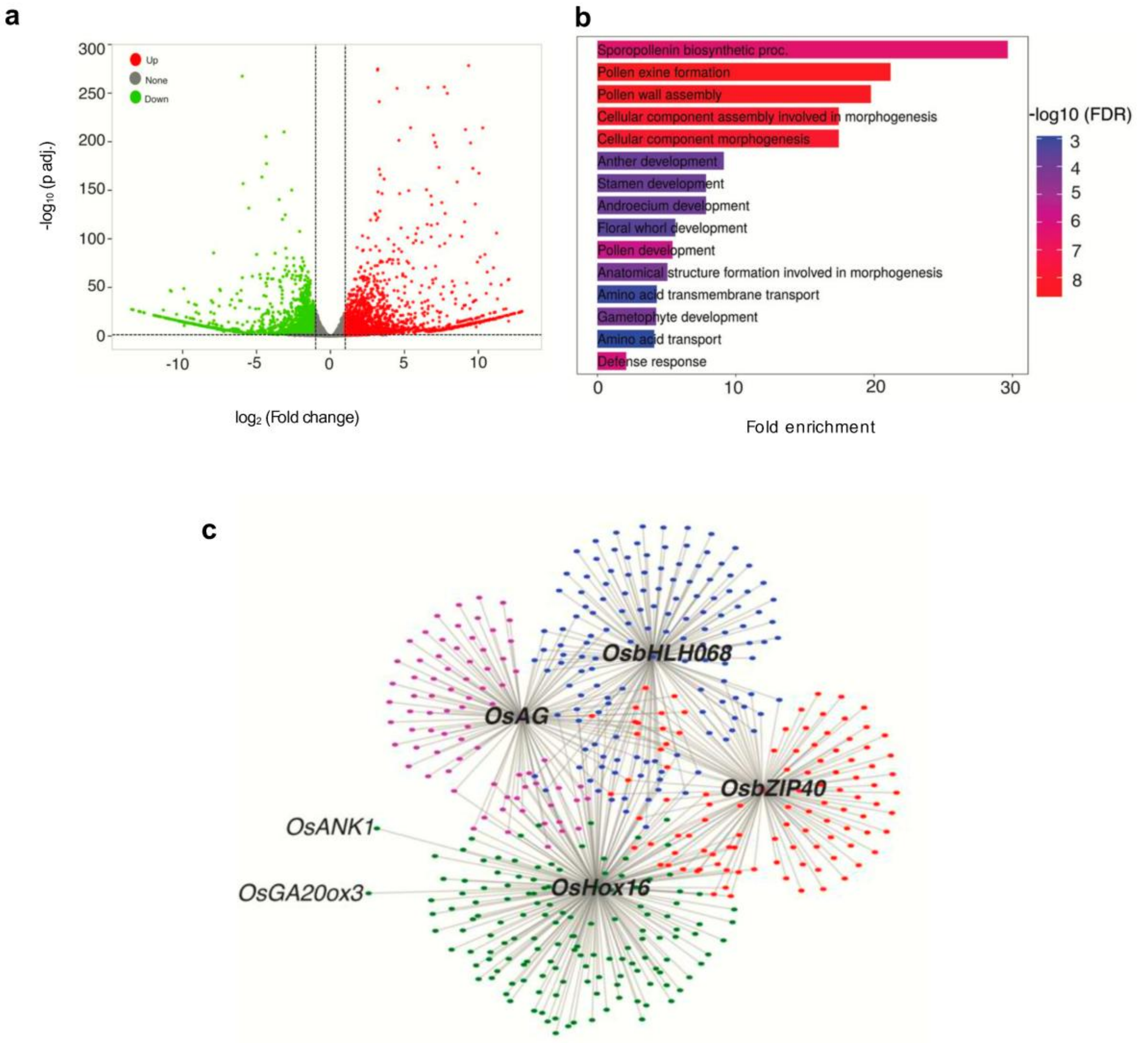


**Supplementary Fig. 19| RNA-seq analysis of differentially expressed genes (DEGs). a,** volcano plot of differentially expressed genes (DEGs) from inflorescence tissue. **b,** Gene Ontology (GO) enrichment analysis of upregulated DEGs. The x-axis displays the fold enrichment of the specific GO terms. Bar colours represent the statistical significance of the enrichment, denoted as -log10 (false discovery rate, FDR), ranging from blue (lower significance) to red (higher significance). The analysis highlights significantly enriched pathways involved in reproductive organ development, including sporopollenin biosynthesis, pollen exine formation, and anther development. **c,** Transcription factor (TF) regulatory networks showing interactions between source TFs and upregulated target DEGs. The network highlights key central source TFs, including *OsHox16*, *OsAG*, *OsbHLH068*, and *OsbZIP40*, alongside their associated downstream gene clusters. Nodes represent individual genes, and edges denote predicted regulatory interactions. Specific upregulated downstream targets of interest, such as *OsANK1* and *OsGA20ox3*, are explicitly highlighted within the *OsHox16* sub-network.

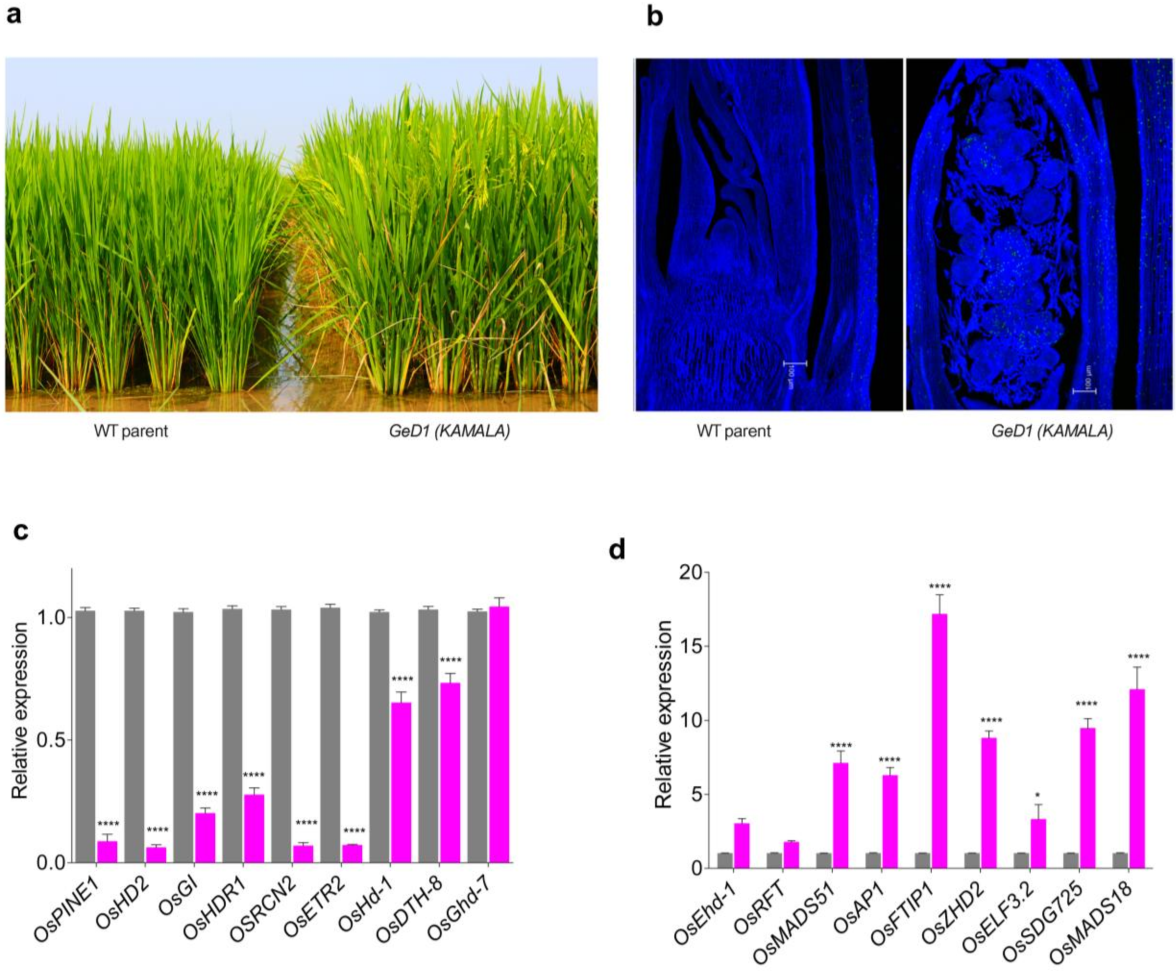


**Supplementary Fig. 20| GeD1 (*KAMALA)* flowers 20-25 days earlier and matures faster than the WT parent in both Rabi and Kharif seasons.** **a**, field-grown plants of WT parent (left panel) and *KAMALA* (right panel) at ICAR-IIRR, showing early flowering phenotype in *KAMALA*. **b,** EdU-stained actively dividing cells images of shoot apices with enhanced cell proliferation in GeD1 (*KAMALA*) (scale bar, 100 µm). **c-d**, relative expression of negative regulators (**c**) and positive regulators (**d**) of flowering, measured by RT-qPCR. A schematic diagram illustrating the rice flowering pathway and role of these genes is explained in Supplementary Fig. 20. Bar graph data represent mean values from three biological replicates ± s.e.m. Statistical significance between *KAMALA* and WT was evaluated using one-way ANOVA followed by Tukey's HSD test ($P < 0.05$). Asterisks (*) indicate significance levels: * = $P$ value < 0.05 and **** = $P$ value < 0.00001.

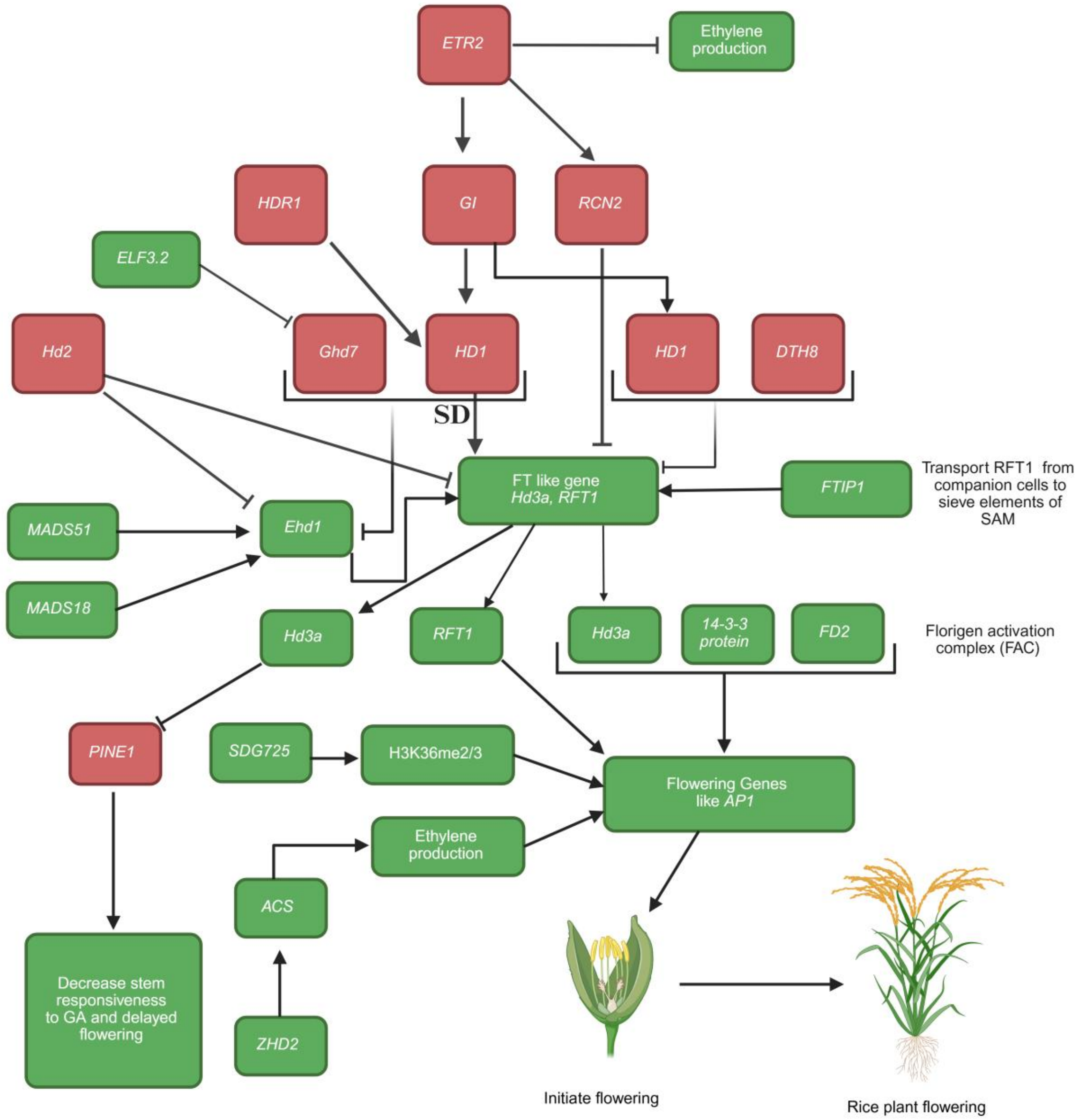


**Supplementary Fig. 21| Flow diagrammed of responsible genes in the early flowering in rice.** The flowering associated genes *OsFTIP1, OsMADS18, OsMADS51, OsSDG725, OsZHD2, OsAP1, and OsELF3.2* showed up-regulation *in KAMALA*. The transportation of florigen protein from companion cells to sieve elements of the shoot apical meristem (SAM) is dependent on the FTIP1 (Florigen Transport Inhibitor Protein 1) [Liu et al., 2012]. MADS51 and MADS18 are positive regulators for Ehd1 gene [Yin et al., 2019] which directly promotes flowering by inducing expression of *Hd3a* and *RFT*1 genes [Kim et al., 2007]. The florigen genes *Hd3a* and *RFT1* are considered as the central hub for flowering initiation [Komiya et al., 2008]. Downstream, 14-3-3 protein and FD2 form a complex with Hd3a known as Florigen Activation Complex (FAC), that initiate flowering through flowering gene AP1 [Taoka et al., 2011]. Whereas RFT1 independently promote the expression of AP1 [Komiya et al., 2008]. The SDG725 promote methylation at H3K36me2/3 of AP1 gene that leads to flowering initiation [Sui et al., 2012 & 2013]. The *OsPINE1, OsHD2, OsGI, OsHDR1, OSRCN2, OsETR2, OsHd-1, and OsDTH-8* showed down-regulation in *KAMALA*. Independent Hd3a protein promote the flowering via inhibiting the PINE1, which decreases responsiveness towards GA that leads to delay flowering [Taoka et al., 2011]. Ethylene homeostasis is maintained positively by the ZHD2 [Yoon et al., 2020] through ACS expression and negatively by ETR2 expression [Wuriyanghan et al., 2009]. ETR2 promotes the expression of GI (GIGANTEA) and RCN2. RCN2 directly interacts with florigen genes Hd3a and RFT1 and inhibits their expression [Zhou et al., 2021]. The red and green boxes genes represent the negative and positive regulator for the flowering.

**a**

| PCR Component | Volume (1X) |
|---|---|
| Nuclease free water | 7 µl |
| 10X Taq Buffer | 1 µl |
| 2.5 mM dNTPs | 0.5 µl |
| 10 µM Forward Primer (CTCACTTTATCCTTGCATGAC) | 0.4 µl |
| 10 µM Reverse Primer (TGAGGGGTCGTCATTTTG) | 0.4 µl |
| Taq Polymerase | 1 unit |
| DNA | ~200 ng |
| Total | 10 µl |

**b**

**PCR Conditions**

- 95°C - 3 min.
- 95°C - 30 sec.
- 50.5°C - 3 min
- 72°C - 1 min
- 72°C - 5 min

35 cycles

**c**

**Gel Electrophoresis conditions**

- 2.5% agarose used for gel preparation in 1% TAE buffer
- Run at 110 volts for minimum 4 hours
- Visualized the gel in Gel-documentation

**d**

**CAPS Marker**

- Used same PCR product of co-dominant marker
- Restriction digestion with NruI (Prefer NEB make enzyme for better performance) as guided with leaflet
- Run 1% agarose gel after restriction
- Visualized the gel in Gel-documentation unit

**e**

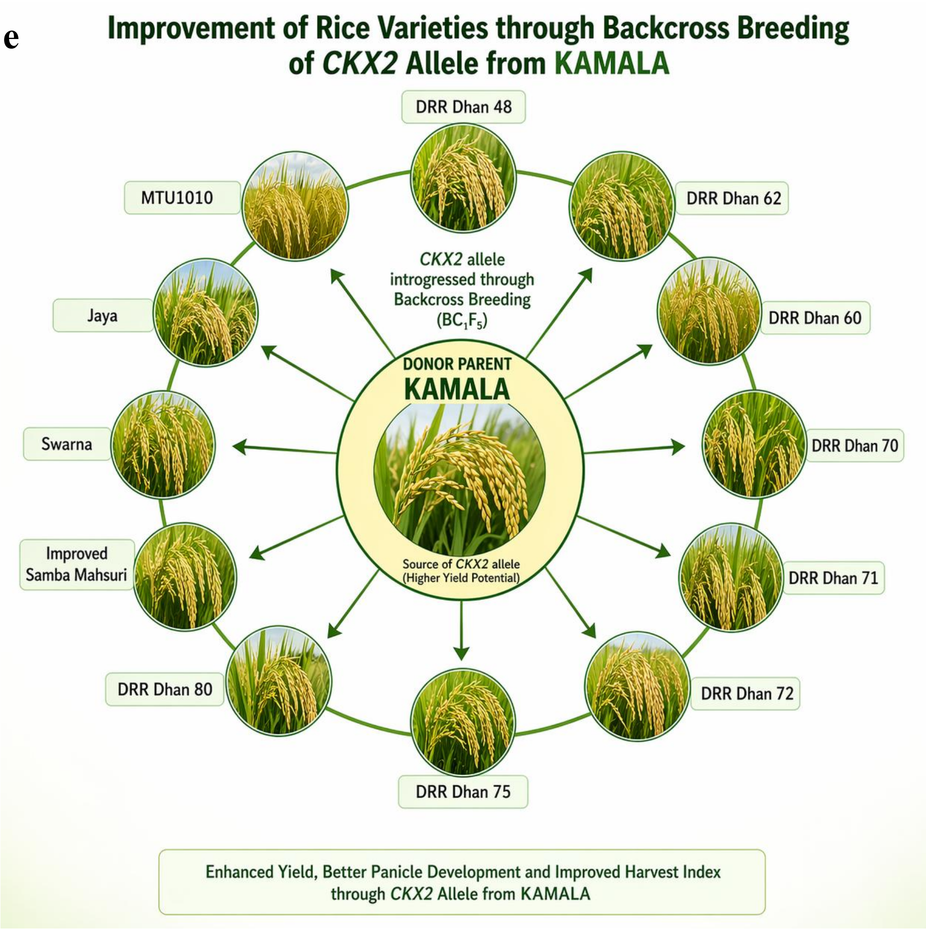


**Supplementary Fig. 22| Protocol for the identification of *KAMALA-CKX2* allele using PCR and CAPS markers and its introgression into popular rice varieties**. **a**, specified PCR components and primers, **b**, PCR conditions, **c**, gel electrophoresis conditions, **d**, protocol for use as CAPS marker, and **e**, ongoing efforts of marker assisted introgression of *KAMALA*-CKX2 into popular rice varieties (Image: created by ChatGPT).

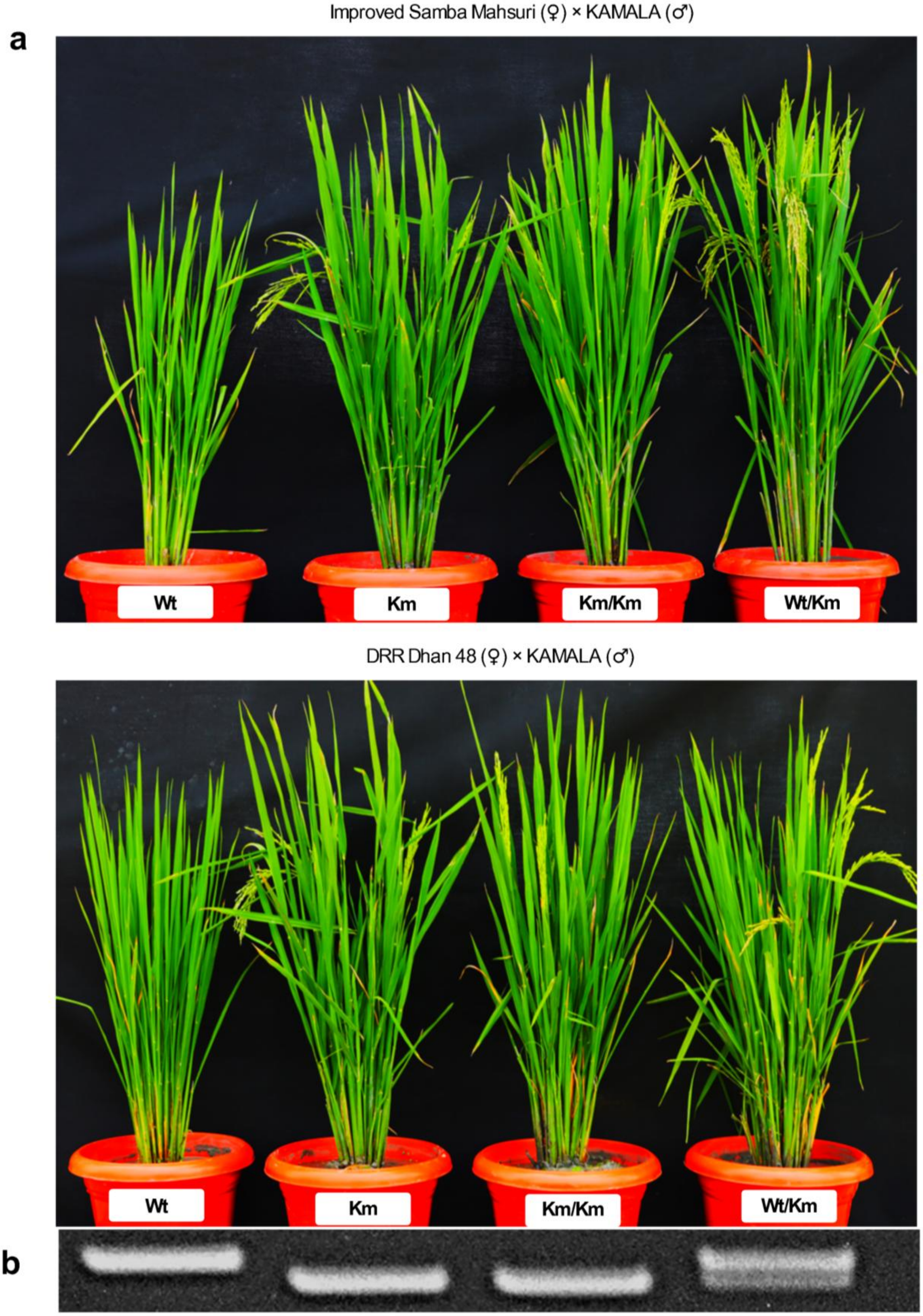


**Supplementary Fig. 23| Genetic analysis of *KAMALA-CKX2* mutation. a,** Improved Samba Mahsuri and DRR Dhan 48 (Wt) were crossed with *KAMALA* (Km). The homozygous (Km/Km) and heterozygous plants (Wt/Km) were analyzed in BC1F5 generation. **b,** Agarose gel showing amplification of Wt and *KAMALA* alleles of CKX2 using the DNA markers developed in this study. Heterozygous plants exhibit amplification of both WT and *KAMALA*-specific bands.

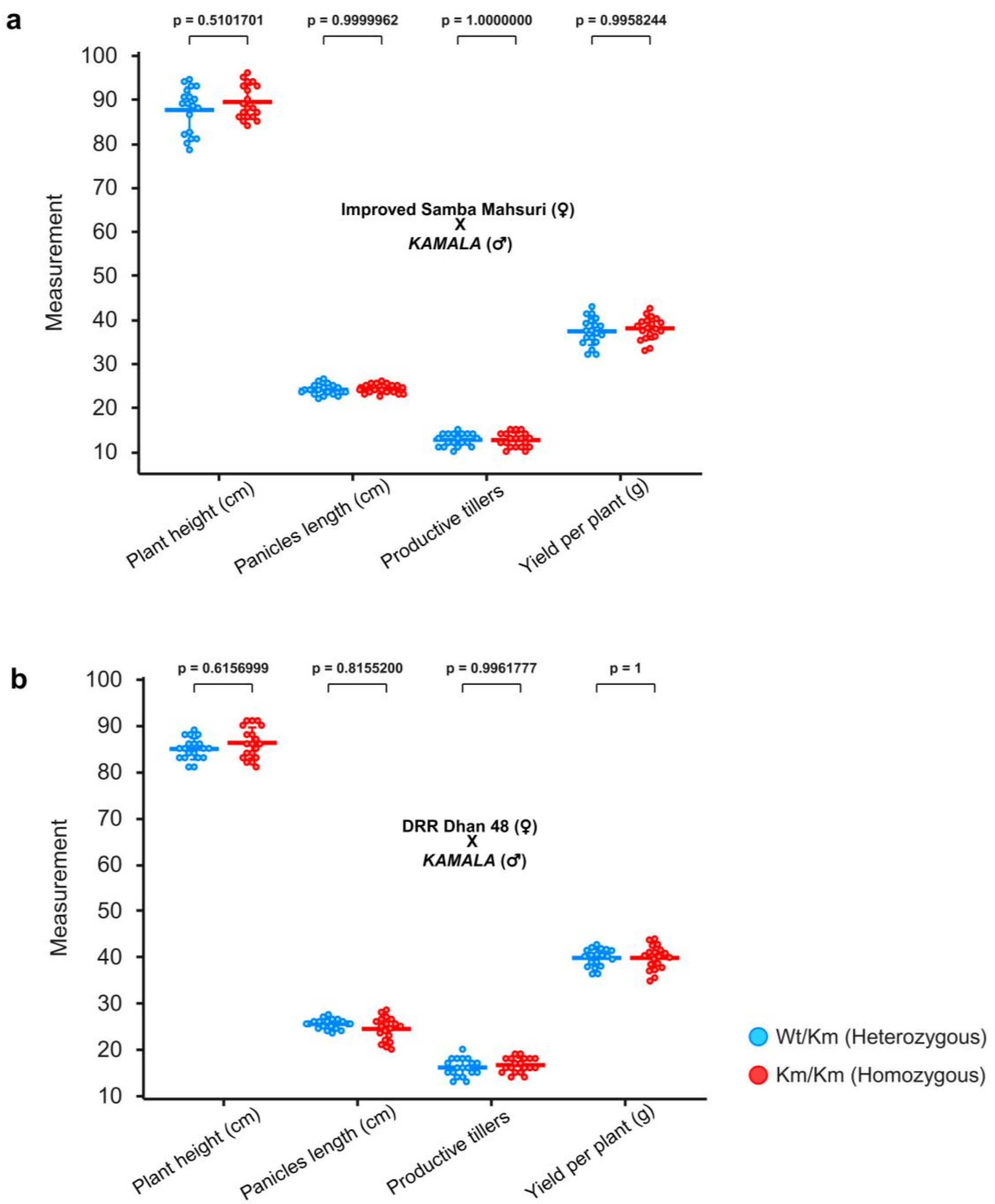


**Supplementary Fig. 24| Phenotypic evaluation of agronomic traits of homozygous and heterozygous plants (for *KAMALA CKX2* allele) shown in Supplementary Fig. 22.** Comparison of plant height (cm), length of panicles (cm), productive tillers number, and yield per plant (g) of heterozygous (grey circles) and homozygous (pink squares) plants of (a) Improved Samba Mahsuri (♀) × *KAMALA* (♂) and (b) DRR Dhan 48 (♀) × *KAMALA* (♂). Data points represent individual plants, with horizontal lines indicating mean ± sem. Statistical significance was determined using Two-way ANOVA followed by Bonferroni's multiple comparisons test (sample size-19).

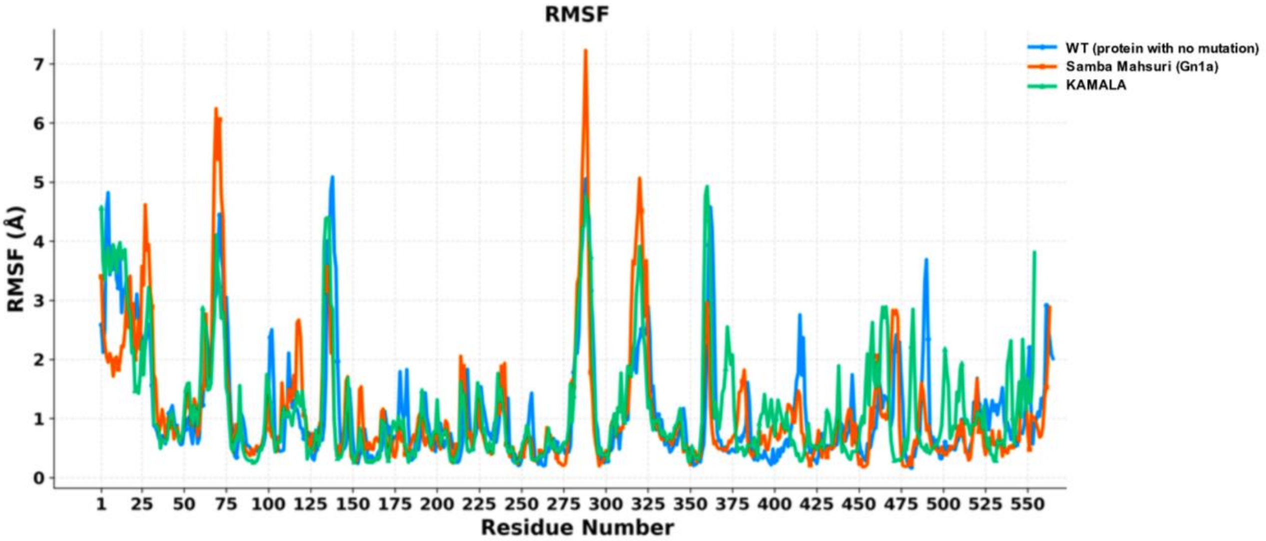


**Supplementary Fig. 25| Protein flexibility analysis and three-dimensional structural models of CKX2.** Protein structural flexibility of CKX2 was analyzed for the WT (protein with no mutations), Samba Mahsuri (Gn1a; carrying a natural 6 bp deletion in exon 1), and *KAMALA* using CABS-flex 3.0. The root-mean-square fluctuation (RMSF) for each residue is plotted on the Y-axis, where RMSF represents the average deviation of amino acid residues from their equilibrium positions, explaining the intrinsic flexibility of protein.

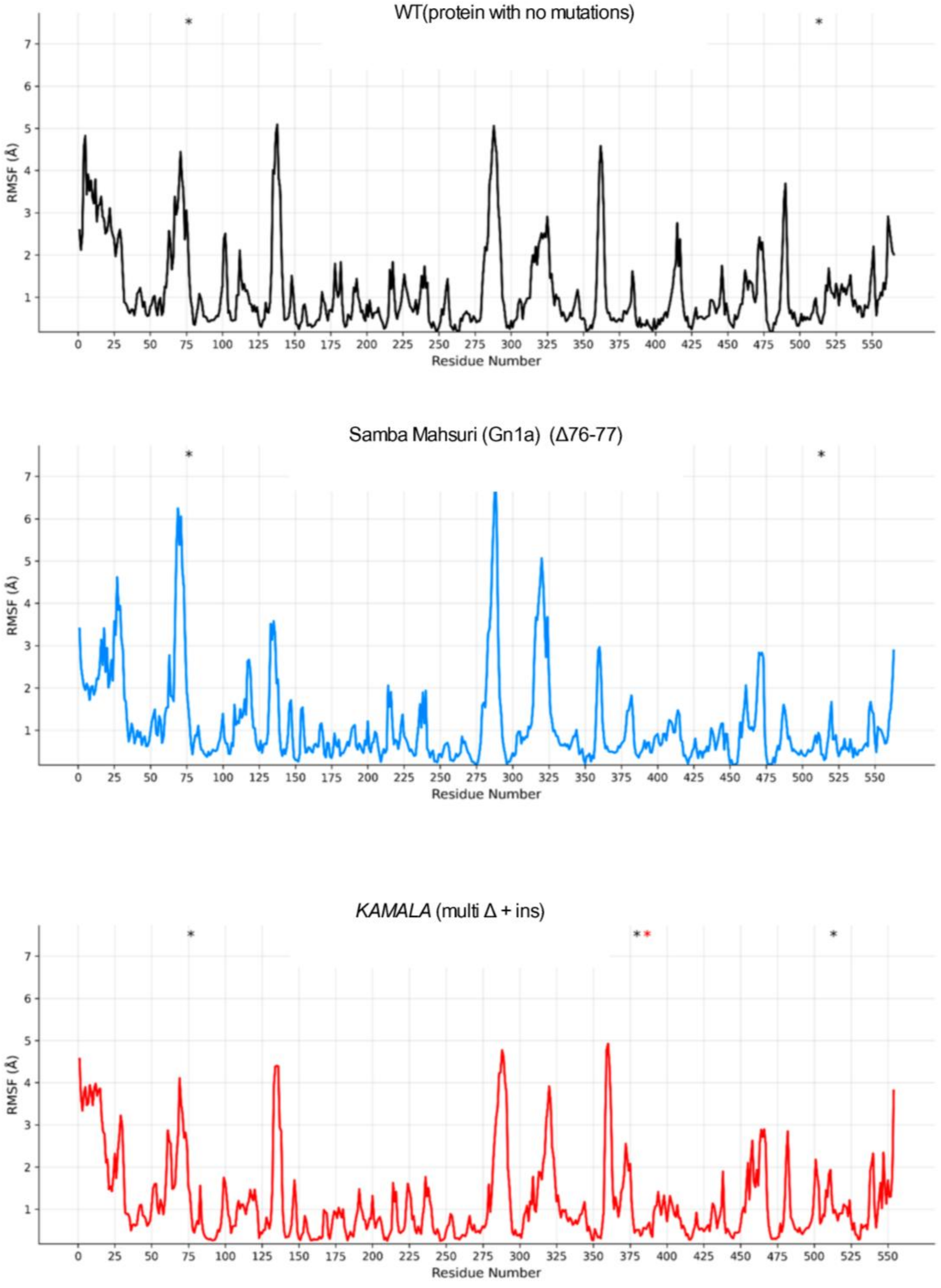


**Supplementary Fig. 26|** Protein structural flexibility of CKX2 analyzed for the WT (protein with no mutations), Samba Mahsuri (Gn1a; carrying a natural 6 bp deletion in exon 1), and *KAMALA* using CABS-flex 3.0. The root-mean-square fluctuation (RMSF) for each residue is plotted on the Y-axis, where RMSF represents the average deviation of amino acid residues from their equilibrium positions, explaining the intrinsic flexibility of protein.

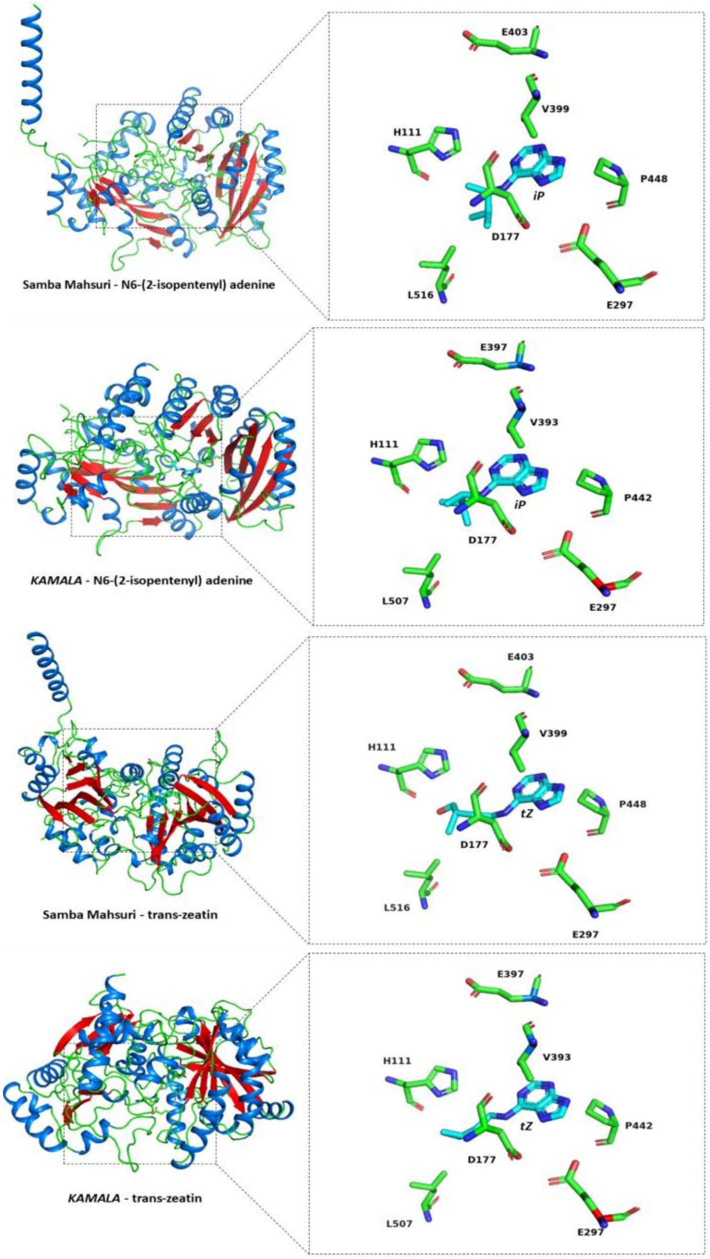


| Confidence metrics | SM-Tz | *KAMALA*-Tz | SM-iP | *KAMALA*-iP |
|---|---|---|---|---|
| **pTM** | 0.92 | 0.93 | 0.92 | 0.92 |
| **pLDDT** | 0.87 | 0.85 | 0.86 | 0.84 |
| **pIC50 Binding Affinity (kcal/mol)** | -5.88 | -5.89 | -6.49 | -6.45 |
| **Binding Probability** | 0.56 | 0.53 | 0.52 | 0.43 |

**Supplementary Fig. 27| Prediction of CKX2-substrate (cytokinin) structural complex and binding affinities.** Compared to Samba Mahsuri (SM), the *KAMALA* variant exhibits diminished cytokinin binding affinity. Boltz-2 (Passaro et al 2025), an AI driven structural biology foundation model was used to analyze the structural complexes of SM and *KAMALA* CKX2 with cytokinin substrate-iP (N6-(2-isopentenyl) adenine) and tZ (trans-zeatin). The binding affinity of iP and tZ were predicted considering the protein structure, and FAD and cytokinin binding residues of experimentally characterized maize CKX (Kopečný et al 2016). All the four structural complexes (SM-iP, *KAMALA*-iP, SM-tZ and *KAMALA*-tZ) were predicted with high confidence (pTM: 0.92–0.93). *KAMALA* shows lower binding probability (0.43) with N6-(2-isopentenyl) adenine.

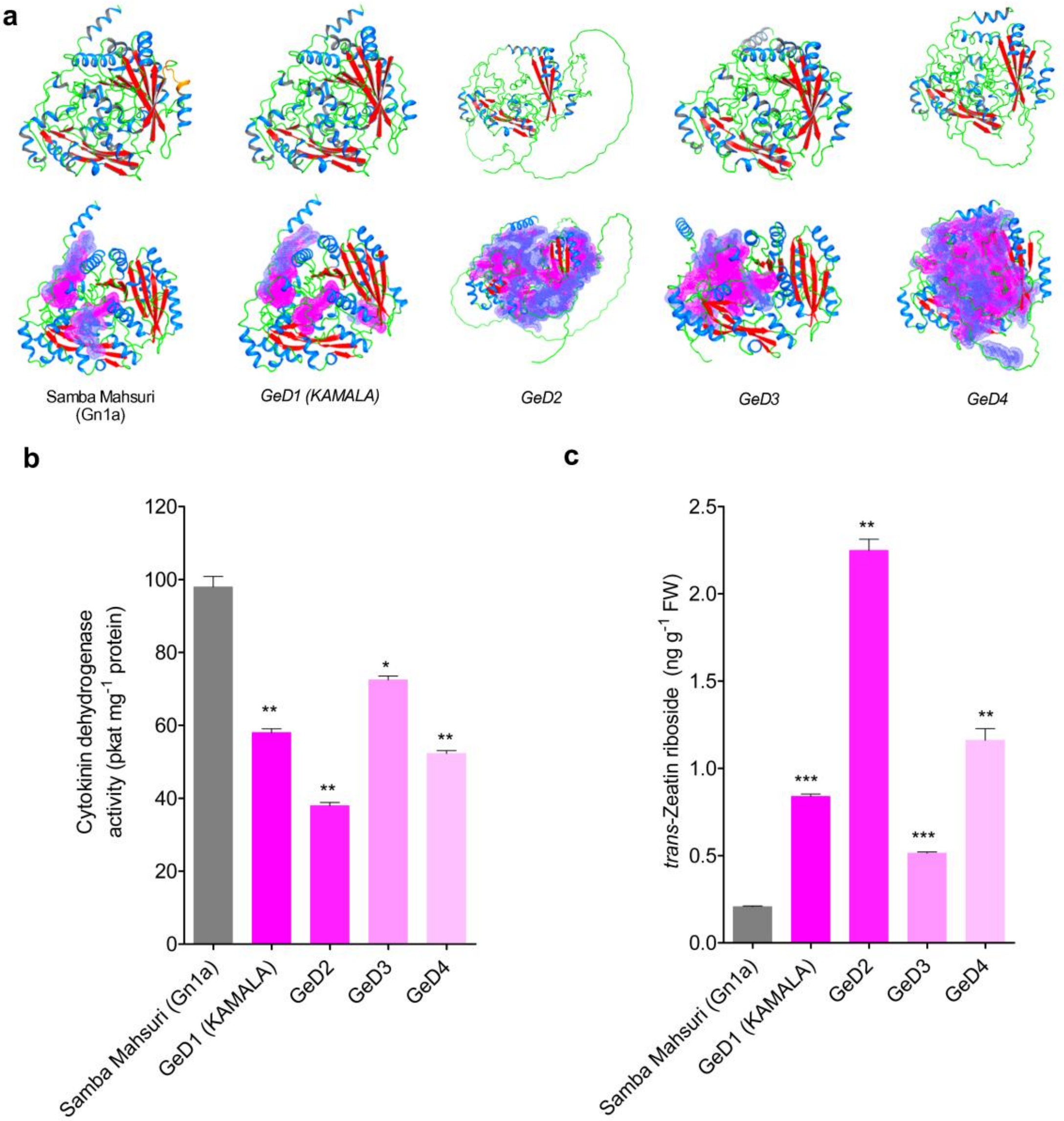


**Supplementary Fig. 28| CKX2 structure prediction, enzyme activity and cytokinin levels in genome edited lines.** **a,** predicted three-dimensional (3D) structural models of the CKX2 enzyme and analysis of active site cavities. 3D structural models of the protein from Samba Mahsuri and four gene-edited lines-GeD1 *(KAMALA),* GeD2, GeD3, and GeD4, were predicted using AlphaFold3. The upper row illustrates the predicted structure. All models are colored by secondary structure: alpha-helices are shown in marine, beta-strands in red, and loops/turns are shown in green. The truncated or distorted structures in the gene-edited lines indicate premature termination or significant frame-shift mutations affecting proper protein folding. The bottom row highlights the predicted active-site cavities computed with CavitOmiX. The top cavities with the largest volume are shown as a magenta/purple density in the superposition models, emphasizing the dramatic loss of the functional active site structure and predicted cavity volume in the functional knockout lines. **b,** CKX enzyme activity measured in total protein from the inflorescence tissue (using the same sample as in c). **c,** Concentration of trans-zeatin riboside (tZR) in inflorescence tissue. All data represent the mean along with the standard error of the mean (s.e.m.) from at least three independent replicates. Statistical significance compared to the WT was determined by a two-tailed Student's t-test. Asterisks (*) indicate significance levels; * = $P$ value < 0.05, ** = $P$ value < 0.01, and *** = $P$ value < 0.001.

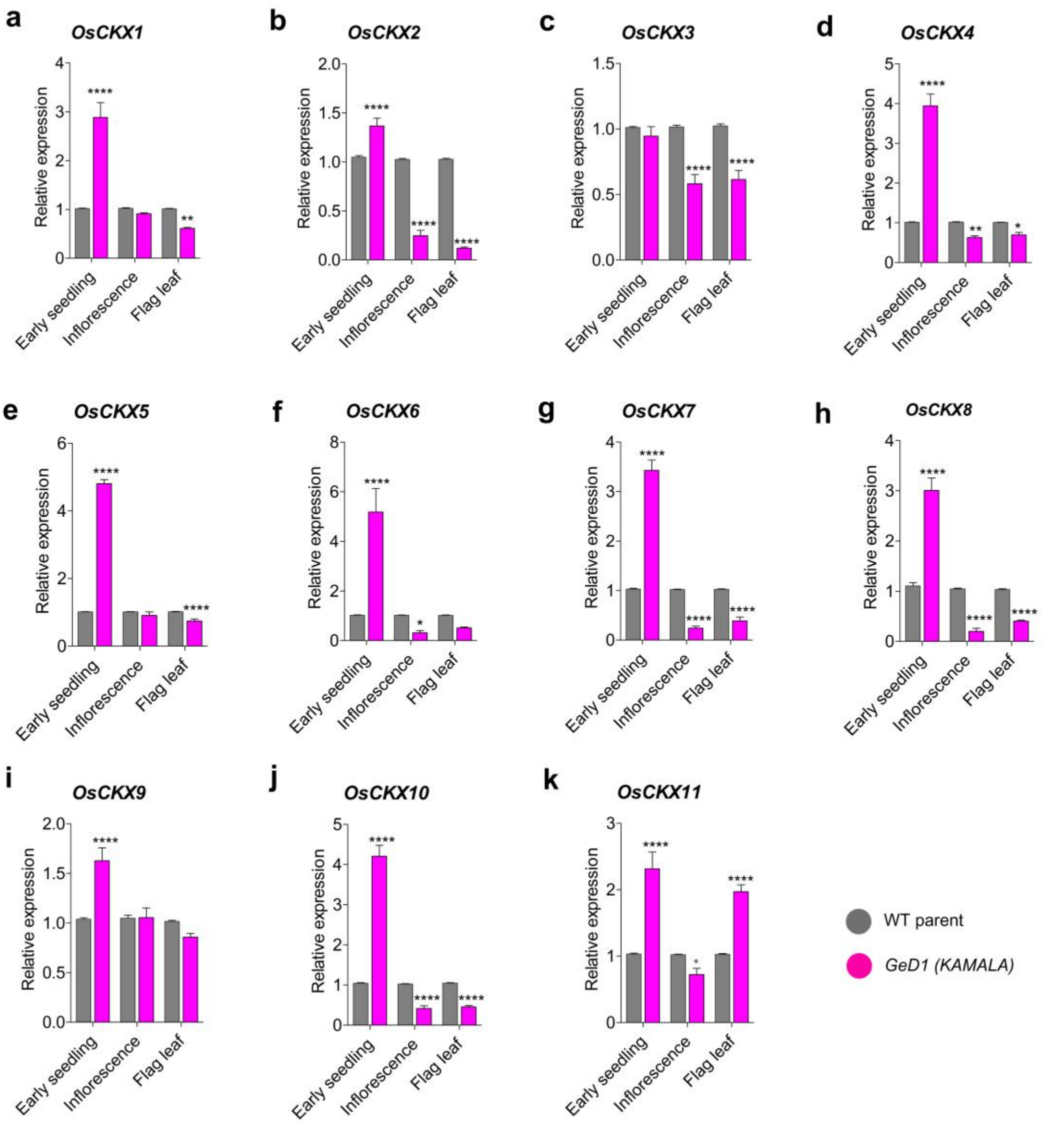


**Supplementary Fig. 29| Relative expression of the cytokinin oxidase/dehydrogenase (*OsCKX*) family genes in the early seedling (21d old), inflorescence and flag leaf of the GeD1 (*KAMALA)* and WT parent**. **a**, *OsCKX1* (*Os01g0187600*), **b**, *OsCKX2* (*Os01g0197700*), **c**, *OsCKX3* (*Os10g0483500*), **d**, *OsCKX4* (*Os01g0940000*), **e**, *OsCKX5* (*Os01g0775400*), **f**, *OsCKX6* (*Os02g0220000*), **g**, *OsCKX7* (*Os02g0220100*), **h**, *OsCKX8* (*Os04g0523500*), **i**, *OsCKX9* (*Os05g0374200*), **j**, *OsCKX10* (*Os06g0572300*) and **k**, *OsCKX11* (*Os08g0460600*). Rice Actin gene is used as the internal control. Data are presented as mean values + s.e.m., n = 3 biological replicates. Statistical significance was assessed using one-way ANOVA followed by Tukey's Honestly Significant Difference (HSD) test ($P < 0.05$). Asterisks (*) indicate significance levels: * = $P$ value < 0.05; ** = $P$ value < 0.001 and **** = $P$ value < 0.00001.

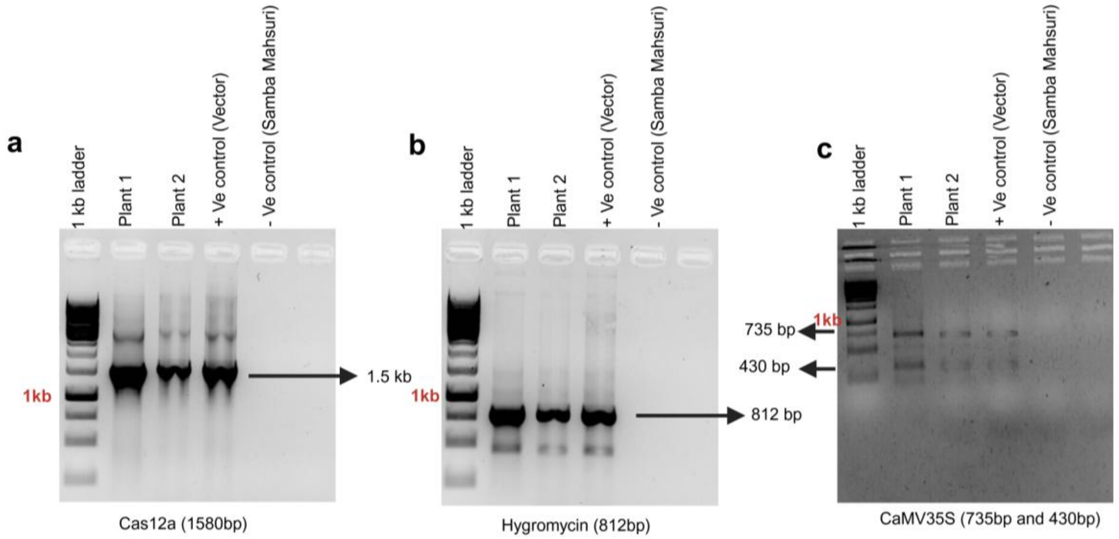


**Supplementary Fig. 30| Molecular confirmation of T-DNA integration in transgenic rice lines. a-c**, A representative gel image showing PCR-based validation T-DNA integration in two independent transformed lines (Plant 1 and Plant 2). Gel electrophoresis panels display the specific amplification of the Cas12a nuclease gene (expected size 1.5 Kb) (**a**), Hygromycin B phosphotransferase gene (812 bp) (**b**), and the CaMV35S promoter elements (735 bp and 430 bp) (**c**). For all panels, lanes correspond to: 1 kb DNA ladder, PCR from genomic DNA of transgenic plants- Plant 1 and Plant 2, plasmid DNA (+Ve control), and genomic DNA of non-transformed WT parent (-Ve control) . The presence of PCR amplification of T-DNA genes in Plant 1 and Plant 2 confirms the successful genetic transformation and integration of the Cas12a editing gene casette into the host genome.

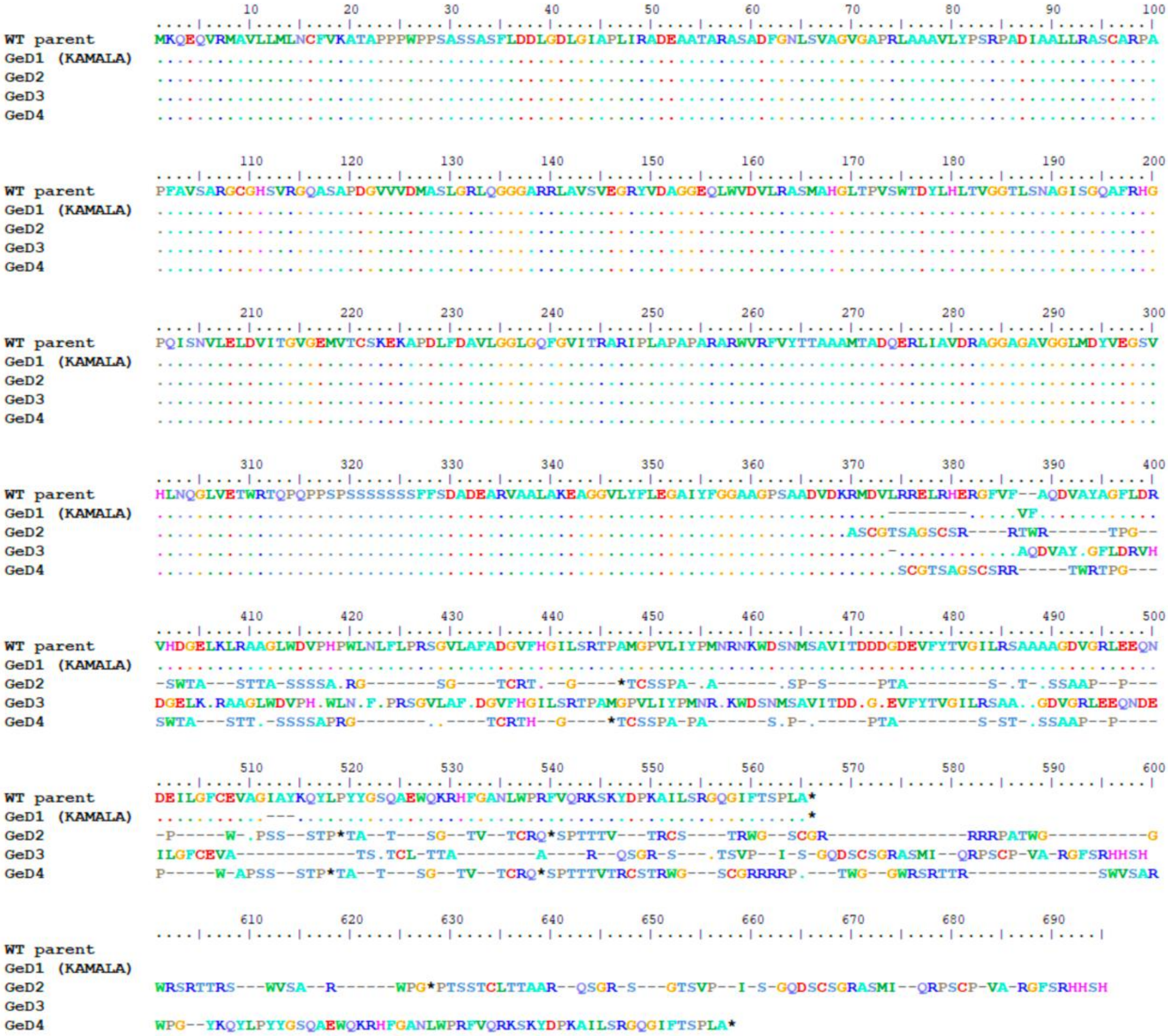


**Supplementary Fig. 31a| Amino acid sequence alignment of WT parent and GeD variants.** Multiple sequence alignment comparing the WT parent protein sequence with four derivative variants: GeD1 (*KAMALA*), GeD2, GeD3, and GeD4. Residues that are identical to the WT parent sequence are denoted by dots, while dashes represent gaps introduced to optimize the alignment consensus. Amino acid substitutions and insertions are indicated by their standard single-letter codes. Notably, an insertion of two amino acids (valine and phenylalanine; VF) at position 386 in the *KAMALA* sequence introduces a structural gap in the reference topology, which consequently disrupts the proper downstream alignment of the GeD3 sequence in this region.

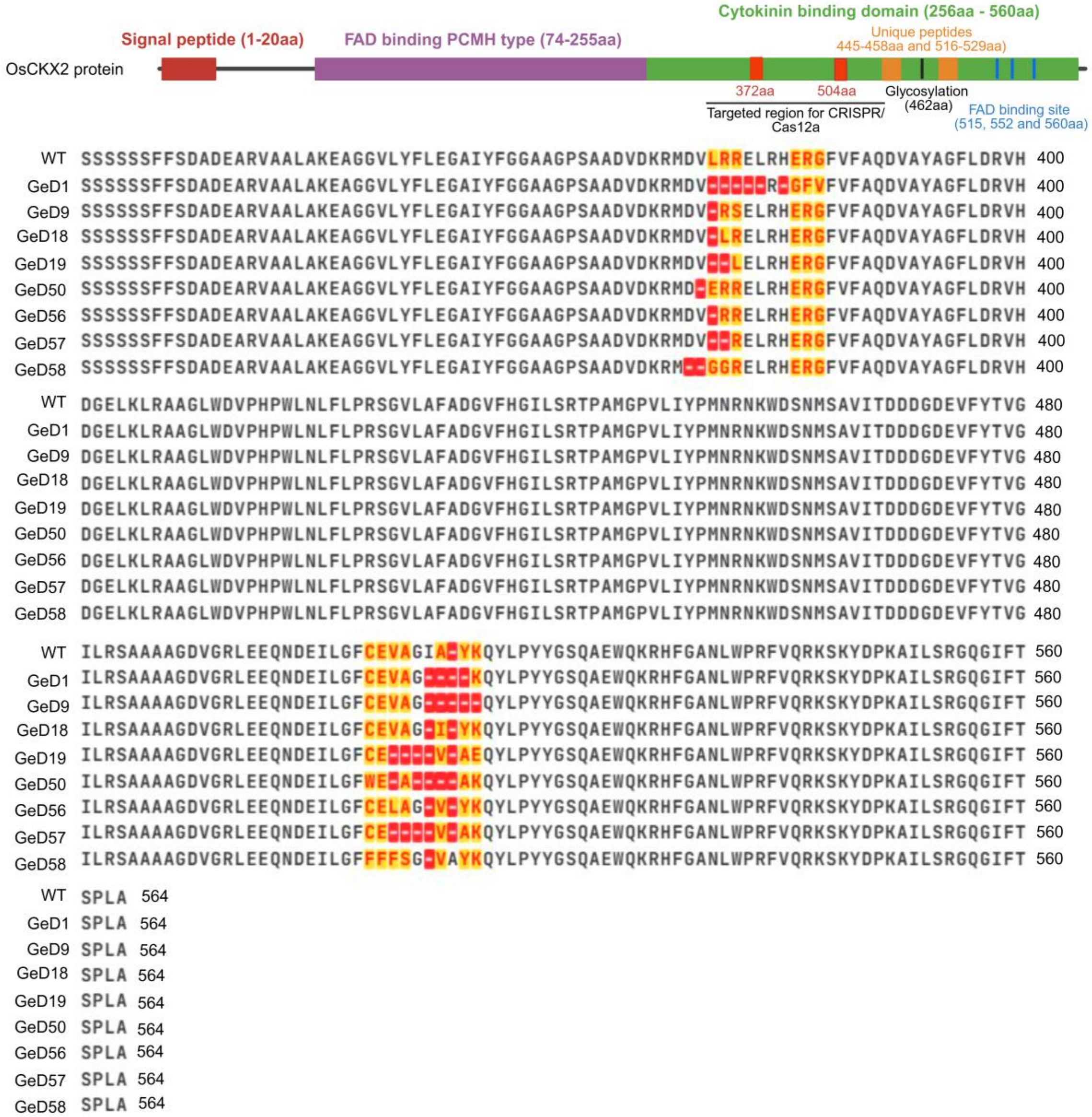


**Supplementary Fig. 31b| Multiple sequence alignment of amino acids**-GeD lines having in-frame deletion in both exon 3 and exon 4 (int Δ E3 + E4) and use the same stop codon as WT Samba Mahsuri

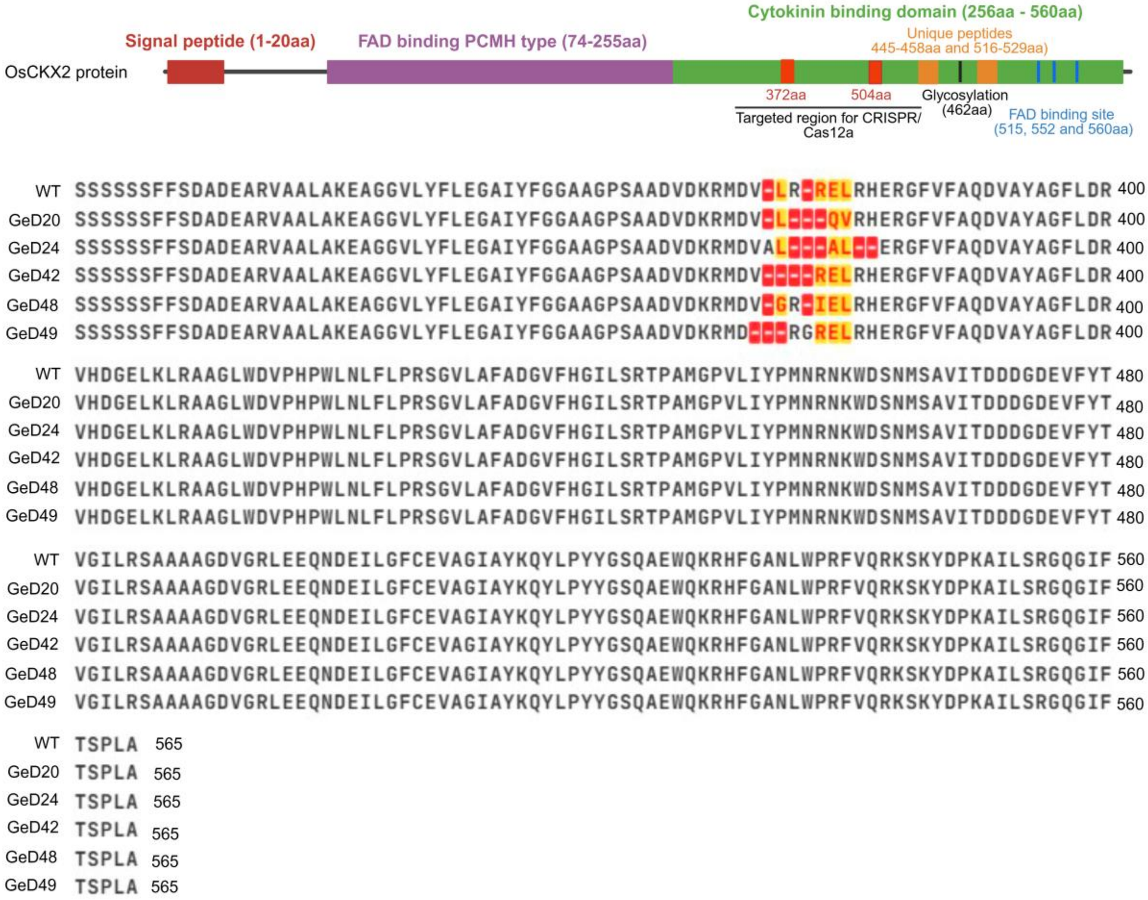


**Supplementary Fig. 31c| Multiple sequence alignment of amino acids**-GeD lines having in-frame deletion in exon 3 (int Δ E3) only and use the same stop codon as WT Samba Mahsuri

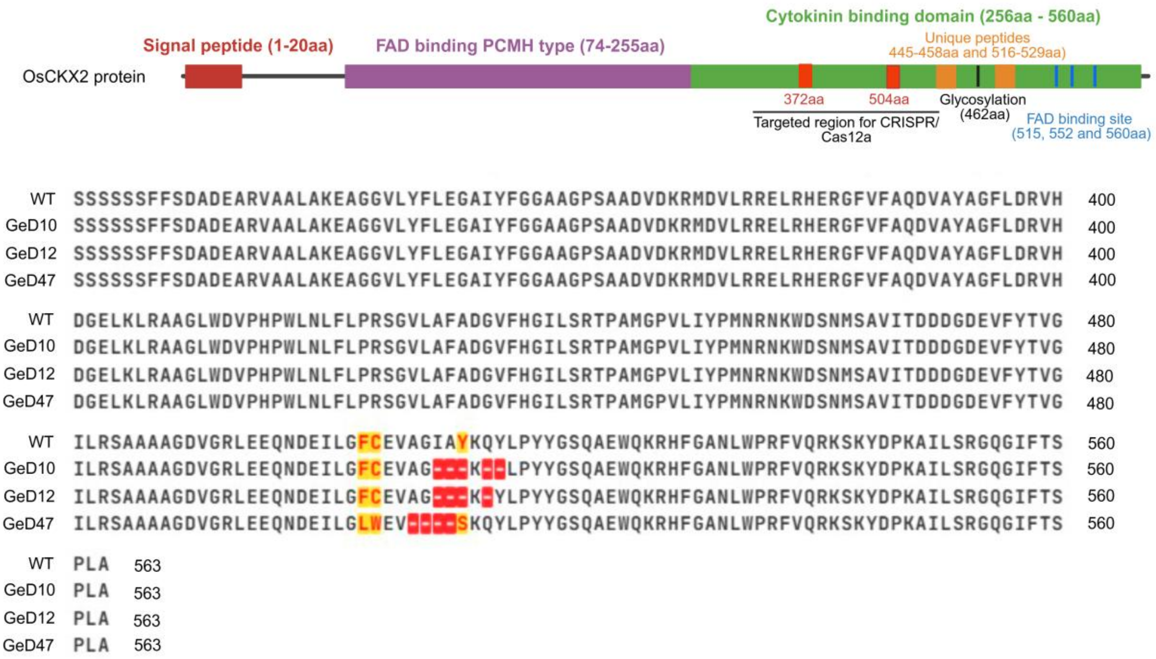


**Supplementary Fig. 31d| Multiple sequence alignment of amino acids**-GeD lines having in-frame deletion in exon 4 only (int Δ E4) and use the same stop codon as WT Samba Mahsuri

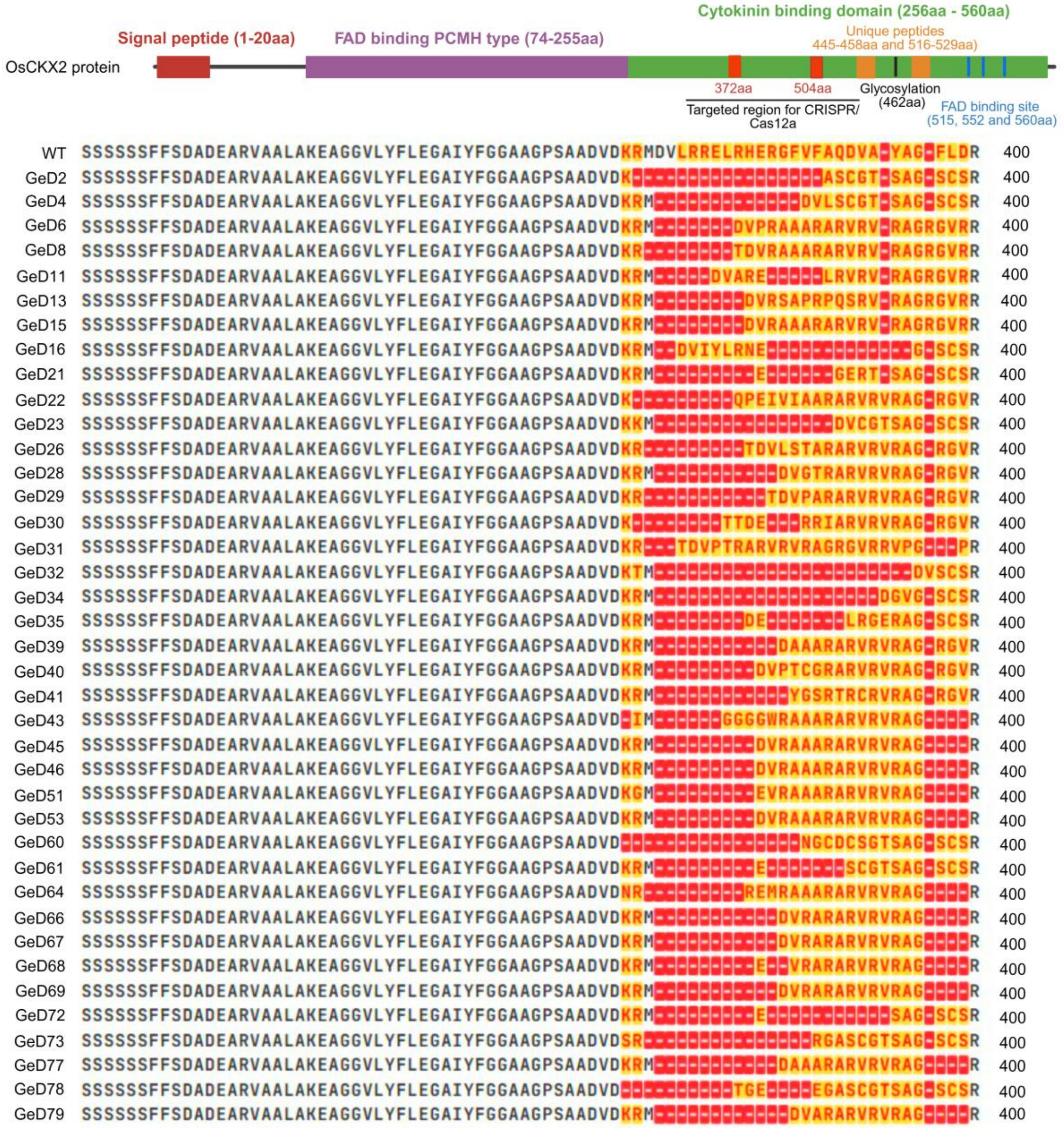


**Supplementary Fig. 31e| Multiple sequence alignment of amino acids**-GeD lines having frameshift mutation in exon 3 causing premature stop codon (Stop E3)

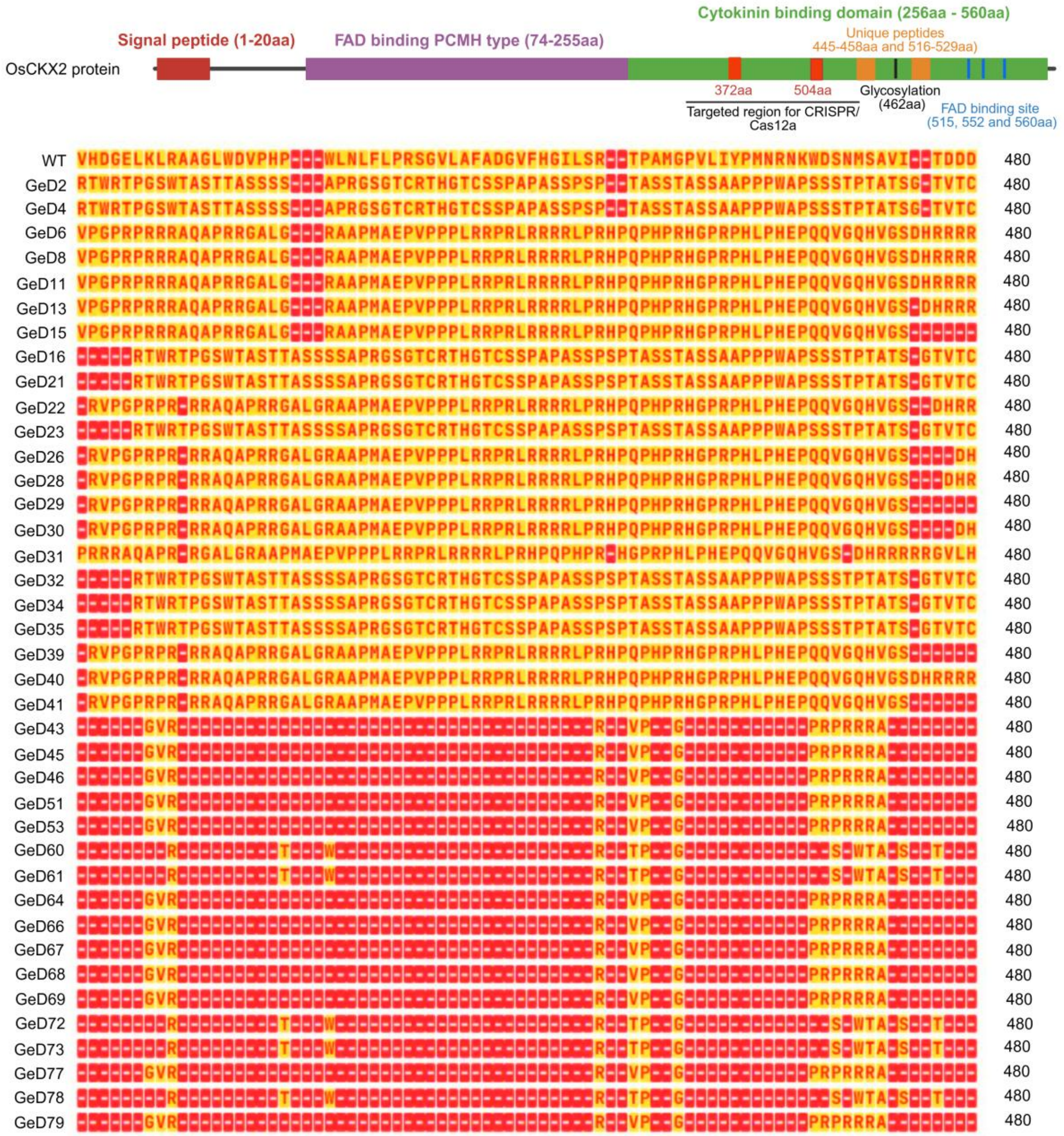


**Supplementary Fig. 31e continue..**

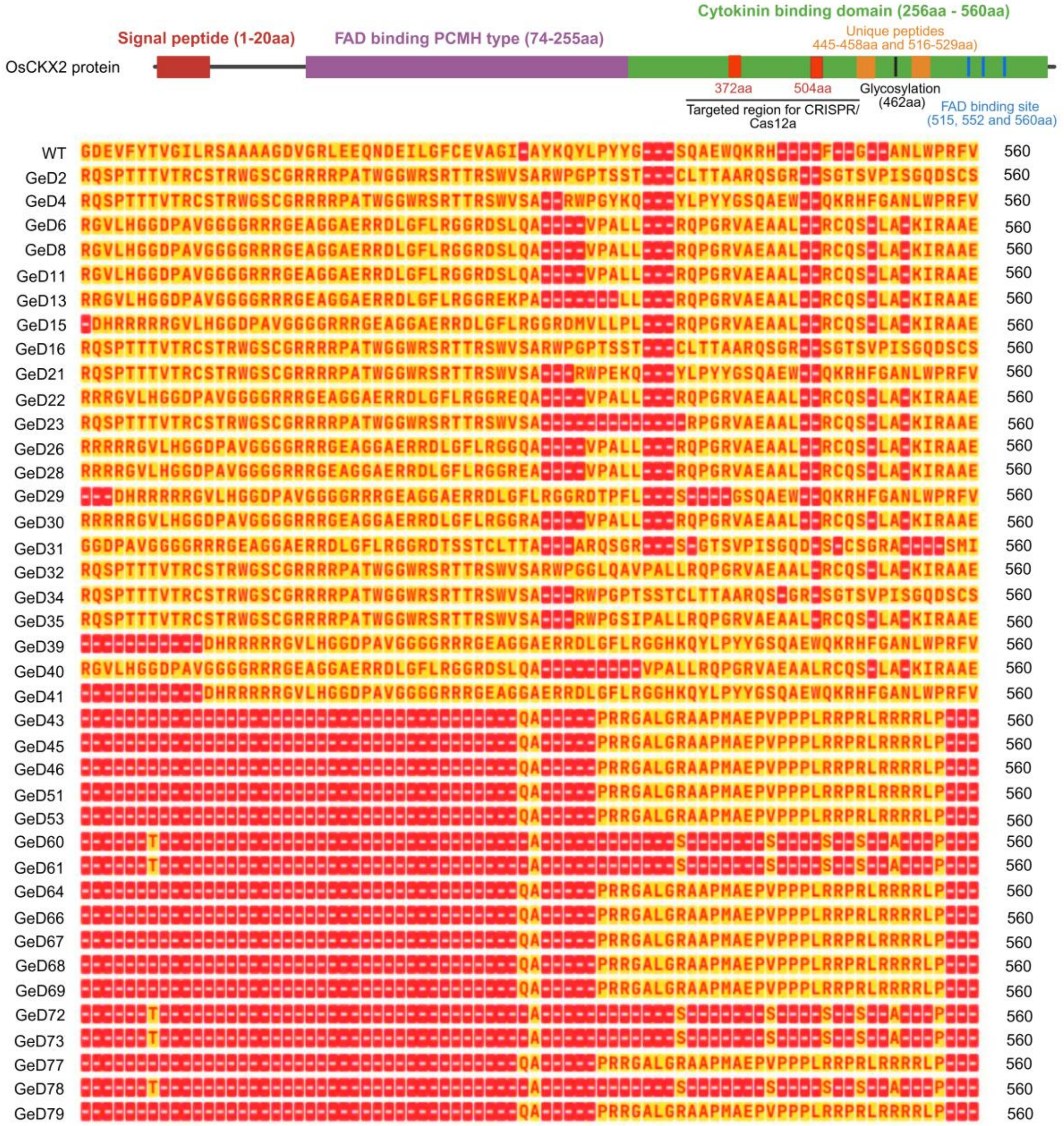


**Supplementary Fig. 31e continue..**

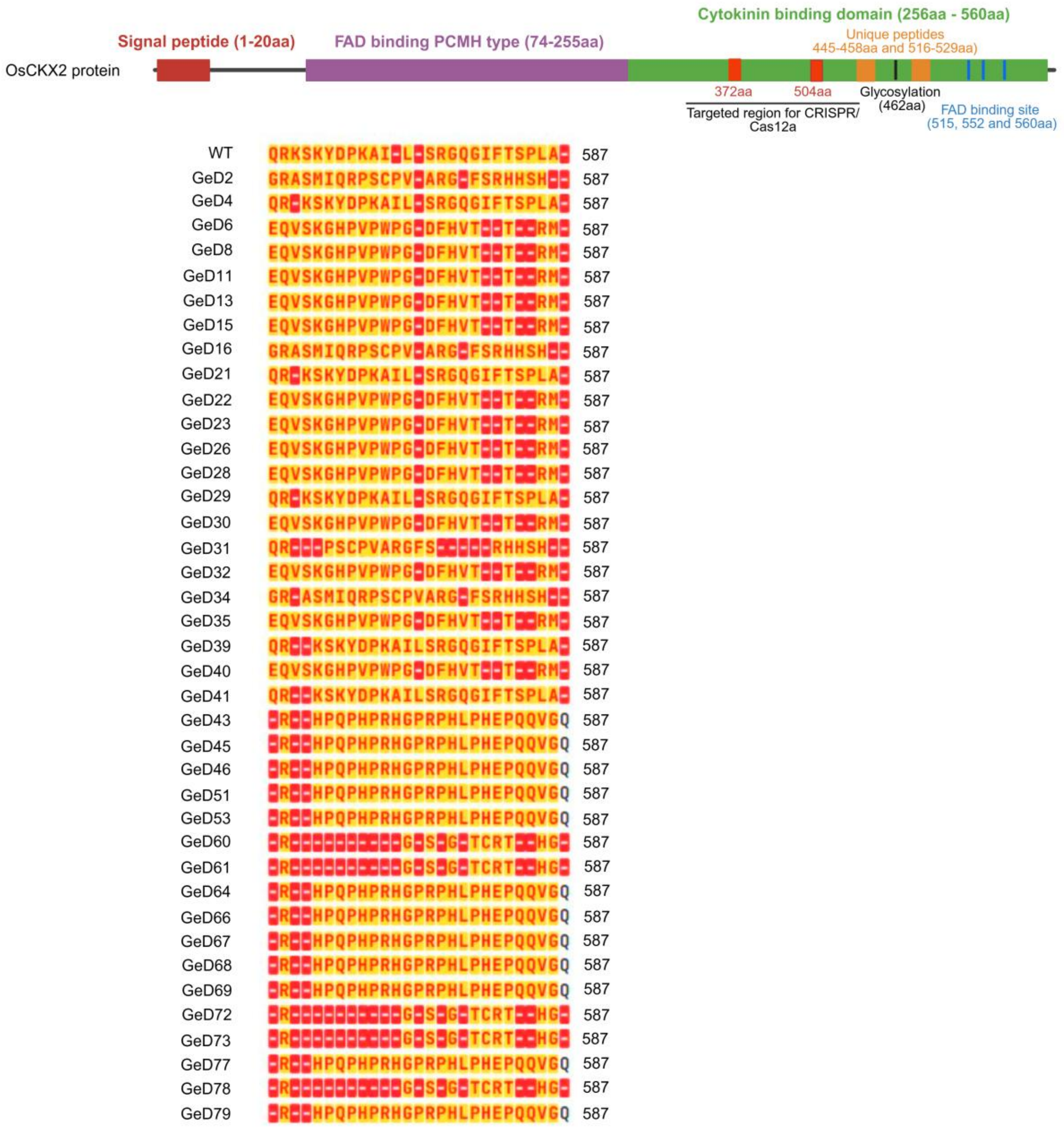


**Supplementary Fig. 31e continue..**

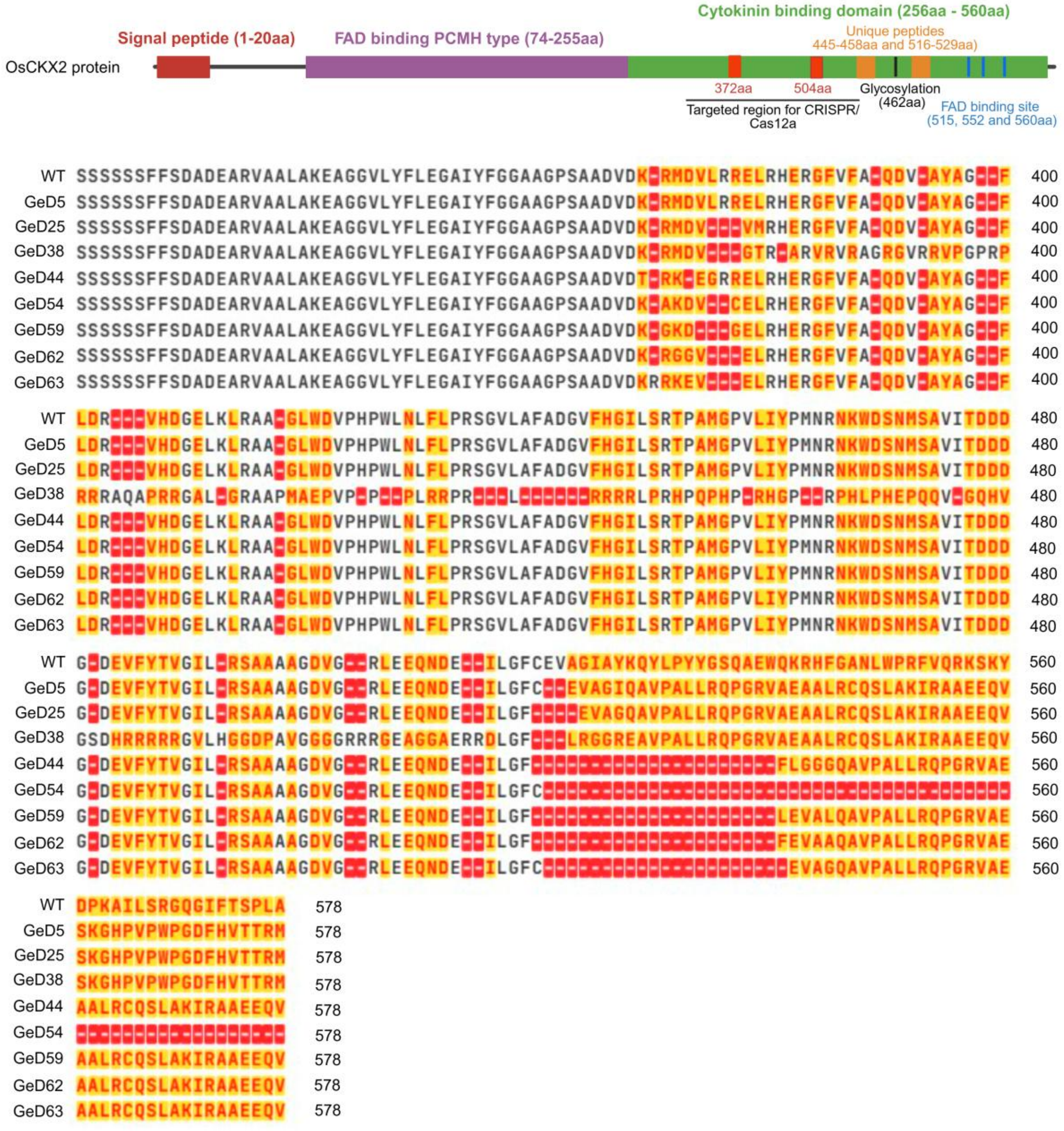


**Supplementary Fig. 31f| Multiple sequence alignment of amino acids**-GeD lines having frameshift mutation in exon 4 causing premature stop codon (Stop E4)

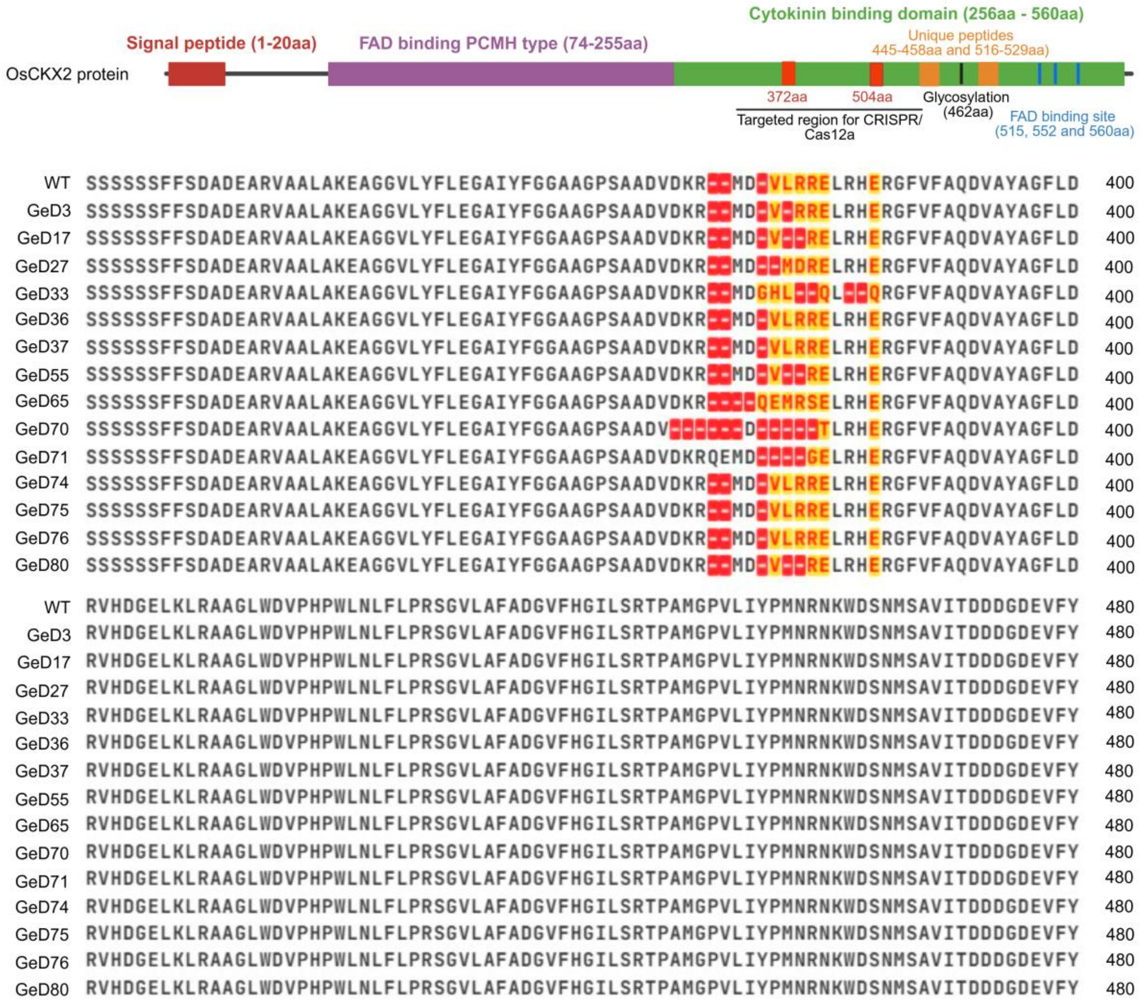


**Supplementary Fig. 31g| Multiple sequence alignment of amino acids**-GeD lines having frameshift mutation in exon 4 causing loss of stop codon

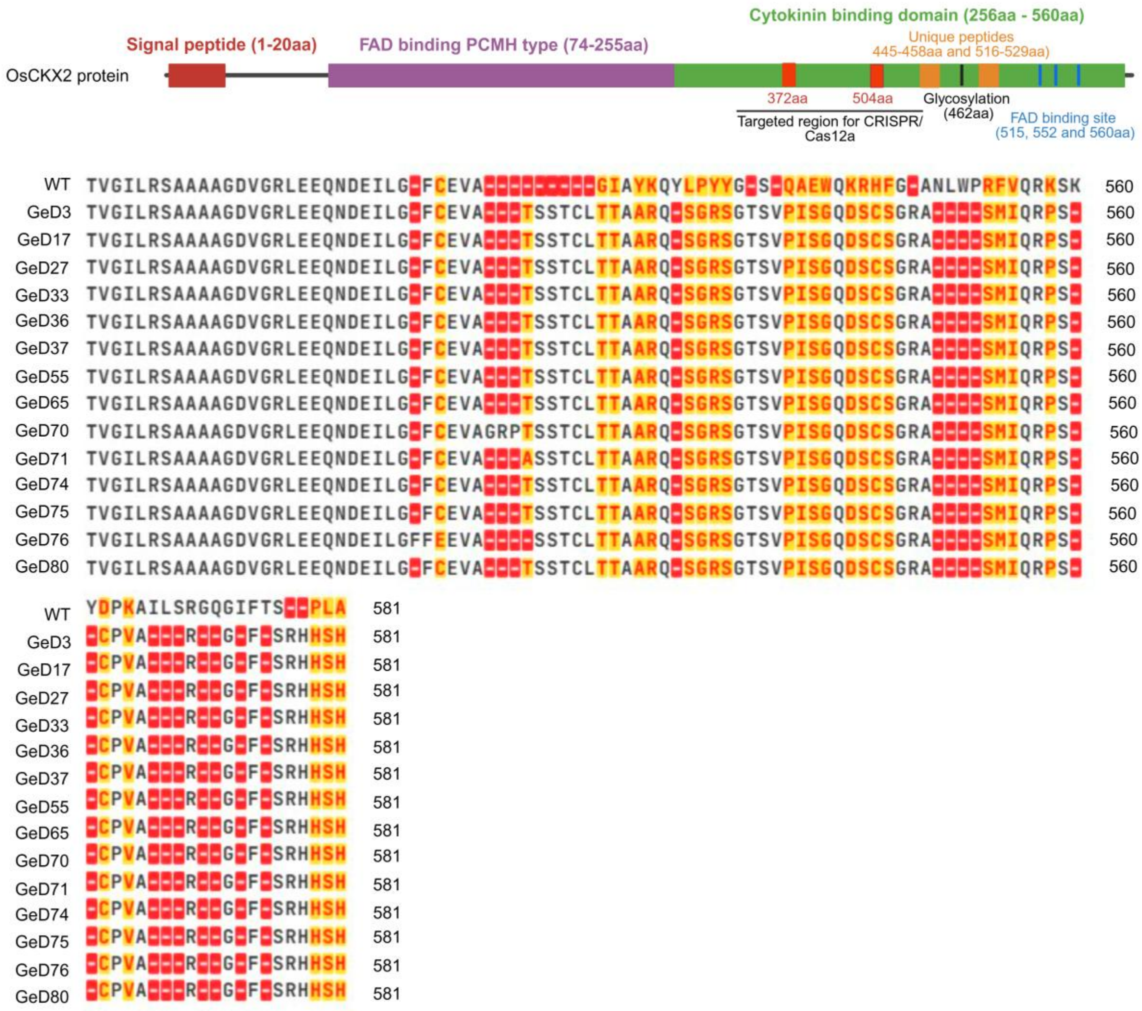


**Supplementary Fig. 31g continue..**

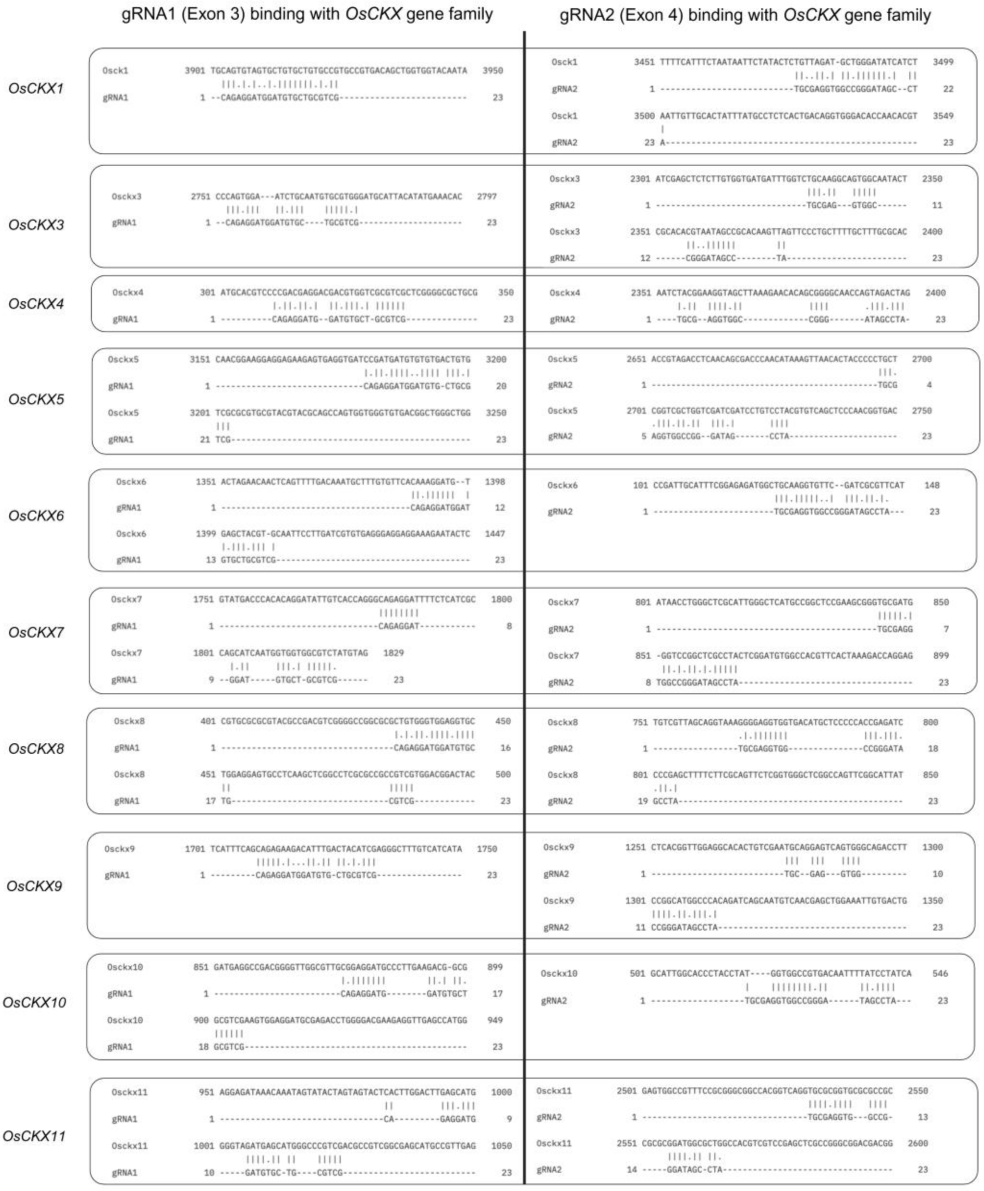


**Supplementary Fig. 32 | *In silico* off-target analysis of *OsCKX2*-targeting sgRNAs across the rice *OsCKX* gene family.** This sequence alignment compares gRNA1 (exon 3) and gRNA2 (exon 4) with other *OsCKX* family members (*OsCKX1, OsCKX3–OsCKX11*). The left panels show gRNA1 binding and off-target sites. The right panels display the corresponding gRNA2 alignments. Vertical bars indicate exact matches between nucleotides. Dashes mark gaps or mismatches nucleotides that do not align between the guide RNA and non-target sequences. Nucleotide positions refer to the reference genome. This analysis demonstrates that the selected sgRNAs for *OsCKX2* are highly specific. All other paralogous family members exhibit significant mismatches and gaps, which minimize the risk of off-target genome editing.
