## Supplementary Notes for "*KAMALA:* improved yield by finetuning cytokinin oxidase activity in rice"

**Supplementary Note-1**

### Regulation of *KAMALA*

- In India, all activities related to Genetically Engineered Organisms or cells are regulated under Rules 1989, notified by the Ministry of Environment, Forest and Climate Change (MoEF&CC). The Rules 1989 fall under the Environment Protection Act, 1986.
- MoEF&CC vide office Memorandum (F.No. C-12013/3/2020-CS-III). dated 30th March 2022, notified exemption of genome edited plants falling under the categories of Site Directed Nuclease-1 (SDN-1) and SDN-2.
- Following this, the Department of Biotechnology, Ministry of Science and Technology, notified 'Guidelines for the Safety Assessment of Genome Edited Plants, 2022 on 17 May 2022. Under these Guidelines, Standard Operating Procedures (SOPs) for Regulatory Review of Genome Edited Plants under SDN-1 and SDN-2 Categories, 2022, were issued on 04$^{th}$ October 2022.
- The SOPs provide a regulatory roadmap and requirements for research, development to meet the threshold for exemption of genome edited plants under the categories of SDN-1 and SDN-2. The SOPs can be accessed at www.dbtindia.nic.in and http://ibkp.dbtindia.gov.in.
- The research, development, and exemption of *KAMALA* (genome edited rice line developed using SDN-1 genome editing) was done strictly as per the guidelines defined by SOPs. The exemption procedure followed for *KAMALA* registration is given below:

### Summary of the step-wise process for regulation of KAMALA

1. Prior to project initiation, an application was submitted to the Institutional Biosafety Committee (IBSC) to initiate genome editing of *CKX2* in the rice variety Samba Mahsuri (February 2018). The application included details of both target gene and crop, guide RNA and vector sequences with annotations, information on genes and promoters in the construct, antibiotic resistance markers, bacterial strains, and biosafety requirements of the project and the laboratory.

2. The application was reviewed and approved by IBSC for submission to the Indian Biosafety Knowledge Portal (IBKP).

3. The Review Committee on Genetic Manipulation (RCGM) examined the application, granted approval for conducting the work under contained conditions, and assigned a Unique Application Code (UAC).

All experimental activities were conducted in contained facilities with regular reporting to IBSC (2018–2023).

**Documentation:** A detailed description of the entire procedure, including DNA sequences, reagents, vectors, and genome-editing tools used; classification of the genome-editing category; Pedigree records of *KAMALA*; identification and evidence of the generation in which homozygous mutations were obtained; records of the backcross generation in which *KAMALA* was confirmed to be free from exogenous DNA (including records of selfing or backcrossing) based on evidences as per the protocol prescribed in the Standard Operating Procedure (SOP)–Section 5; comprehensive phenotypic data of *KAMALA*; assessment of potential unintended phenotypic changes; information on existing identical allele(s), if any; annual compliance reports; and all four annexures as specified in the SOPs.

4. A dossier (including project details, molecular and physiological data, and annexures) was prepared as per SOPs and submitted to IBSC. The documentation part was critical for preparation of dossier.

5. The dossier was evaluated by IBSC, followed by in-person presentation of complete project by investigator which primarily included evidence on mutation characterization and heritability, antibiotic sensitivity, and PCR-based confirmation of absence of exogenous DNA as per SOP–Section 5. Additionally, we did whole genome sequence analysis as described by Huguet-Tapia et al (Nature Biotechnology, 2025, https://doi.org/10.1038/s41587-025-02805-7). Upon satisfactory evaluation, IBSC approved the application.

6. Issuance of the Certificate by IBSC as per the Annexure V was obtained on 14$^{th}$ February 2023.

7. Submission of the Minutes of the IBSC meeting, dossier, certificates along with four annexures and checklists as prescribed in the SOPs to RCGM and IBKP.

8. RCGM endorsed the IBSC decision on 31 May 2023. Accordingly, *KAMALA* was deemed approved for environmental release as per guidelines of the Government of India.

9. Subsequently the evaluation and varietal release process were conducted for *KAMALA* as per established procedures by AICRP (All India Coordinated Research Project). AICRP conducted multi-location field evaluation of *KAMALA* alongside entries developed through various breeding approaches. The seed amount required by AICRP for multi-location testing was 8.0 Kg. *KAMALA* was evaluated across three seasons: the wet season in 2023, the dry season in 2024, and the wet season in 2024.

10. Multi-location data on yield, pest and disease response, and nutrient response were compiled and analyzed. The AICRP data are publicly available at http://aicrip-intranet.in/. Based on the consistent superior performance, KAMALA was presented to the Varietal Identification Committee (VIC) on 26 April 2025 and approved as a "Farmer Variety."

11.The Union Minister of Agriculture & Farmers Welfare formally dedicated *KAMALA* to farmers on 04 May 2025.

12. The final step was submission of the proposal for final release and notification is currently under consideration by the Central Varietal Release Committee (CVRC).

13. In parallel, the Indian Council of Agricultural Research (ICAR) is in the process of securing a Cas12a license for both humanitarian distribution to small and marginal farmers and commercial use. The licensing process with the Broad Institute, USA, is being facilitated by Agrinnovate India Ltd. (AgIn), the commercial arm of ICAR under the Department of Agricultural Research & Education (DARE).

14. Pending successful completion of steps 12 and 13, we anticipate distribution of KAMALA seeds to farmers in wet season of 2026

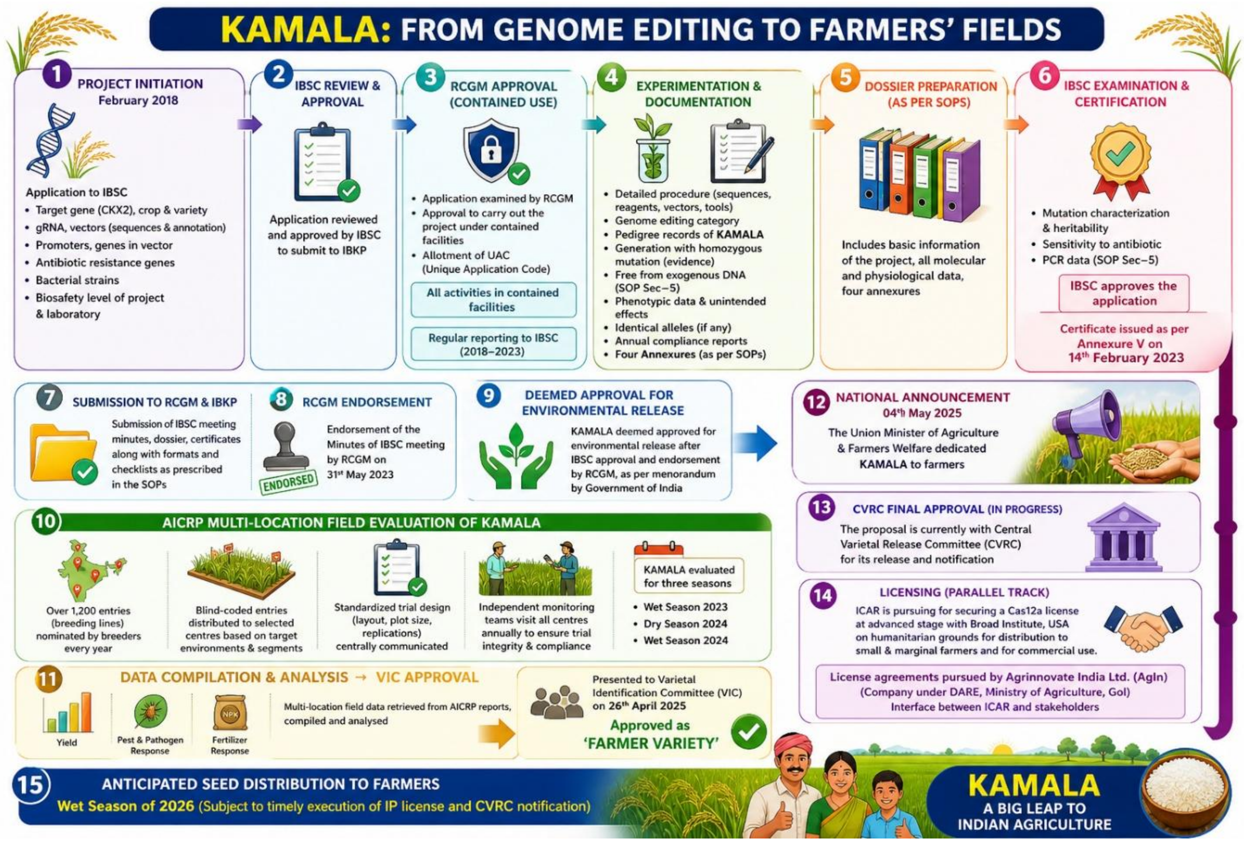


Image: created by ChatGPT

### Annexure-V

#### Format for communicating confirmation of the absence of exogenous introduced DNA from SDN-1 and/or SDN-2 Genome Edited plants by IBSCs

To

The Applicant
Name of the Institute (Affiliation)
Address

The IBSC in its meeting held on ___ has reviewed the application titled _________ for genome edited plant having unique identification number(s)_____under the category of SDN-1/SDN-2 for seeking exemption from Rules, 1989 based on the Office Memorandum F. No. C/12013/3/2020/CS-III dated 30.03.2022 issued by Ministry of Environment, Forest and Climate Change, Government of India.

The details of the genome edited plant(s) are as follows:

**Name of genome edited line(s):**

**Name of the genus and species:**

**SDN-1 or SDN-2:**

**Name/ID of Edited locus:**

**Trait(s), if applicable:**

After a detailed examination of data and methods as specified in the "*Standard Operating Procedures for Regulatory Review of Genome Edited Plants under SDN-1 and SDN-2 categories*", the IBSC confirms the absence of exogenous introduced DNA in the above-mentioned genome edited line(s).

Further development and evaluation of these line(s) are exempted from the Rules, 1989, and will be carried out as per other applicable Laws/Acts/Rules.

***Signature of the Chair, IBSC***

***Copy for information to:-***

1. *Member Secretary, RCGM*
2. *Member Secretary, GEAC*
3. *Food Safety and Standards Authority of India (FSSAI)*
4. *Dept. of Animal Husbandry & Dairying*
5. *Department of Agriculture Research and Education, MoA&FW*

**Supplementary Note-2**

**RNA-seq analysis identified candidate genes associated with improved agronomic traits**

In addition to the qPCR analysis of known specific genes associated with particular phenotype or tissue discussed in relevant section (Supplementary Table 6), we performed transcriptome analysis of inflorescence tissues to identify several other genes that may contribute to the phenotypic differences observed in *KAMALA* relative to WT (Supplementary Figure 19). In *KAMALA*, mRNA levels of 1,254 genes were significantly increased (Supplementary Table 7), while 752 mRNA levels were decreased compared to WT (Supplementary Table 8). Gene Ontology (GO) analysis showed strong enrichment for reproductive development processes, with ~50% of the top GO terms (10-20-fold enrichment) associated with male organ development, gametophyte formation, and floral whorl patterning (Supplementary Fig. 19 and Supplementary Table 7). These results are consistent with the established role of cytokinin as a positive regulator of reproductive development in rice (Yamburenko et al 2017; Tregear et al 2026). Promoter analyses of genes with elevated mRNA levels were used to identify overrepresented binding sites for several transcription factors (TFs) (Supplementary Fig. 19). Notably, OsbHLH068, implicated in inflorescence axillary meristem formation, along with OsAG-group proteins, OsHox16 (HD-Zip I family), and OsbZIP40, emerged as potential regulatory factors. These genes are known to modulate key downstream targets involved in floral organogenesis. To validate the mRNA levels for some of the genes identified here, RT-qPCR was performed on RNA from flag leaf and inflorescence tissues and the results corroborated the RNA-seq findings (Fig. 4k). The *Os02g0490000* (*OsPUB73*), a crucial component of the regulation of male reproductive development involved in tapetum and pollen exine formation (Chen et al 2019), exhibited significantly higher mRNA levels in *KAMALA* relative to WT. In addition, differential level of *OsGA2ox10, OsGA20ox3*, and *OsANK1* transcripts was observed between *KAMALA* and WT (Fig. 4k).

### Supplementary Note-3

#### Key Confidence Metrics of AlphaFold 3 predicted structures of CKX2 protein in WT and GeD lines

| | WT | WT (Samba Mahsuri) | GeD1 (KAMALA) |
|---|---|---|---|
| **pLDDT** | > 90 [Except N-terminal and few loop Regions] fraction_disordered: 0.05 | > 90 [Except N-terminal and few loop Regions] fraction_disordered": 0.06 | > 90 [Except N-terminal and few loop Regions] fraction_disordered: 0.06 |
| **PAE** | 0.76 | 0.76 | 0.76 |
| **pTM** | 0.90 | 0.90 | 0.89 |
| **ipTM** | 0.90 | 0.90 | 0.89 |

| | GeD 2 | GeD 3 | GeD 4 |
|---|---|---|---|
| **pLDDT** | >90 for core regions and <50 for disordered regions fraction_disordered: 0.39 | >90 for core regions and <50 for disordered regions fraction_disordered: 0.13 | >90 for core regions and <50 for disordered regions fraction_disordered: 0.22 |
| **PAE** | 0.76 | 0.76 | 0.76 |
| **pTM** | 0.62 | 0.83 | 0.75 |
| **ipTM** | 0.62 | 0.83 | 0.75 |

**pLDDT (Per-residue Local Distance Difference Test):** Assesses local confidence. Scores above 90 represent high backbone and side-chain accuracy, while scores below 50 indicate potential disordered or low-confidence regions.

**PAE (Predicted Aligned Error):** Measures the expected position error between any two tokens (not just amino acids) in the structure. Lower PAE values indicate better confidence in the relative orientation of different domains.

**pTM (Predicted Template Modeling):** Evaluates the overall folding topology and accuracy of the complex. Scores above 0.5 are generally confident.

**ipTM (Interface Predicted Template Modeling):** Specifically measures the accuracy of the relative positions of subunits in complexes. Scores higher than 0.8 represent confident, high-quality inter-chain predictions.

**Supplementary Note-4**

### Tissue staining and microscopy

The 60-day old plant root tissue (1 cm below to shoot junction), mid-leaf and stem ($1^{st}$ internode) were collected for staining in the transverse panel. Samples were stored in 10% formaldehyde, followed by paraffin block preparation for a 10 µm thick section through a microtome (Leica RM2125 RTS). Sections were stained with H&E (hematoxylin and eosin)(Bancroft and Layton 2013), and the slide was scanned in a NanoZoomer-SQ Digital slide scanner (Hamamatsu Photonics K.K., Japan). The visual comparison of leaf, root, and grain were captured on the Leica M205 FA microscope (Germany).

The shoot apical region of 8-week-old field-grown plants was dissected and incubated in ½ MS medium containing 10 µM EdU for 1 h under 200 $\mu mol\ m^{-2}\ s^{-1}$ light at 28 °C. Following incubation, shoot apices were washed and transferred to fresh ½ MS for 4 h before being fixed in 4% paraformaldehyde (PFA) in PBS, for 16 h at 4 °C. Fixed samples were washed again in PBS, dehydrated through an ethanol gradient, and embedded in paraffin. Sections of 10 µm thickness were cut, dewaxed, and rehydrated through a reverse ethanol gradient. The sections were then permeabilized and processed with the Click-iT reaction cocktail (Click-iT EdU Cell Proliferation Kit) following the manufacturer's instructions. For anatomical visualization, Calcofluor White was used as a counterstain. EdU-labeled cells were imaged using a Leica TCS SP8 confocal laser scanning microscope with Alexa Fluor 488 settings, using a 10× objective, Argon laser at 5% intensity, and PMT detector set to 780% gain, with excitation at 488 nm and emission collected between 500–540 nm.

### RNA isolation and RT-qPCR

Tissues at various growth stages, including early seedlings leaf (21-day-old plants), inflorescences, and flag leaves, were collected in 2 mL Eppendorf tubes, immediately frozen in liquid nitrogen, and stored at -80 °C. RNA isolation was carried out using TRIzol Reagent (Invitrogen) as per the standard protocol(Solanki et al. 2019). Reverse transcription was done using PrimeScript $1^{st}$ strand cDNA Synthesis Kit (Takara). Approximately 1 µg total RNA was used per reaction. Oligo(dT) primer was used to prepare cDNA from mRNAs. The RT reaction was carried out at 42 °C for 60 min, followed by inactivation at 70 °C for 15 min. To measure transcript levels by RT-qPCR, a Bio-Red 96CFX Real-time PCR machine was used with Luna Universal qPCR Master Mix (New England Biolabs). Three biological replicates and two technical replicates were performed for each assay. The program for thermal cycling was as follows: Step 1: 1 cycle, 95 ˚C for 3 min, Step 2: 40 cycles, 10 sec at 95 °C, 30 sec at 60 °C. The melt curve analysis was performed with 5 sec at 65 °C and 5 sec at 95 °C, with an increment of 0.5 °C. The actin gene was used as the internal reference gene. Relative mRNA levels were calculated using a $2^{-}\Delta\Delta$ct methodology (Solanki, et al. 2019). The heat map plot included relative mRNA values in the figures for presentation.

**RNA sequencing and analysis**

For RNA sequencing, samples were collected at the developmental stage of inflorescence primordia from WT and *KAMALA*. Following the paired end (2x150 bp) sequencing on the Illumina NovaSeq 6000 platform, Fastp software was employed to eliminate adapter sequences and filter out low-quality reads. Trimmomatic and HISAT2 were utilized to trim the reads and align them to the reference genome (Os-Nipponbare-Reference-IRGSP-1.0). The normalized read count data were subsequently analyzed to calculate the |log2fold change| value using DESeq2 applying stringent criteria of *P* value <0.05 and |log2fold change| > 2 to identify differentially expressed genes (DEGs). Enrichment analysis of these DEGs was conducted using iDEP and ShinyGO software (https://bioinformatics.sdstate.edu/go/; https://bioinformatics.sdstate.edu/idep/). The Rice Annotation Project Database (RAP-DB) was accessed to retrieve the upstream 1kb sequences of all the up regulated candidate genes. Concurrently, transcription factor motifs were obtained from JASPAR database (https://jaspar.elixir.no/). All these retrieved promoter sequences and TF motifs were served as input to MEME suite-FIMO to ascertain target specific- transcriptional factor binding motifs. The output generated from FIMO facilitated the construction of a network illustrating the interactions between transcription factor binding sites and their corresponding genes employing Cytoscape 3.10.4 (https://cytoscape.org).

**Measurement of cytokinin oxidase activity**

The CKX enzymatic activity was quantified at the inflorescence developmental stage corresponding to approximately 5 cm in length, as described earlier (Joshi et al. 2018; Libreros-Minotta and Tipton 1995; Powell et al. 2013; Tsago et al. 2020). Briefly, 500 mg fresh-weight frozen tissue from WT and *KAMALA* plants was homogenized and prepared as a slurry by mixing 2 mL chilled buffer (50 mM potassium acetate, 1 mM $MgSO_4.7H_2O$, 2 mM $CaCl_2.2H_2O$ and 0.5 mM DTT). Subsequently, 7.5 µL protease inhibitor cocktail (Himedia, ML051) was added to the homogenate. It was centrifuged at 15871 × g for 20 min at 4 °C, the supernatant was collected and mixed with 0.015 volume of an aqueous solution comprising 5%

(v/v) polyethyleneimine, 0.5 mM phenylmethanesulfonyl fluoride and 0.5 mM Nα-tosyl-L-lysine chloromethylketone (Sigma-Aldrich, T7254). This mix was centrifuged at 15871 × g for 20 min at 4 °C, and the sediment was discarded. A total of 200 μL supernatant was added to 400 μL reaction mixture containing 75 mM Tris-Cl pH 8.5, 0.5 mM Dichlorophenolindophenol (Himedia, RM350) and 0.15 mM N6-isopentenyl adenine (iP) (Himedia, PCT0807). The reaction mixture was placed in a water bath at 37 °C for 1 h. To stop the reaction, 300 μL 40% trichloroacetic acid (Sigma-Aldrich, A71328) was added and mixed by vertexing. The mixture was centrifuged at 13523 × g for 5 min at 4 °C. A total of 200 μL 3% 4-aminophenol prepared in 6% trichloroacetic acid (Amresco) was added to the supernatant and incubated at room temperature for 10 min. Absorbance was measured at 352 nm using a UV-visible spectrophotometer (Eppendorf, Germany). Total protein was quantified using Bradford reagent (Himedia, MB092). The specific activity of CKX was expressed as units $mg^{-1}$ protein.

**Phytohormone extraction and profiling**

Approximately 5 cm grown inflorescence tissue and 2 cm primary root tissue from the shoot-root junction at the vegetative stage (45-day-old plants) were collected and immediately frozen in the liquid nitrogen, and stored at -80 °C. A total of 100 mg homogenized tissue powder was mixed with 500 μL pre-chilled extraction solvent consisting of 2-propanol: MilliQ water: concentrated HCl in a ratio of 2:1:0.002 (v/v/v). After mixing, extraction was carried out by shaking the sample at 100 rpm for 30 min at 4 °C in a thermomixer. Thereafter, 1 mL of dichloromethane (DCM) was added to the sample mix, and the incubation continued for 30 min at 4 °C with shaking at 100 rpm. The sample was centrifuged at 13,000 × g for 10 min at 4 °C. The lower phase (~900 μL) was carefully collected, transferred to a fresh 1.5 mL Eppendorf tube, and dried completely using a SpeedVacμL (Thermo Scientific, USA). The dried residue was dissolved in 100 μL of pre-cooled 100% methanol followed by centrifugation at 10,000 × g for 2 min. The sample solvent was transferred to an injection vial, and 7.5 μL was injected into reverse phase Hypercil GOLD C18 column (2.1×75 mm, 2.7 m) (Thermo Scientific) for Ultra Performance Liquid Chromatography (UPLC) [Aquity UPLCTM System, a quaternary pump, and an autosampler (Waters, Milford, MA, USA)] /ESI-MS [Exactive-Plus Orbitrap mass spectrometer (Thermo Fisher Scientific, USA)] analysis. Phytohormone profiling was performed at the Repository of Tomato Genomics Resources (RTGR), University of Hyderabad (https://rtgr.uohyd.ac.in/) and Novelgene Technologies (https://novelgenetech.com/).

**Supplementary Note-5**
